# Sorghum embryos undergoing B chromosome elimination express B-variants of mitotic-related genes

**DOI:** 10.1101/2025.09.30.679447

**Authors:** T Bojdová, L Hloušková, K Holušová, R Svačina, E Hřibová, I Ilíková, J Thiel, G Kim, R Pleskot, A Houben, J Bartoš, M Karafiátová

## Abstract

**Background:** Selective DNA elimination occurs across diverse species and plays a crucial role in evolution and development. This process encompasses small deletions, complete removal of chromosomes, or even the elimination of entire parental genomes. Despite its importance, the molecular mechanisms governing selective DNA elimination remain poorly understood. Our study focused on the tissue-specific elimination of *Sorghum purpureosericeum* B chromosome(s) during embryo development.

**Results:** In situ B chromosome visualisation, complemented by transcriptomic profiling and gene-enrichment analysis, allowed us to identify 28 candidate genes poten-tially linked to chromosome elimination. We show that elimination is a developmentally programmed process, peaking during mid-embryogenesis and nearly completed at later stages, leaving B chromosomes only in restricted meristematic regions. Genome sequencing revealed that the sorghum B chromosome is of multi-A chromosomal origin, has reduced gene density, is enriched in repetitive sequences, and carries a novel centromeric repeat (SpuCL166). Transcriptome analyses identified B-specific variants of kinetochore, cohesion, and checkpoint genes that are expressed during active elimination, while structural modeling of CENH3 and CENP-C indicated functional divergence at the kinetochore interface.

**Conclusion:** Here, we provide the first comprehensive genomic and transcriptomic characterization of B chromosome and its elimination in *Sorghum purpureosericeum*. Our findings suggest that B chromosomes express modified mitotic machinery to control their own fate. By establishing a framework of candidate genes, this study opens new avenues for dissecting the molecular mechanisms of chromosome elimination and provides a critical foundation for understanding how genomes evolve to regulate and tolerate supernumerary chromosomal elements.

## Background

DNA elimination represents a dramatic form of DNA silencing that has been studied for more than a century [1]. It plays a pivotal role in evolution and development, contributing to speciation, adaptation, and sex determination in various species [2, 3, 4, 5]. Selective DNA elimination can involve the deletion of defined DNA sequences, complete removal of entire chromosomes or subgenomes [6] or even the elimination of part or entire parental genome during interspecific hybridization [7, 8]. Further, sex determination based on selective DNA elimination has been documented in *Sciara* [9] and other insects, such as the parasitoid wasp *Nasonia vitripennis* [10].

Supernumerary B chromosomes represent another intriguing case of selective DNA elimination as in some species, B chromosomes are eliminated in a tissue-specific manner [11, 12, 13, 14, 15]. While plants represent the majority of about 3,000 species in which B chromosomes have been identified, tissue-specific elimination of B chromosomes is rarely documented in plants [16]. In grasses *Poa alpina* and *Agropyron cristatum,* B chromosomes are eliminated from adventitious roots but retained in primary roots [13], whereas in *Aegilops speltoides, Aegilops mutica* and *Poa timoleontis*, B chromosomes are eliminated from primary roots [12, 14, 17]. A particularly striking example of selective DNA elimination is documented in wild sorghum, *Sorghum purpureosericeum,* which shows a near-complete B chromosome elimination in mature somatic tissues [18], resembling the model of germline-specific chromosomes (GRC) in songbirds, which are eliminated from all somatic cells [19, 20]. This wide phylogenetic distribution of selectively eliminated DNA suggests that the mechanisms behind selective DNA elimination evolved independently across species [4].

Despite the increasing interest in the topic, the mechanisms underlying selective chromosome elimination remain unclear. A key challenge in this field is to understand which cellular processes are responsible for the selective identification of particular chromosomes and DNA sequences for elimination. The mechanism of chromosome recognition has been best analysed in unicellular ciliates, where specific DNA sequences are targeted for elimination through short non-coding RNAs (sncRNA) [21, 22, 23, 24]. However, sncRNAs’ role in DNA elimination has never been demonstrated in multicellular organisms, and no alternative mechanism of targeted chromosome recognition has been described in any species.

In animals, the mechanism of selective elimination of DNA is often associated with abnormal kinetochores and specific epigenetic modifications of histones [25, 26, 27]. For B chromosomes, elimination is associated with the formation of micronuclei, caused by faulty chromosome segregation [3, 28]. As a result, the chromosomes remain at the metaphase plate, form micronuclei, and undergo degradation. Indeed, this mechanism of B chromosome elimination has been observed in the embryos of sorghum *S. purpureosericeum* and *Aegilops speltoides* – the two plant species recently studied for B chromosome elimination [15, 18, 29, 30]. While it is proposed that the elimination process is controlled by B chromosome-expressed genes, the genes involved remain unidentified. A pilot study aiming to identify genes involved in the process of B chromosome elimination in *Ae. speltoides* revealed the B-specific variant of the kinetochore gene *NUF2* and additional candidates, which require further investigation [29].

This study aims to take the first step towards the understanding the B chromosome elimination during embryonic development in wild sorghum *S. purpureosericeum*. With this in focus, we created and analyzed the first genome assembly at the contig level, annotated A and B chromosome-encoded genes and analyzed the transcriptome in the developing embryo. Sequence analysis revealed a B chromosome-specific (peri)centromere composition and the multi-A chromosome origin of the B chromosome. Differential expression analysis uncovered a set of B chromosome-encoded genes associated with chromosome segregation. For *in silico* protein modelling, we focused on the kinetochore proteins acting in proximity of centromeric DNA and identified significant polymorphisms at the their interacting interface.

## Results

### Embryo differentiation is accompanied by extensive B chromosome elimination

Our preliminary data revealed that B chromosome elimination occurs extensively during embryo development [18]. In this context, we performed experiments on the embryos of wild sorghum in three developmental stages (Fig. 1) characterized by progressing organogenesis (Fig. 1B1–D1). To describe the dynamics of the elimination process, we used FISH with a B-specific probe on longitudinal cryo-sections of the embryos in the three developmental stages.

**Fig. 1:**
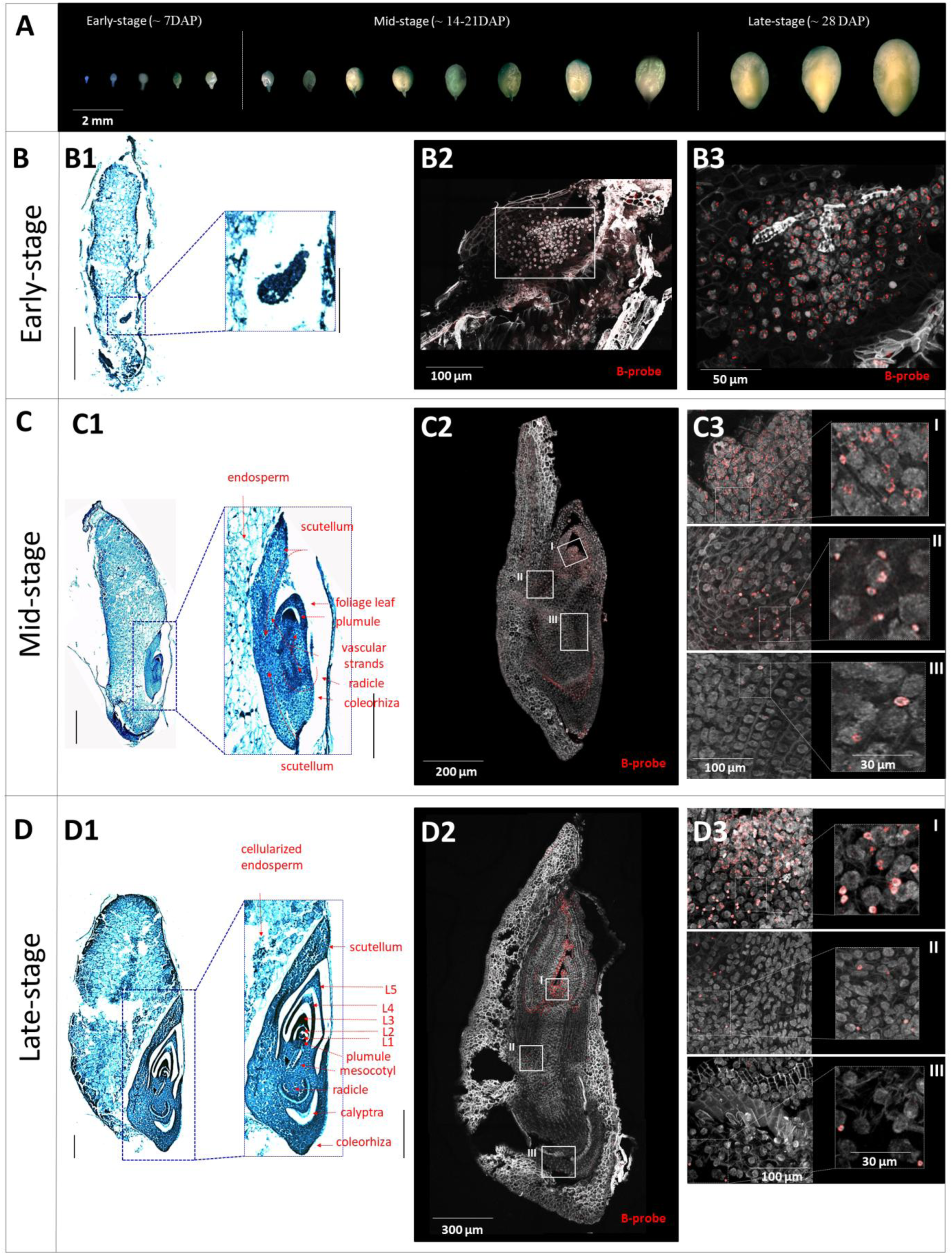
Progression of the B chromosome elimination in the context of embryonal development visualized using fluorescent *in situ* hybridization with B-specific probe. **A** Scaling of embryo growth in wild sorghum and categorization of the embryos **B-D** Progress of B chromosome elimination in early **(B)**, mid **(C)** and late stages of embryo development **(D)**. **B1-D1** Size and anatomy of the embryos in targeted stages on toluidine blue-stained sections; whole-seed section (left) scale bar: 0.5 mm; embryo (right) scale bar: 0.3 mm. **B2-D2** B chromosome distribution visualized using FISH with B-specific probe on embryo cryo-sections. White-framed regions were further used to investigate the micronucleation of the B chromosome. **B3-D3** Magnified white-framed regions from B2-D2 of SAM (I), mesocotyl (II) and root (III) (on the left) and detail of micronuclei (on the right). No micronuclei were detected in the early-stage **(B3).** Tissue of plumule and foliage leaves in mid-stage **(C3-I)** and in late-stage **(D3-I)** show a mosaic pattern of micronuclei and +B-nuclei. Mesocotyl of mid-stage embryos is micronuclei rich **(C3-II)**; while their number in late stage is strongly reduced **(C3-II)**. In both developing **(C3-III)** and developed **(D3-III)** radicles, only residual micronuclei are observed.

The early-stage embryos revealed a uniform distribution of B chromosomes in most cells (Fig. 1B2,3, Additional file 1: Fig. S1), which is consistent with the observations in *Ae. speltoides* [15]. B chromosome-specific signals were uniformly observed within the nuclei. Cells lacking B chromosomes or containing micronuclei were rare at this stage. Thus, a negligible level of B chromosome elimination happens in undifferentiated cells before the establishment of embryonic organs.

In mid-stage embryos, extensive B chromosome elimination becomes frequent (Fig. 1C2, Additional file 1: Fig. S1). Embryo development progressed towards organogenesis, characterized by the visible formation of the radicle, plumule and leaf primordia. B chromosomes have already been excluded from the nuclei of the radicle and mesocotyl, and even the micronuclei have mostly been degraded. Simultaneously, the whole aerial part exhibited B-specific signals, and the B chromosome-positive cells showed a mosaic pattern. Except for the zone of plumule, where the B chromosomes constantly occurred in all nuclei, the cells of the upper mesocotyl and embryonic leaves were affected by the elimination of B chromosomes and showed a micronuclei-enriched pattern (Fig. 1C3 I-III).

In late-stage embryos, the occurrence of the B chromosome elimination dropped dramatically, because very few cells preserving B chromosome remain. The B chromosome distribution pattern contrasted strongly with previous stages and reached nearly its final shape (Fig. 1D2, Additional file 1: Fig. S1). B chromosomes remained in limited meristematic regions (SAM and axillary buds) and in an unspecified cell lineage bordering the marginal layer of the youngest foliage leaf. Surprisingly, the cells in these regions preserve the ability to eliminate B chromosomes as the micronuclei were still observed (Fig. 1D3 I-III).

Our FISH experiments uncovered the dynamics of the B chromosome elimination in developing embryos and showed the ambiguous association of this process with cell differentiation. Interestingly, the process of B chromosome elimination does not stop with the maturity of embryos but continues through the plant growth [18].

### The B chromosome of S. purpureosericeum is of multi-A chromosomal origin and carries a centromeric B-specific repeat

To generate the first genomic sequence of *S. purpureosericeum*, we sequenced the DNA of plants harbouring the B chromosome (+B). We combined the sequences derived from Illumina and Oxford Nanopore Technology (ONT) platforms. 127 gigabases (Gb) of long ONT reads were employed in the *de novo* assembly of the *S. purpureosericeum* genome utilizing the SMARTdenovo assembler, followed by a polishing series with long and short reads (Fig. 2A). The draft assembly yielded a sequence of 2.81 Gb in length, comprising 19,520 contigs and N50 321.01 kb (Table 1). This result exceeds by 7% the previously estimated genome size of 2.21 Gb per haploid genome (1C) combined with the B chromosome size of 421 Mb as reported by Karafiátová et al. [30]. The BUSCO analysis within the Poales lineage yielded a completeness score of 98.7% (Fig. 2B). K-mer analysis was used to compare the frequency of 39-mers derived from 0B and +B Illumina reads along diagnostic (low-copy) regions of each contig in the assembly. A total of 4,437 contigs were identified as B chromosome contigs, collectively spanning 420 Mbp (Fig. 2C, Additional File 2). Of these, 212.9 Mbp were classified with high confidence, while 207 Mbp were identified with low confidence. Annotation implementing braker3 pipeline revealed 56,920 genes with 97.7% BUSCO completeness (Fig. 2B). Among these annotated genes, 4,490 were identified as being located on the 4,437 B chromosome contigs (Additional file 3).

**Fig. 2:**
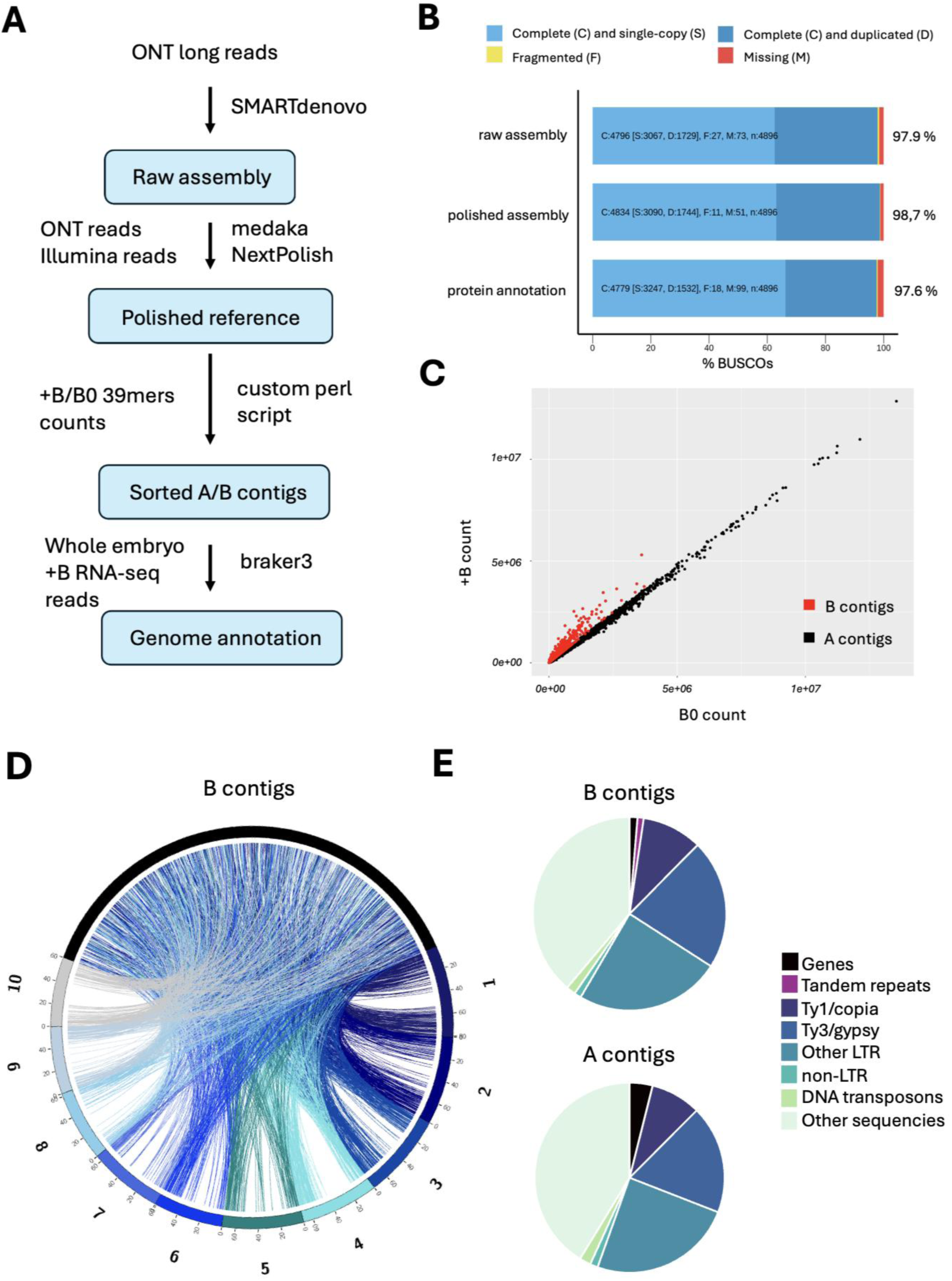
Sequence assembly and annotation of the *S. purpureosericeum* B chromosome. **A** Schematic workflow of genome assembly, sorting of contigs and genome annotation. **B** Results of BUSCO analysis after raw assembly, polishing process and for annotated genes. **C** Scatter plot of counts of 39-mers for +B vs. 0B sample. Each dot represents a contig in the assembly; B contigs are highlighted in red. **D** Circos plot (not in scale) depicting syntenic relationship between 10 *S. bicolor* chromosomes (1-10; shades of blue) and *S. purpureosericeum* B-contigs (black). Each ribbon corresponds to a homolog pair. **E** Sequence content of *S. purpureosericeum* A chromosomal complement (all A-contigs; 2.4 Gb) and the B chromosome (B-contigs; 420 Mb).

**Table 1:**
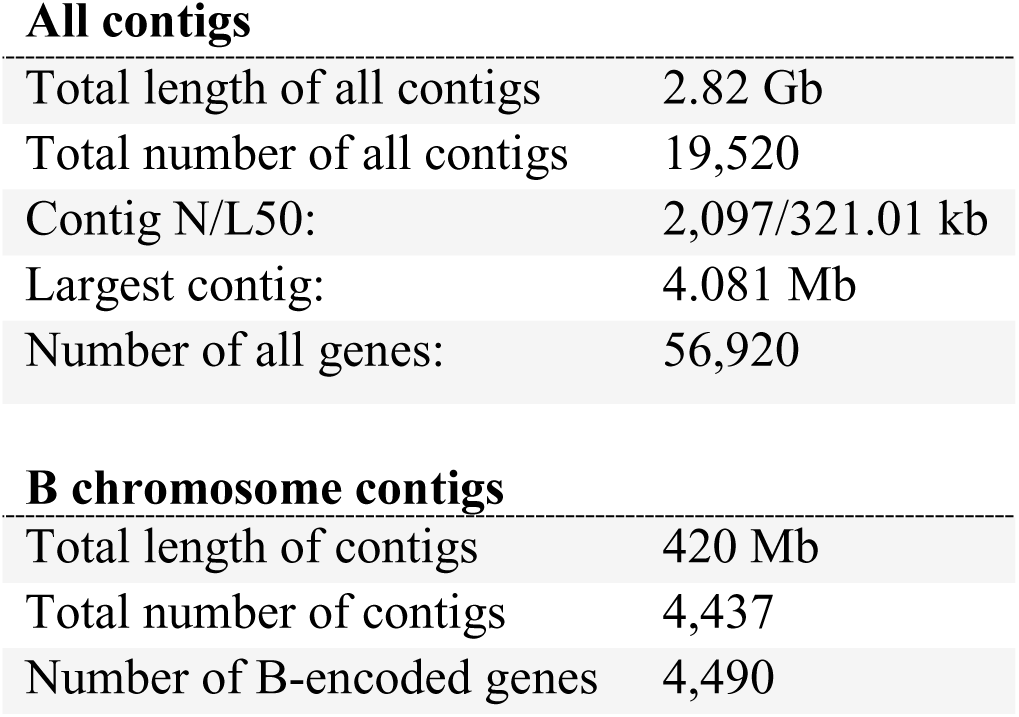
Genome assembly and annotation summary.

To further investigate the origin of B chromosome-located genes, a blast search of those genes was performed against *S. bicolor* genes [31] (Fig. 2D). The analysis identified 2,973 B-located genes out of total 4,490 with the hit to *S. bicolor* proteins. Syntenic analysis revealed no obvious standard A chromosome as progenitor of the B chromosome, instead the B chromosome is composed of multi-A chromosome derived sequences (Fig. 2D). This results are consistent with previous studies suggesting that genes found on B chromosomes are primarily derived from numerous A chromosomes of the particular species carrying the B chromosome [15, 32, 33, 34].

Coding sequences represented 1.3% of the length of B chromosome contigs. This is notably lower than the 3.9% gene content observed in the A chromosomal sequence, indicating a relative depletion of genes within the B-contigs (Fig. 2E). Conversely, B-contigs showed a higher proportion of repetitive sequences (60%) compared to 55% in the A chromosomes. While LTR retrotransposons are slightly accumulated in the B chromosome, DNA transposons are more abundant in the A genome (Fig. 2E, Additional file 1: Table S1). This pattern suggests that B chromosome accumulates some repetitive sequences during evolution.

To address whether the centromeric sequence composition of the A and B chromosomes differs, we retrieved and assembled reads containing centromeric repeats [30]. This *de novo* centromere assembly comprise 431 contigs with a cumulative length 7.57 Mb, across which the 137 bp tandem repeat SpuCL4 [30] represents over 86 % of the cumulative contig length. Due to highly repetitive nature of the sequence, it was difficult to extract contigs of B chromosomal origin using the same kmer approach as for genome assembly. Only 13 contigs (245 kb) were assigned to the B chromosome. Interestingly, we identified a novel B chromosome-specific tandem repeat (SpuCL166) that co-occured with canonical centromeric repeat in six contigs (Fig. 3A). Fluorescence *in situ* hybridization confirmed that tandem repeat SpuCL166 is located in (peri)centromeric region of the B chromosome, where it colocalize with the A and B chromosome shared centromeric repeat SpuCL4 (Fig. 3B-E). Also, the signal size of the SpuCL4 centromeric repeat (Fig. 3D; in green) seems to be larger on B than on A chromosomes. Thus, the (peri)centromere sequence composition differs between A and B chromosomes, with the B chromosome expanded by the presence of SpuCL166.

**Fig. 3:**
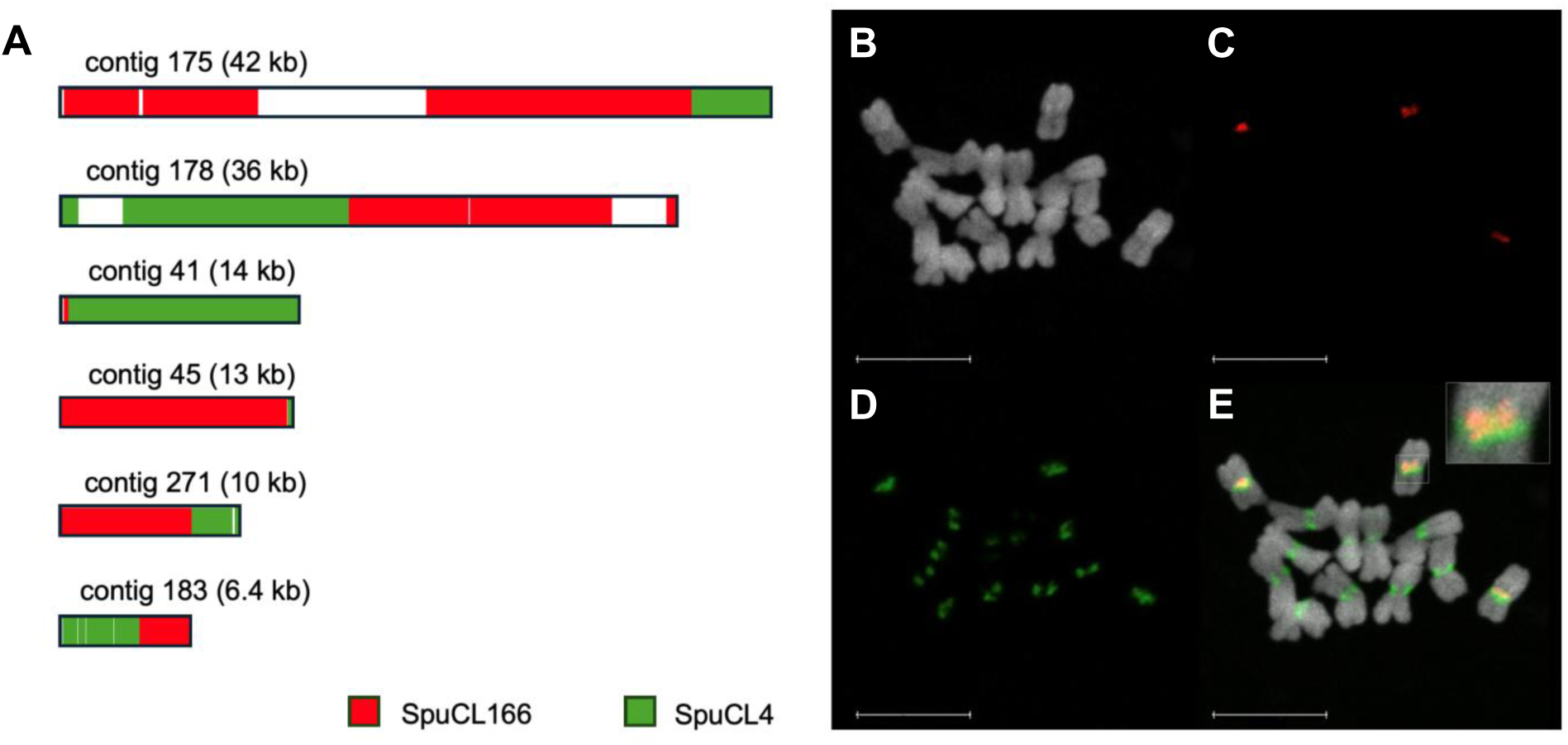
Composition of B chromosome centromere. **A** Assembled contigs containing both the B-specific tandem repeat SpuCL166 (red) and the centromeric repeat SpuCL4 (green). Out of 431 contigs, only six were found by BLAST to contain both repeats. In all six contigs, the SpuCL166 and centromeric SpuCL4 sequences colocalize. **B-D** Co-localization of centromeric repeat SpuCL4 and B-specific repeat SpuCL166 on mitotic chromosomes of *Sorghum purpureosericeum* plant harbouring 3 B chromosomes using FISH. **B** DAPI counterstained mitotic chromosomes, **C** signal of B-specific repeat, **D** signal of centromeric SpuCL4 repeat, **E** merged picture showing clear co-localization of B-repeat with the canonical centromere sequence with the detail of B chromosome centromere in right upper corner. Scale bar: 10 µm.

### Presence of the B chromosome influences the transcriptome of S. purpureosericeum embryos

To address the expression and effect of the B chromosome-encoded genes during the process of B chromosome elimination, we performed RNA-seq of whole embryos and compared the mRNA abundance between +B and 0B embryos. In total, 81.57 Gb of 2×150 bp pair-end reads were generated for all samples comprising three biological replicates at each developmental stage — early (∼7 DAP), mid (∼14-21 DAP) and late (∼28 DAP) (Fig. 4A; Additional file 4). The presence or absence of the B chromosomes was determined through PCR screening on the corresponding endosperms. On average, 86.3 ± 9.1 % of reads exhibited unique mapping to the *S. purpureosericeum* genome (Additional file 4).

**Fig. 4:**
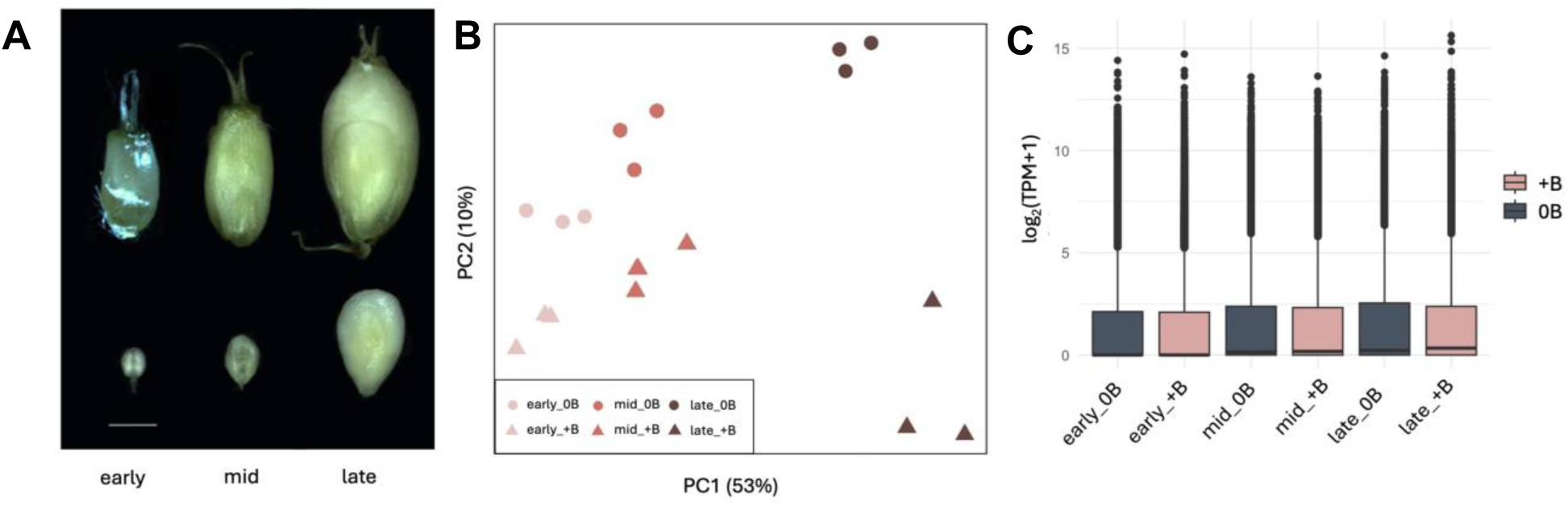
Transcriptome profiling of *S. purpureosericeum* embryos during development. **A** Stages of embryo development analyzed for transcriptomic profiling. Scale bar – 1 mm. **B** Principal Component Analysis (PCA) variance of 18 samples. **C** Boxplot showing the log (TPM+1) transformed expression levels of all genes across RNA-seq samples representing different developmental stages (early, mid, late) and conditions (+B and 0B). Each boxplot represents the average expression values across three biological replicates for each group. The central line within each box indicates the median expression level, while the box boundaries correspond to the interquartile range (IQR) (25th to 75th percentiles). Data points beyond the whiskers are plotted as outliers. Y-axis represents Log2 (transcripts per million (TPM)+1) transformed transcript levels of embryo RNA-seq samples.

We conducted a principal component analysis (PCA) to assess differences in gene expression among developmental stages in the presence and absence of the B chromosome. Clustering analysis showed expected grouping based on the age of the embryos, as evidenced by the separation along PC1 (Fig. 4B). The B chromosome appears to cause a visible shift in the expression profiles (10% PC2 variance) in all embryo stages. However, the developmental stages had a more substantial influence on the transcriptomic profile than the presence of B chromosome, explaining 53% of the variance along PC1. Notably, the most distant cluster comprised the late +B samples. Similarly, the median gene expression was highest for late +B samples, suggesting that the B chromosome had the biggest effect on the gene expression in mature embryos (Fig. 4C).

### Differential expression and gene ontology unveiled 21 candidate genes associated with B chromosome elimination

With the aim of identifying candidate genes involved in the process of the B chromosome elimination, we conducted differential expression and gene ontology analyses. Differential expression analysis in the early-embryonic stage identified 378 upregulated and 1738 downregulated genes in the presence of the B chromosome (Fig. 5A; Additional file 5). In the mid-embryonic stage, 135 upregulated and 131 downregulated genes were identified in the same comparison (Fig. 5A; Additional file 5). Finally, 205 and 33 genes were up- and downregulated in the presence of the B chromosome in late embryos (Fig. 5A; Additional file 5). Although we have to consider that also the A-located genes may contribute to the elimination process, we presume that the majority of the genes controlling B chromosome elimination are transcribed from the B chromosome (i.e. upregulated in +B compared to respective 0B samples) as was proven in the B chromosome elimination in *Ae. speltoides* [29, 35] and nondisjunction phenomenon in maize [36, 37, 38] and rye [34]. Consequently, the location of upregulated genes was compared to the B chromosome contig set (Additional file 2). This comparison revealed 92 unique B chromosome-located upregulated genes, with the distribution across the stages as follows: 33 genes upregulated at the early-stage, 54 genes at the mid-stage and 35 genes at the late-embryonic stage (Fig. 5B; Additional file 5). Notably, only three genes were shared across all stages, and 19 genes were shared between the mid-stage and late-stage (Fig. 5B).

**Fig. 5:**
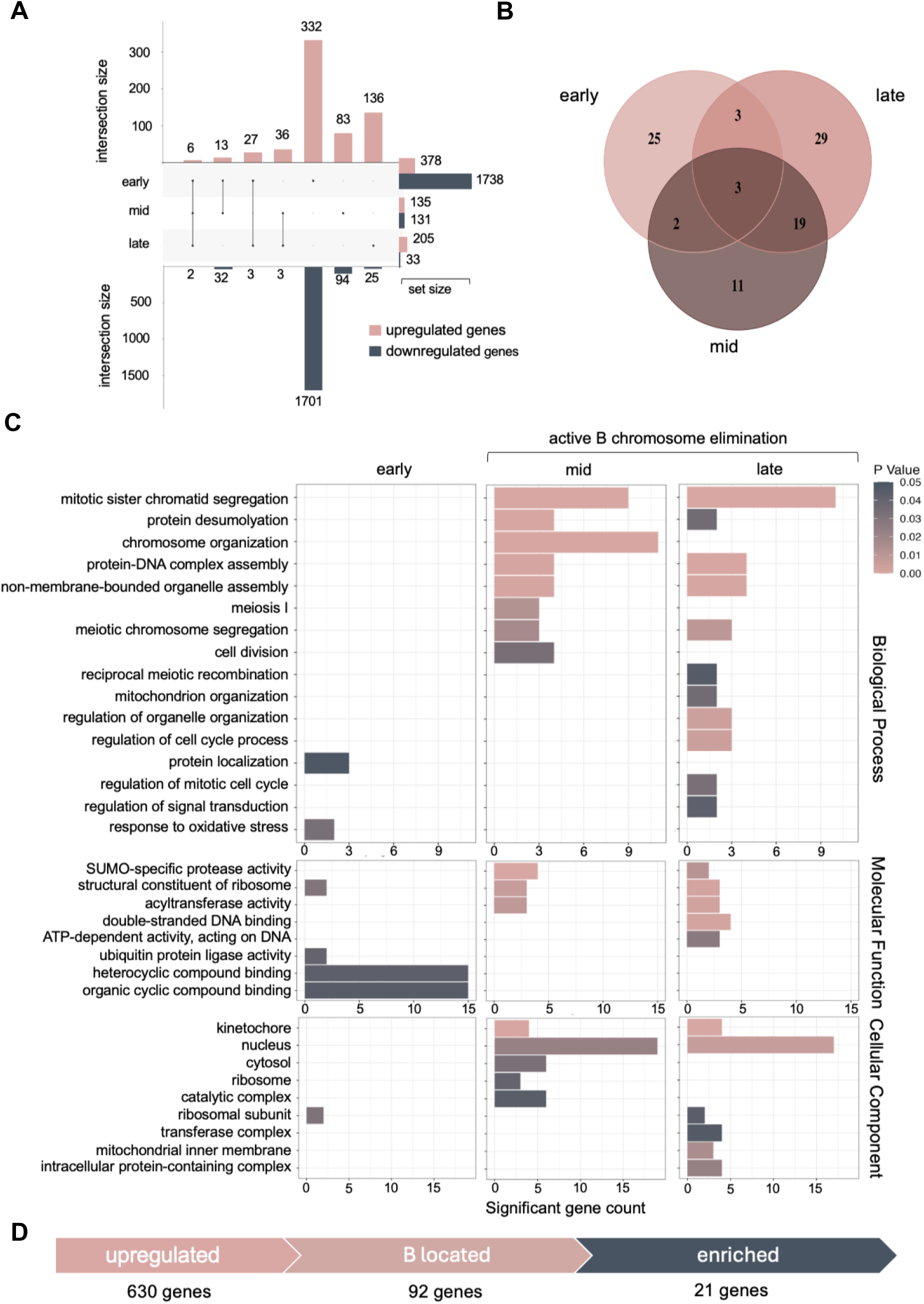
Comparative analysis of differentially expressed genes in the presence of the *S. purpureosericeum* B chromosome at three developmental stages. A UpSet plot showing the overlaps and unique genes resulting from differential analysis of gene expression in embryos in three comparisons: early_+B vs early_0B, mid_+B vs mid_0B and late_+B vs late_0B. Top – upregulated genes, bottom – downregulated genes. **B** Venn diagram showing the overlaps of B chromosome located upregulated genes. **C** Gene set enrichment analysis of upregulated B-located genes. The barplot visualizes all significantly enriched gene ontology (GO) terms (p-value < 0.05). Bar length represent number of identified upregulated genes; colour correspond to significance level. **D** Process of candidate gene selection.

Subsequently, a gene set enrichment analysis (GSEA) was employed, comparing upregulated B-localized genes for each stage with all B-localized genes as the reference set (Additional file 6, 7). This comparison was used to specifically focus on the B chromosome-encoded candidates. Sixteen significantly enriched GO terms were identified among Biological processes in three developmental stages (2/8/11 for early/mid/late embryo, respectively) with some overlapping terms between mid- and late-stage (Fig. 5C). Similarly, eight terms (4/3/5) were enriched among Molecular functions and nine terms (1/4/6) among Cellular component GO terms, again with some terms shared. Clearly, some terms are enriched in both the mid- and late-developmental stages (i.e. in the stages when the B chromosome is eliminated). Among them, GO:000070 - *mitotic sister chromatid segregation*, GO:0065004 - *protein-DNA complex assembly*, GO: 0016926 - *protein desumoylation*, GO:0016747 - *acyl transferase activity*, GO:0016929 - *deSUMOylase activity*, GO:0005634 - *nucleus*, and GO:0000776 - *kinetochore* appear to be the most intriguing for the B chromosome elimination process. Furthermore, the GO terms that are enriched either in the mid- or late-stages should be considered for potential significance. These comprise GO:0003690 - *double-strand DNA binding activity*, and GO:0008094 - *ATP-dependent activity acting on DNA*. Altogether, these terms represent 21 genes overexpressed specifically in samples with the B chromosome in mid-, late- or both developmental stages (Table 2, Fig. 5D).

**Table 2:**
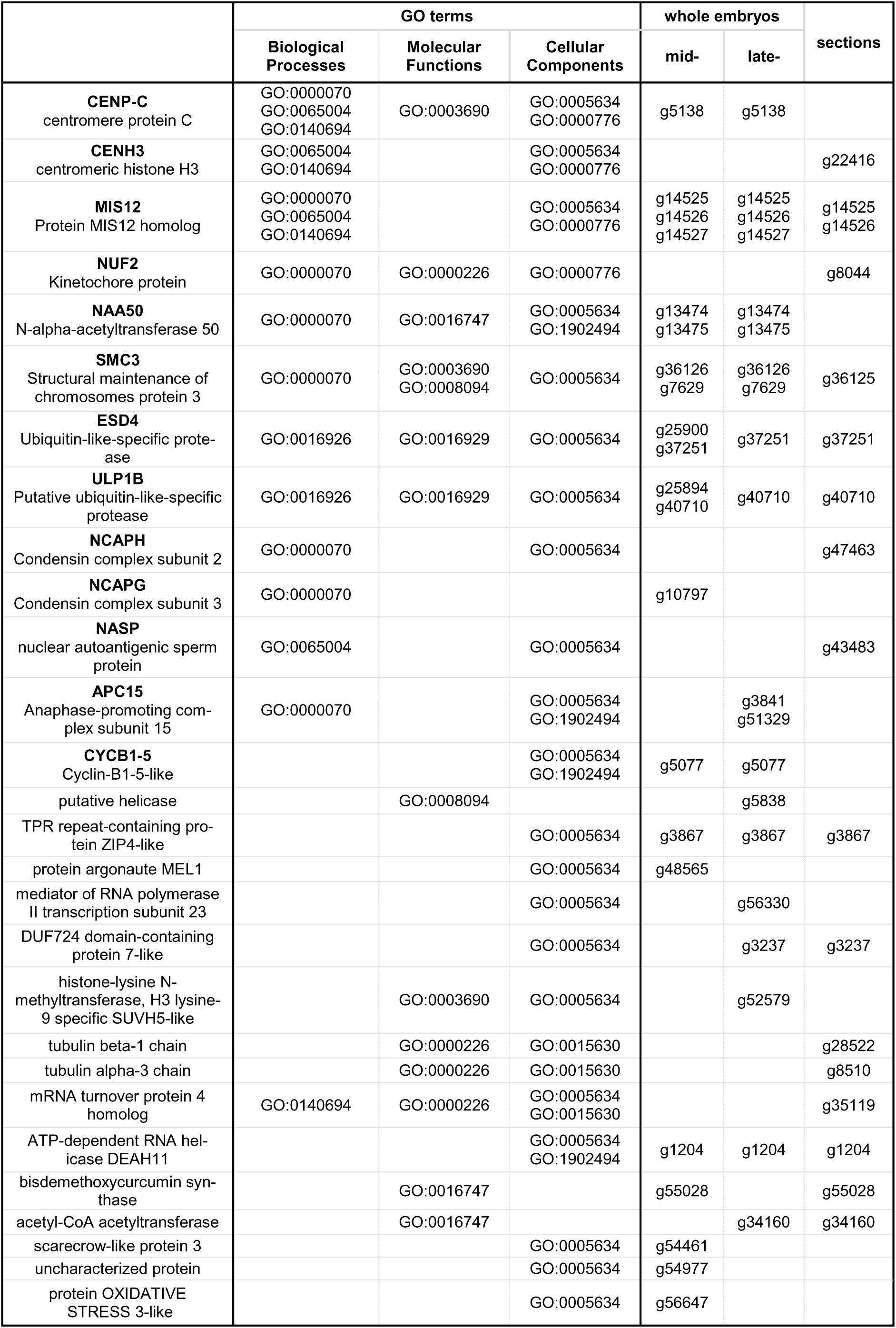
List of candidate proteins controlling B chromosome elimination raised from analysis of transcriptomes of whole embryos and LCM samples.

### Transcriptome analysis of dissected subregions of mid-stage +B/0B embryos extended the list of candidate genes

To ensure the candidate genes were not overshadowed within the broader context of whole embryo gene expression from regions where B-elimination does not occur, an additional sample set was utilized. It is composed of laser-captured microdissected (LCM) regions of embryos at the mid-developmental stage (Additional file 1: Fig. S3, S3), where B chromosome elimination occurs. The differential expression analysis between +B (with active B chromosome elimination) and 0B samples identified 81 upregulated and 51 downregulated genes (Additional file 1: Fig. S4, Additional file 5). Forty-six upregulated genes were found to be encoded by the B chromosome, and out of these, ten belong to significantly enriched GO terms identified in whole-embryo transcriptome analysis (Table 2, Additional file 1: Fig. S4). Further, two additional relevant enriched GO terms have been identified that might be associated with the B chromosome elimination process - GO:0000226 - *microtubule cytoskeleton organization*, and GO:0015630 - *microtubule cytoskeleton (*Additional file 1: Fig. S5). Altogether, the candidate gene set was completed by seven additional genes, which were found to be upregulated exclusively in LCM-generated samples.

The final set of identified genes represents a distinct group of candidates related to cell division. The function of many of them is well described, while the effect of others is less apparent. Among the genes, we detected the crucial components of both inner (*CENH3*, *CENP-C*) and outer kinetochore (*MIS12*, *NUF2*). Further, the genes encoding components of the cohesin (*SMC3*) and condensin (*NCAPG*, *NCAPH*) complex and the genes controlling cell-cycle checkpoint (*APC15*, *CYCB1-5*) can be considered as meaningful candidates associated with the B chromosome elimination process.

### B chromosome-located candidate genes encode proteins distinct from the corresponding A chromosome-encoded variants

To assess the functional capabilities of the candidate genes, we investigated whether there are amino acid differences between the particular candidate gene product and the corresponding A chromosome encoded protein. We aligned amino acid sequences with the EMBOSS Needle Pairwise Sequence Alignment tool [39] as only non-synonymous mutations (and truncations) might potentially alter the function of the candidate proteins. The alignments highlighted single amino acid polymorphisms (Additional file 8) as well as truncation of some B-specific variants of proteins. For instance, the A- and B-variants of CENH3 differ in two amino acid residues, with an overall sequence identity of 98.7%. In other cases, the degree of sequence identity ranged from 99.01% for DEAH11, to as low as 40.82% for one of the copies of ESD4 (Additional file 8). Some genes were found in multiple copies on the B chromosome, but with the exception of *ESD4*, they encode proteins identical in their amino acid sequence. The only candidate without polymorphism between A and B chromosomal copies was histone-lysine N-methyltransferase SUVH5-like, which showed 100% amino-acid sequence identity.

As one of the criteria for the selection of candidate genes was their localization on the B chromosome, the *in silico* assignment was validated by PCR for the selected genes *CENH3*, *CENP-C*, *SMC3*, *APC15*, *CYCB1* and *NAA50*, all of which have been demonstrated to play a specific role in mitotic processes. We designed primers specific to polymorphic sequences that differentiate between the A and B chromosome variants (Additional file 1: Table S3). The PCR results confirmed that all these genes are located on the B chromosome, as the amplification occurred exclusively in the +B samples (Additional file 1: Fig. S6). This list of candidate genes suggests that critical changes leading to B chromosome elimination affect the cellular machinery involved in chromosome segregation. In particular, the expression of B-variants of these genes suggests that the elimination might result from an interplay between complex changes at the level of kinetochore structure and chromatid cohesion controlled by the modified checkpoint.

The network of genes orchestrating proper chromosome distribution is likely complex and our list of candidates is not necessarily complete. Unfortunately, some relevant genes may have been missed due to statistically non-significant expression changes. Furthermore, spurious expression of some B chromosome localized genes in 0B samples was detected (Fig. S7), hampering detection of some candidates. This could be caused by false positive assignment of contigs to the B chromosome. In additon, approximately ∼7 % of genes exhibit perfect homology to their A chromosomal homologs.

### CENH3 displays amino acid changes on the modelled interaction interface with CENP-C

Among the detected candidate genes, CENH3 and CENP-C represent known interacting partners [41–43]. To address the polymorphism between the A and B chromosome encoded CENH3 variants, we modelled the CENH3-containing nucleosomes using AlphaFold3 [40] (Fig. 6). As a representative of CENH3-interacting partners, we used CENP-C [41] and to this end, we modelled the complex with the C-terminal domain of CENP-C (AA 686 to 764). This domain is responsible for the CENP-C interaction with the nucleosome complex (Fig. 6A) [42, 43]. Focusing on the interplay between CENH3 and CENP-C, we identified that both amino acid residues distinct between the CENH3 variants are located at the interface between CENH3 and CENP-C (Fig. 6D). Similarly, we mapped some of the differences between A- and B-chromosomal variants of CENP-C to its C-terminal domain, with the mutations likely affecting the interacting interface between CENH3 and CENP-C. More in detail, analysis of Protein Interactions by Surface Area (PISA) revealed that I92V in CENH3 directly affects amino acid residue involved in the interaction, while I144M is located immediately adjacent to the interacting site (Fig. 6E, Additional file 9). Similarly, the polymorphisms E709K and V758I in CENP-C were identified as adjoining interacting amino acids. The model suggested complementary changes in B chromosomal variants of the inner kinetochore proteins (CENH3 and CENP-C), thus not excluding the formation of an altered kinetochore on the B chromosome.

**Fig. 6:**
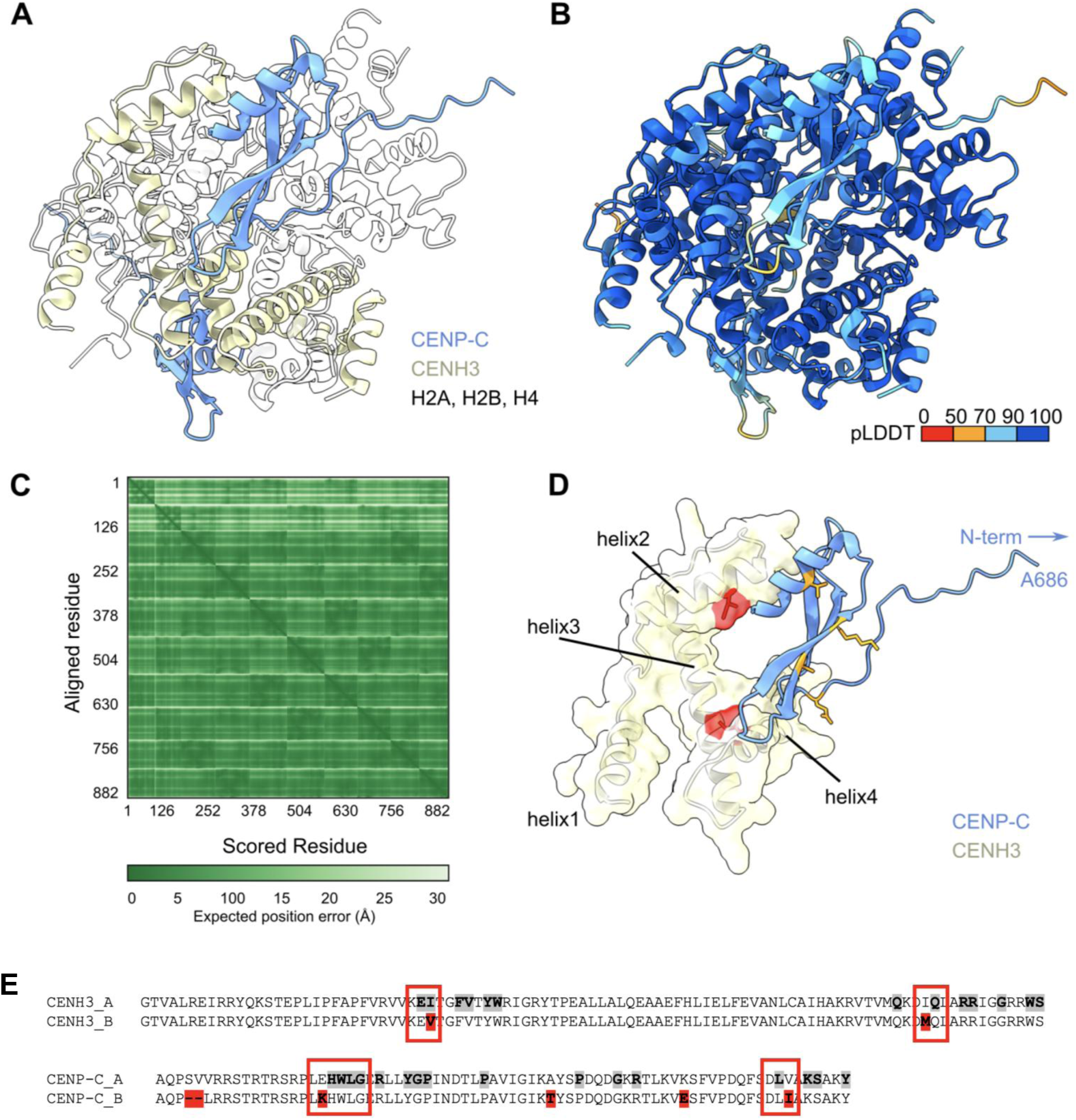
Differences in the CENH3 A and B variants are located at the interface between CENH3 and CENP-C in the nucleosome complex. **A** CENH3-CENP-C nucleosome complex was predicted using AlphaFold3. CENP-C is highlighted in light blue, CENH3 in light yellow, and other histone proteins are shown in transparent white. **B** CENH3-CENP-C nucleosome complex coloured based on the pLDDT (the predicted local distance difference test) score. All subunits of the CENH3-CENP-C nucleosome complex are predicted with high confidence. **C** Expected position error for the CENH3-CENP-C nucleosome complex showing that the position of individual complex subunits in respect to each other is confidently predicted. **D** Only CENP-C and CENH3 are shown. In addition to the ribbon representation, the transparent solvent excluded surface of CENH3 (light yellow) is shown with highlighted amino acid residues that differ between CENH3 variants (red). The CENP-C polymorphism is highlighted in orange. **E** Graphical visualization of PISA analysis for CENH3 and CENP-C. The interpolated alignments illustrate the interaction sites between the CENH3_A protein variant and CENP-C_A protein variant. Interacting residues are highlighted in bold and grey-undercolored. Polymorphisms detected in B-encoded protein variants are undercolored in red. The key mutations are highlighted in red boxes. For clarity, only the subdomains of CENH3 (residues 62-156) and CENP-C (residues 692-764) are depicted.

### Similar changes occurred in the evolution of B chromosome-specific CENH3 variants in maize and Ae. speltoides

Polymorphisms in the A- and B-encoded CENH3 variants in *Sorghum* suggested that its interacting interface might be functionally modified. To test whether similar evolutionary forces shaped the CENH3 of other plant B chromosomes too, we mapped differing amino acid residues on the Alphafold3 structure for two other species harbouring B chromosomes encoding CENH3, namely *Z. mays* and *Ae. speltoides* [34, 33]. Interestingly, most polymorphisms between A- and B-encoded copies of the gene were indeed located in helix2 and helix4 of CENH3, corresponding to the same interface as in *Sorghum* CENH3 (Fig. 7). To support the idea that the same changes evolved repeatedly, we performed a phylogenetic analysis of CENH3, including their B-specific variants (Fig. 7E, Additional file 10). In *Sorghum* and maize, A chromosome-encoded copies of CENH3 were the closest relatives for B-specific copies of the gene. In *Ae. speltoides*, the B-chromosomal variant is more distal to the A-chromosomal one than CENH3 in closely related bread wheat. Altogether, these results indicate independent evolution in each species.

**Fig. 7:**
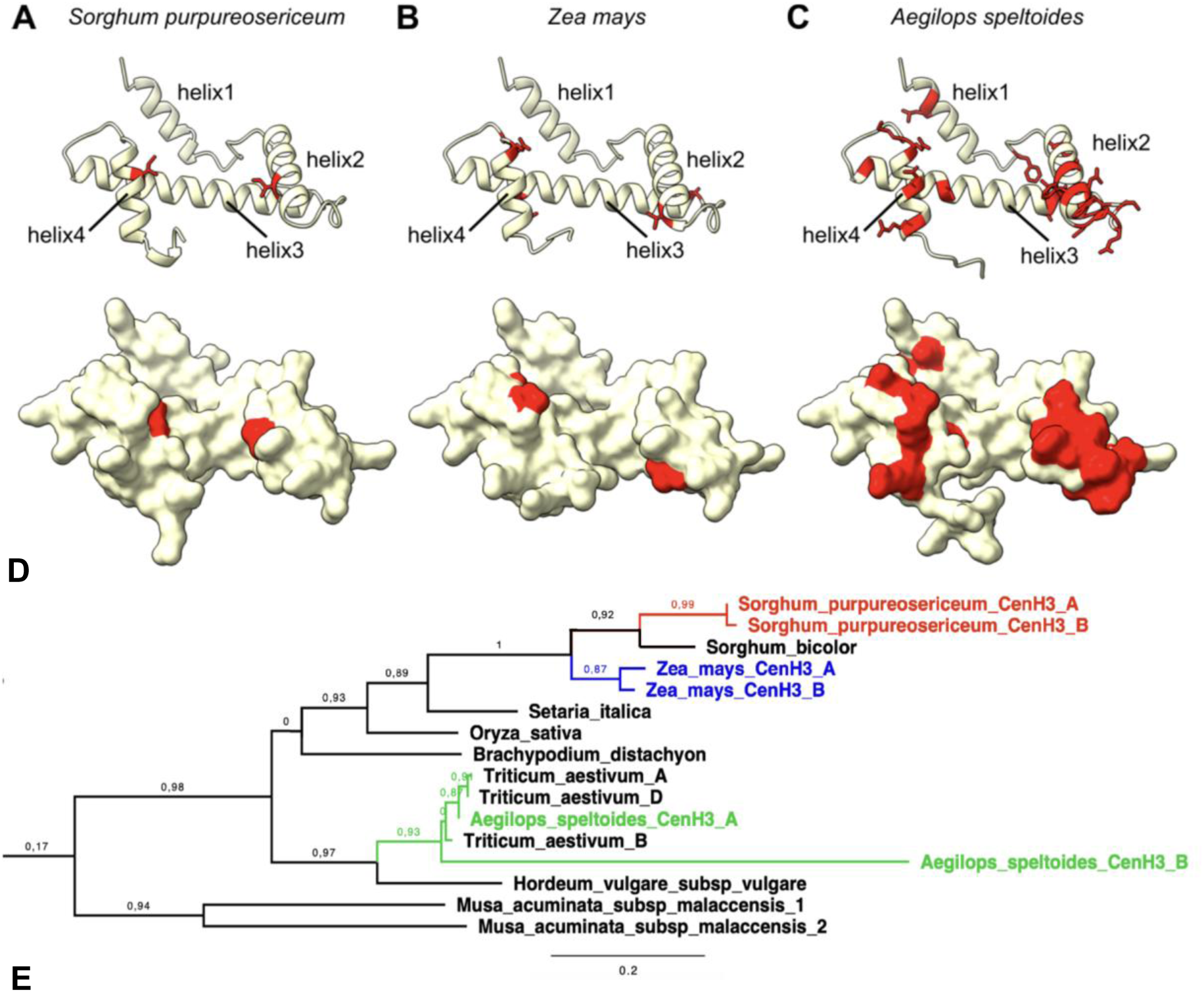
Evolutionary comparison of polymorphism between CENH3 variants. **A** AlphaFold3 prediction of the *Sorghum purpureosericum* CENH3 structure (unstructured amino acids 1-62 are not shown for clarity) with highlighted amino acid residues that differ between CENH3 variants (red). The ribbon representation is shown on the top with marked helices in the structure. On the bottom, the solvent-excluded representation is displayed. **B** AlphaFold3 prediction of the *Zea mays* CENH3 structure (unstructured amino acids 1-61 are not shown for clarity) with highlighted amino acid residues that differ between CENH3 variants (red). The ribbon representation is shown on the top with marked helices in the structure. On the bottom, the solvent-excluded representation is displayed. **C** AlphaFold3 prediction of the *Aegilops speltoides* CENH3 structure (unstructured amino acids 1-62 are not shown for clarity) with highlighted amino acid residues that differ between CENH3 variants (red). The ribbon representation is shown on the top with marked helices in the structure. On the bottom, the solvent-excluded representation is displayed. **D** Protein sequence alignment of CENH3 A and B variants in *Sorghum purpureosericeum, Zea mays* and *Aegilops speltoides*. **E** Phylogenetic tree of CENH3 proteins in monocot species. Proteins from species possessing B chromosomes are highlighted in red (*Sorghum purpureosericeum*), green (*Aegilops speltoides*), and blue (*Zea mays*). The full phylogenetic tree is available as Additional file 11.

## Discussion

### Sequence composition of the Sorghum purpureosericeum B chromosome supports the general model of B chromosome origin

B chromosomes are undoubtedly spectacular genetic elements and their origin remains an evolutionary mystery. Despite being non-essential or even harmful to the host, if the number of B chromosomes is high, they exist in thousands of species. The widely accepted hypothesis of B chromosome origin assumes that B chromosomes are derived from different A chromosomes as a by-product of the evolution of the standard karyotype [76, 77, 78, 79]. More recently, it was proposed that proto-B chromosomes represent a chromoanagenesis product. In the second step, proto-B chromosomes accumulate additional multi-A chromosome sequences over time. Consequently, the original structure of the early stage proto-B chromosomes becomes masked by continuous sequence incorporation [80]. As such, B chromosomes share the main sequence profile with the chromosomes of the standard complement. Indeed, all sequence types from coding genes to the most abundant repeats can be found in the sorghum B chromosome (see Fig. 2E). A comparable evolutionary trend was found in maize and rye [32, 33]. Nevertheless, the B chromosomes usually accumulated more repetitive sequences than A chromosomes. This is due to the distinct evolutionary path they take. Except for genes required for the drive of B chromosomes, the majority of sequences carried on B chromosomes (including genes) are not subjected to selection pressure as genes encoded by A chromosomes [34, 33]; and thus provide a “save harbour” for the sequences of various types which can be freely inserted. This absence of selection makes the B chromosome prone to mutation, allowing a high activity of transposons resulting in high evolutionary dynamics of the B-located sequences (reviewed in [81, 79]). Interestingly, the evolution of the B chromosomes led to the emergence of B-specific repetitive sequences [30, 82, 83, 84]. Typically, they are abundant tandem-repeat sequences organized in large blocks at particular chromosome loci. In *S. purpureosericeum*, three such B-specific repeats have been described [18, 30]. In this study, we identified another tandem repeat that is located in the (peri)centromeric region of the *S. purpureosericeum* B chromosome and might be involved in the elimination process of the B chromosome. This repeat (SpuCL166) has no apparent homology to any *S. purpureosericeum* sequence. The only similar sequences (with approximately 70% similarity) found in available databases are satellite sequences *Pp*TR-2, and *Pp*TR-3 from *Poa pratensis* [85], indicating a unique composition of the (peri)centromeric region of the *S. purpureosericeum* B chromosome.

### Sorghum B chromosome encodes candidate genes for its elimination

Although the precise mechanisms of B chromosome elimination remain unclear, our findings provide valuable insights into this process. We identified a set of B chromosome-specific genes that are upregulated in tissues undergoing B chromosome elimination and, at the same time, cluster in GO terms linked to mitosis-related processes. Our list of candidate genes potentially contributing to the B chromosome elimination process includes genes whose A chromosome copies are known to be involved in kinetochore establishment [44], sister chromatid cohesion [45] and regulation of chromosome segregation during mitosis [46]. This suggests that the B chromosome might control its own transmission/elimination by expressing B-specific paralogs of those mitosis-related genes. If B chromosome elimination is indeed a strictly controlled process [15, 25, 47, 48], the detection of numerous B-specific kinetochore and cohesin subunits expressed during active elimination supports this hypothesis.

Our transcriptomic data revealed that the B chromosome expresses specific variants of a surprisingly high number of kinetochore proteins (CENH3, CENP-C, MIS12, and NUF2) during its elimination. The same or other B-specific kinetochore components were detected in available transcriptomes of maize, rye and *Ae. speltoides*, where the B chromosomes display a chromosome-specific performance (chromosome elimination and or chromosome drive) [29, 33, 34]. Previously it has been shown that mutations or overexpression of kinetochore genes can disrupt cell division and promote chromosomal missegregation [49, 50]. In unstable plant and animal hybrids, uniparental chromosome elimination occurs due to e.g. the failure of spindle attachment resulting from kinetochore misassembly [50, 51, 52]. However, eliminating B chromosomes in *Ae. speltoides* clearly show microtubule attachment and CENH3 immunosignals [15], which may indicate that the elimination of B chromosomes proceeds via a different mechanism. The lagging of the B chromosome, leading to its elimination, is likely caused by factors other than missing attachment to microtubules.

In addition, dysregulated sister chromatid cohesion likely contributes to B chromosome elimination. Commonly, sister chromatids are held together by the cohesin complex, which encircles the DNA and is cleaved by the enzyme separase during the metaphase-to-anaphase transition [53, 54]. Interestingly, the B chromosome encodes specific variants of cohesin subunit SMC3, anaphase-promoting complex subunit APC15, and the regulatory protein CYCB1, which modulates separase activity. Mutations in the cohesin subunits and the cohesion-related genes have been the subject of intense research as they induce chromosomal missegregation resulting in tumorogenesis [55, 56, 57, 58, 59]. Cancer-related mutation affecting cohesion leads frequently to aneuploidy caused by missegregation of chromosomes [60], occasionally resulting in micronuclei formation and chromosome loss [61]. Another candidate, APC15, is known for the regulation of the interaction of anaphase-promoting complex (APC) and spindle assembly checkpoint (or Mitotic checkpoint complex; MCC). Depletion of APC15 enhances MCC interaction with the APC [62, 63]. The presence of a B-specific variant of *APC15* suggests that the APC might be regulated differently in +B cells, e.g. by prolonging APC inactivation beyond spindle attachment control through MCC-mediated inhibition. Nevertheless, the role of APC15 as well as other candidate proteins in B chromosome elimination requires further validation.

## Conclusion

Through detailed spatio-temporal analyses, we demonstrated that elimination predominantly occurs during mid-embryogenesis and is largely completed by the late embryonic stage, with residual B chromosomes persisting only in specific meristematic tissues. This precise timing, coupled with the identification of 28 B chromosome-encoded candidate genes and novel centromeric repeat SpuCL166, underscores the active role of the B in directing its own fate. Our findings highlight the complexity of the elimination mechanism and can pave the way for a deeper understanding of the intricate cellular processes underlying the B chromosome behaviour. Despite advances in knowledge and methodology, we are still far from fully under-standing the mechanism of elimination in plants and any new findings will be highly valuable.

## Methods

### Plant material and cultivation

Seeds of diploid wild sorghum *Sorghum purpureosericeum* (Hochst. ex A.Rich.) Schweinf. & Asch. were used in all experiments - 2n = 2× = 10, for plants without B chromosome (0B); 2n = 2× = 10 + 2B, for plants possessing two B chromosome(s). Sown seeds represented the progeny of the original seed stock obtained from genebank ICRISAT (accession no. 18947). Plants were cultivated at the short-day conditions of 10-hours daylight and temperatures of 27°C (day) and 22°C (night). Upon the initial emergence of spikes, the presence/absence of B chromosomes within the plants was assessed through flow cytometry, employing the method refined by Karafiátová et al. [30].

### Reference sequence and annotation

#### Whole genome sequencing

Whole genome sequence of 0B and +B plant was obtained using combination of Illumina and Oxford Nanopore Technologies (ONT). High-quality genomic DNA for Illumina sequencing was isolated from non-fixed young inflorescence using flow cytometry. Roughly 60,000 diploid somatic nuclei from single plant without B chromosome (0B) and plant carrying two B chromosomes (+B) were sorted for Illumina sequencing. To generate Illumina reads, sequencing library was prepared using NEBNext® Ultra™ II DNA Library Prep Kit (NEB, Ipswich, MA, USA), followed by pair-end sequencing run on Illumina NovaSeq 6000 platform. Subsequently, adapter trimming and quality filtering were conducted using the fastp tool with specified parameters (-q 30 -l 50) [86].

High molecular weight (HMW) DNA for ONT was isolated from *Sorghum purpureosericeum* fixed young inflorescence of +B plants. Key steps included flow cytometry for nuclei sorting, treatment with SDS and proteinase K, concentration through centrifugation, and AMPure purification (Additional file 12). Finally, isolated HMW DNA was used for creating sequencing library using Rapid Sequencing Kit protocol (catalog No. SQK-RAD004). Oxford Nanopore MinION sequencing was conducted over 14 runs, with each run lasting 3 days, and each run performed on its own dedicated flow cell. Generated long reads were basecalled with the Guppy5+ basecaller (provided by ONT) and validated using NanoPlot [87].

#### De novo genome assembly and contigs assignment

ONT reads were used for SMARTdenovo genome assembly [88] with specified parameters (-c 1, -e dmo, -k 23). Subsequently, the raw assembly underwent polishing through medaka v1.6.0 (utilizing ONT reads) [89] followed by NextPolish v1.4.1 (employing Illumina reads) [90]. Genome statistics were assesed using QUAST v5.0.2. [91]. The quality assessment of the assembly was performed using BUSCO v5.2.2 with the *Poales* lineages dataset [92]. For contig sorting, 39-mer indexes were generated from the polished reference sequence, as well as the raw B chromosome positive (+B) and B chromosome negative (0B) Illumina reads representing 20x genome coverage using genome tools. A custom Perl script was utilized to identify diagnostic (low-copy) regions, facilitating the removal of repetitive sequences and the exploration of k-mer counts between the reference and +B/0B datasets. The status of B-contigs was assigned automatically when the ratio of +B count to 0B count for 39mers over diagnognostic regions was ≥ 1.1. Contigs meeting this criterion were classified as B-contigs with high confidence. However, due to the high similarity between the B chromosome and the A chromosomes, as well as the occurrence of chimeric A/B contigs, we also classified certain contigs as low-confidence B-contigs when the ratio of +B count to 0B count for 39mers was > 1 and ≤ 1.1. For low-confidence B contigs assignment was further supported by RNA-seq data. Contigs with a ratio of ≤ 1 were automatically designated as A-contigs. To verify the separation of B-contigs into subgroups based on their k-mer counts, we conducted a Welch t-test on the B-contigs group to assess significant differences between 0B and +B k-mer counts with resulting p-value 2.2e-16. The sequence of 137-bp tandem repeat identified to be part of *Sorghum purpureosericeum* centromeres [30] was aligned to raw ONT reads. Extracted (centromeric) reads were assembled into contigs using SMARTdenovo with the same settings as for genome assembly.

#### Genome annotation

First, a custom-made library of repeats was constructed by using entire reference sequence as input for Domain Based Annotation of Transposable Elements (DANTE) tool [93]. DANTE library was then used by RepeatMasker to soft-mask the reference sequence [94] and masked reference was subsequently used to generate full gene structure annotations with BRAKER3 pipeline [95]. For model training as well as evidence for identified coding sequences, a set of embryo +B RNA-seq smaples (Additional file 4) and OrthoDB proteins database for lineage Poaceae v11 [96] (https://www.orthodb.org) were used. RNA-seq reads were mapped to the unmasked reference sequence using HISAT2 v2.1.0 [97] with max-intronlen 50000 parameter. For annotation of UTRs, parameter --addUTR was used to run BRAKER3 pipeline. The quality of the final annotation was evaluated by BUSCO v5.2.2 [92] in the protein mode. The content of A- and B-contigs was assessed based on the composition of coding sequences, tandem repeats, and transposable elements, as determined by genome annotation, RepeatExplorer, and RepeatMasker [94] outputs. Genes encoded by B chromosome contigs were copared against *Sorghum bicolor* NCBIv3 genes set ([31]; max_target_hit=1). The syntenic relationship between B chromosome contigs and *S. bicolor* chromosomes was visualized using Circos (v0.69.9) [98].

### Embryo development and visualization of B chromosome elimination in situ

#### Staging of embryo development

Prior the experiments, dozens of embryos were extracted and lined up to create a developing scale, which served as a reference template for all subsequent experiments (Fig. 1A). To study the progression of B chromosome elimination, embryonic development was divided into three contrasting stages. Early-stage - ∼7DAP, mid-stage ∼14-21DAP and late-stage ∼28 DAP. Developing seeds were dissected and embryos isolated using stereomicroscrope SZX16 (Olympus LS, Tokyo, Japan) with CCD camera. The stage of development was assessed visually by analyzing stereoscope images (Fig. 1A).

#### Wax sections and staining

Developing *sorghum* seeds were isolated at three stages, dehulled and fixed in 4% freshly prepared paraformaldehyde (w/v) in dH_2_O, supplemented with 2% Tween-20 (v/v) and 2% Triton X-100 (v/v), adjusted to pH 7 using HCl, for 12 minutes under vacuum infiltration. Subsequently, the seeds were transferred to fresh fixative solutions and stored overnight at 4°C. Wax embedding and sectioning was perfomed according to Kovačik et al. [99]. Briefly, the fixed samples were rehydrated through ethanol series, cleared with a ROTIHistol series (Carl Roth, Karlsruhe, Germany), and embedded in Paraplast (Sigma-Aldrich). Longitudinal dorsoventral sections, 10μm-thick, were cut using a Reichert-Jung 2030 microtome and mounted on Adhesion Slides Superfrost Ultra Plus (Thermo Fisher Scientific). Prior staining with 1% toluidine blue, the slides were dewaxed with ROTIHistol and rehydrated.

#### Embryo cryo-sectioning and FISH on embryo sections

The developing seeds produced on +B plant from early-, mid- and late-developmental stages were collected and stripped of bran. While whole seeds were processed for early stages for ease of handling, the dehulled embryos were used for mid- and late-stages. The seeds/embryos were fixed in 4% formaldehyde (Sigma-Aldrich) in 1X PBS and then underwent cryo-protective treatment in sucrose series according to Karafiátová et al. [18]. The longitudinally oriented specimens were embedded onto pre-cooled specimen discs in a drop of Surgipath cryo-gel FSC22 blue (Leica, cat. no. 3801481). The sectioning was done using the cryostat Leica CM1950 (Leica Biosystems, Nussloch, Germany) at -18°C. 10µm-thick sections were collected onto the Silane-prep microscopic slides (Sigma-Aldrich, cat. no. S4651-72FA) and stored at 4°C till use. High-quality sections were selected for fluorescent *in situ* hybridization with B-specific probe.

FISH probe specific for the wild sorghum B chromosome was designed based on repetitive sequences identified with RepeatExplorer 2 [100] and TAREAN [101] tools (https://repeatexplorer-elixir.cerit-sc.cz). The analysis was done as described in Karafiátová et al. [30]. Sequence reads used in this analysis, the sequence of resulting repetitive clusters, and the counts of +B/0B reads in each cluster have all been submitted to the Zenodo repository (10.5281/zenodo.14197239). Four B-specific clusters (SpuCL137, SpuCL166, SpuCL193 and SpuCL220) were retrieved from the repeat analysis and used to design the cytogenetic marker for the B chromosome. Primers were designed using Primer3 online software [102, 103, 104] (Additional file 1: Table S2). For the clusters SpuCL137 and SpuCL220, the amplicon was amplified first and then probe was labelled with tetramethylrhodamine5-dUTP (Roche) using nick translation. Probes for clusters SpuCL166 and SpuCL193 were labelled using PCR with the same fluorochrome. The probe labelling was done according to Kara-fiátová et al. [30]. First, the cluster specificity was tested in PCR on 0B and +B genomic DNA. Subsequently, the specificity of the probes for the B chromosome was verified on meiotic metaphase chromosomes. Both methodologies followed protocols published by Karafiátová et al. [18, 30]. Next, specific probes were pooled and used as one robust marker for B chromosome detection.

Processing of the section and FISH was performed as described by Karafiátová et al. [18]. In brief, the embryo sections were immobilized on slide in thin layer of PAA gel. B-specific probe (TMR-labelled) targeting the B chromosome was hybridized onto the sections overnight. After the post-hybridization washes, the DNA was counterstained with 1.5 μg/ml DAPI. The signals were observed using Leica TCS SP8 STED 3X confocal microscope (Leica Microsystems, Wetzlar, Germany), equipped with an HC PL APO CS2 100×/1.44 Oil objective, Hybrid detectors (HyD), and the Leica Application Suite X (LAS-X) software version 3.5.5 with the Leica Lightning module (Leica, Buffalo Grove, IL, USA).

### Transcriptomics of developing embryo

#### Embryo harvesting and endosperm screening

Seeds at defined stages of development were collected from 0B and +B plants. Due to the irregular inheritance of B chromosomes, their presence in the developing embryo had to be determined. Both embryo and endosperm were separately placed in 1.5 ml microtubes, frozen in liquid nitrogen, squashed with a tube pestle and stored at -80 °C. In wild sorghum, the B chromosome status of the endosperm mirrors that of embryo for +B samples [18]. That way, B chromosome presence was verified noninvasively through a PCR screening with B-chromosome specific marker on DNA isolated from corresponding endosperms according to Karafiátová et al. [18]. Three biological replicates for both +B and 0B samples were collected for each developmental stage (early/mid/late)(Fig.2).

#### RNA extraction and RNA sequencing

RNA was isolated using Monarch Total RNA Miniprep Kit (NEB) following the protocol for samples tough to lyse. Samples were eluted into 30 μl of elution buffer and RNA quality and concentration was measured using RNA 6000 Pico or Nano kit (Agilent) on 2100 Bioanalyzer Instrument (Agilent)(Additional file 4). The sequencing libraries were created using NEBNext Single Cell/Low Input RNA Library Prep Kit for Illumina. Libraries were sequenced in 2×150 bp pair-end mode on the Illumina NovaSeq 6000 instrument (Illumina, Inc., San Diego, California).

#### Read mapping and differential expression analysis

The quality of raw RNA-seq data was assessed through fastQC, with subsequent removal of poor-quality reads based on quality (-q 30) and read length (-l 50) criteria using fastp [86]. The mapping of the RNA-seq reads to the reference genome sequence and annotation was carried out using STAR v2.7.11b [105]. Transcript-level quantification was done using RSEM v1.3.3 [106] to obtain raw counts. R package tximport was used to transform raw counts to normalized transcripts per million (TPM) level of expression. Differential expression was analyzed for all stages by the DeSEQ2 R package [107] with 0B samples as a base group to discover upregulated (log2 ≥ 2, p-value ≤ 0.05) and downregulated genes (log2 ≤ -2, p-value ≤ 0.05). Upregulated genes were matched against identified B-contigs to find B-located genes.

#### Functional annotation and GO-enrichment analysis

Functional annotation of all B located genes was performed using Blast2GO Diamond Search [108]. For all genes with a hit, GO term was matched from InterPro database. Next, gene set enrichment analysis (GSEA) was performed using topGO library [109] for all upregulated B-located genes for each stage separately. Importantly, only B-located genes were used as the reference gene set for enrichment analysis to avoid bias caused by overrepresentation of some GO terms among B chromosome-encoded genes comapred to full genome set. Significantly enriched terms were identified based on topGO networks, considering all terminal nodes with a p-value ≤ 0.05. (Additional file 7).

#### Laser-Capture Microdissection (LCM), RNA Extraction and mRNA amplification

Sample processing has been performed as described in Brandt et al. [110]. Briefly, frozen embryos at the mid-stage were glued onto sample plates by O.C.T medium. Cryo-sections (16 µm thickness) of +B and B0 embryos obtained as previously outlined [29], were mounted onto RNase-free PEN membrane slides (MMI, Eching, Germany) and stored in a freezer until use. To remove humidity, the slides were kept for 30 minutes in a desiccator before usage. A specific region, previously identified for B chromosome elimination (between plumule and rootwas isolated utilizing the MMI Cell Cut system (see Additional file 1: Fig. 2, 3). Usually, 60 dissected embryonic regions from around 15 individual embryos were pooled for one biological replicate. Three biological replicates were prepared for both +B and 0B samples. RNA was extracted using the Absolutely RNA Nanoprep Kit (Agilent) and mRNA was amplified by one round of T7-based mRNA amplification using the MessageAmp aRNA Kit (Invitrogen) to generate antisense RNA (aRNA).The quality of the isolated RNA was assessed using the Nano kit (Agilent) on the 2100 Bioanalyzer Instrument (Agilent) (Additional file 4). Library preparation, Illumina sequencing, differential gene expresion and GO-enrichment pipeline followed the same workflow as for analysis of whole embryo transcriptomes.

#### Comparison of A- and B-chromosomal variants of candidate genes

Upregulated, B chromosome encoded genes corresponding at least one of selected significantly enriched GO terms were further characterized (Additional file 3). BLASTP was used to find corresponding copies localized on a standard set of A chromosomes. The similarity between A- and B-chromosomal variants was retrieved from BLASTP results. In adition, pairwise aligments were produced using EMBOSS Needle Pairwise Sequence Alignment [39]. To validate the unique B chromosome gene variants and to further support the identification of B-contigs through k-mer analysis, PCR was employed for selected candidate genes - namely *CENH3*, *CENP-C*, *SMC3*, *APC15*, *CYCB1*, and *NAA50*. Initially, DNA sequences of A and B chromosome variants were aligned using EMBOSS Needle Pairwise Sequence Alignment [39] to identify unique sequences specific for B-specific variants. Next, primers targeting these unique sequencies were designed using primer3 [103](Additional file 1: Table S3). PCR reactions were prepared with 20 ng of genomic DNA from +B or 0B individuals, 1x Standard Taq Reaction Buffer, 0.2 mM dNTPs, 0.5U Taq DNA polymerase (New England Biolabs, Ipswich, MA, USA), 0.5 µM of each primer pair, and distilled water to a final volume of 25 µl. The PCR conditions were as follows: initial denaturation 94 °C/3 min; followed by 30 cycles of 94 °C/30 s; annealing at 55-60 °C/30 s; extension 72 °C/45 s; with a final extension step at 72 °C /10 min (Additional file 1: Table S3).

#### Protein structure prediction and interaction

Different CENH3 proteins and the CENH3-CENP-C nucleosome complex were predicted using the AlphaFold3 server [111]. The modelled subunits of the CENH3-CENP-C nucleosome complex were CENP-C (AA 692-764), CENH3 (AA 62-156), H2A (AA 17-135), H2B (AA 57-148) and H4 (AA 25-103). To identify the interaction interface between CENH3-CENP-C, the PDBePISA was used with default settings [112]. The protein structures were visualized using the UCSF ChimeraX program [113]. The figures were assembled using the Inkscape program (https://inkscape.org/).

#### Identification of CENH3 B chromosome variants in Aegilops speltoides and Zea mays

Publicly available FASTQ data from anthers (undergoing the first pollen mitosis) of *Aegilops speltoides* with 3 B chromosomes (Genotype K2 from Tartus, Syria) were used to *de novo* assemble the transcriptome with Trinity (v2.4.0, default parameters) [34; 114]. TransDecoder (v5.5.0) [115] was used to annotate coding regions within transcripts. Blastp (v2.10.1, default parameters) was used to align the wheat CENH3 to the translated transcriptome of *A. speltoides* + B. CENH3 sequences of *Zea mays* were retrieved from annotation in Blavet et al. [33].

#### Phylogenetic analysis

Protein sequences of CENH3 protein (Additional file 10) were downloaded from two PANTHER subfamilies PTHR11426:SF223 and PTHR11426:SF277 [116] and aligned using MAFFTv7.029 (--globalpair --maxiterate 1000) [117]. The multiple alignment was graphicaly displayed in SeaView v5.0.5 [118], Non-homologous sequence region of *Brachypodium distachyon*, *Brassica campestris*, and *Gycine max*, situated upstreem the CENH3 protein sequence, were removed from the alignment prior to phylogenetic analysis. The phylogenetic tree was constructed with PhyML 3.1 [119], using LG model [120] implemented in SeaView. Approximate likelihood ratio tests [121] were performed to assess branch support. Final phylogenetic tree was depicted and edited with FigTree (http://tree.bio.ed.ac.uk/software/figtree/).

## Supplementary Information

Additional file 1: Supplementary Document: Fig. S1-S7, Table S1-S3 (file contains separate channels of FISH experiments (Fig. S1), pipeline of transcriptomics experiments (Fig. S2), LCM of sections dataset (Fig. S3), additional PCA clustering (Fig. S4) and GSEA analysis (Fig. S5) of sections dataset, electrophoretogram of the B -specific amplicons (Fig. S6), expression levels of B-localized genes (Fig. S7), sequence content in *S. purpureosericeum* (Table S1), primers for B-specific repeat clusters (Table S2) and candidates screening (Table S3)).

Additional file 2: Supplementary Table (file contains A chromosomes or B chromosome contigs length, +B and 0B counts in k-mer analysis and A/B contig identities).

Additional file 3: Supplementary Table (file contains list of B-contigs located genes together with their gene ontologies).

Additional file 4: Supplementary Table (file contains information about the samples used for RNA sequencing).

Additional file 5: Supplementary Table (file contains differential expression data).

Additional file 6: Supplementary Table (file contains GSEA results in whole embryo groups and sections RNA-seq data).

Additional file 7: Supplementary document (file contains hierarchy trees of all significant GO terms).

Additional file 8: Supplementary document (file contains alignments of A/B chromosome variants of candidate genes).

Additional file 9: Supplementary Table (file contains PISA analysis results).

Additional file 10: Supplementary Table: Fig. S1 (file contains sequencies of CENH3 proteins).

Additional file 11: Supplementary document (file contains full figure of phylogenic tree of CENH3 proteins).

Additional file 12: Supplementary document (file contains protocol for HMW isolation for ONT sequencing).

## Funding

This work was supported by The Czech Science Foundation project no. 22-02108S (awarded to MK) and 23-04887S (awarded to JB); by The Czech Academy of Sciences project no. DAAD-22-02 (awarded to MK). AH was supported by the DFG (HO1779/30-1 and 30-2; HO1779/34-1; HO 1779/26-2), JT was supported by DFH (TH 1876/5-2).

## Authors’ contributions

Experimental design: JB and MK. Whole genome seguencing: RS, TB, LH. Embryo transcriptomics: TB, LH, KH, GK. Cytogenetics and microscopy: TB, MK, LH. LCM-based isolation of embryo regions and sample processing for RNA-seq: LH, TB, JT. Protein modeling: RP, II and JB. Phylogeny: EH. Manuscript preparation: TB, JB, MK and AH. All authors discussed the results and commented on the manuscript.

## Acknowledgements

We thank the International Crop Research Institute for the Semi-Arid Tropics (ICRISAT) for providing the original seed stock of *Sorghum purpureosericeum*. We thank Jitka Weisserová and Magdalena Stejskalová for their technical assistance. Computational resources were provided by the e-INFRA CZ project (ID:90254), supported by the Ministry of Education, Youth and Sports of the Czech Republic and by the ELIXIR-CZ project (ID:90255), part of the international ELIXIR infrastructure.

## Declarations

### Ethics approval and consent to participate

Not aplicable.

### Consent for publication

Not aplicable.

### Availability of data and materials

Raw sequence reads are available from the European Nucleotide Archive (ENA) under accession PRJEB83623 (whole genome sequencing short reads, Nanopore reads) and PRJEB83179 (RNA-seq). The GCA assembly ID of the +B genome assembly is GCA_964647645.

### Competing interests

The authors declare no competing interests.

### Authors’ information (optional)

Institute of Experimental Botany of the Czech Academy of Sciences, Centre of Plant Structural and Functional Genomics, Šlechtitelů 31, 779 00 Olomouc, Czech Republic ^2^ Department of Cell Biology and Genetics, Faculty of Science, Palacky University, Šlechtitelů 27, 779 00 Olomouc, Czech Republic ^3^ Leibniz Institute of Plant Genetics and Crop Plant Research (IPK), 06466 Seeland OT Gatersleben, Germany ^4^ Institute of Experimental Botany of the Czech Academy of Sciences, Laboratory of Integrative Structural Biology, Rozvojová 263,165 00 Praha 6 - Lysolaje, Czech Republic

## Notes

https://zenodo.org/records/14197239

### Competing Interest Statement

The authors have declared no competing interest.

