## Additional file 1 for "Sorghum embryos undergoing B chromosome elimination express B-variants of mitotic-related genes"

Additional file 1: Supplementary Document: Fig. S1-S7, Table S1-S3.

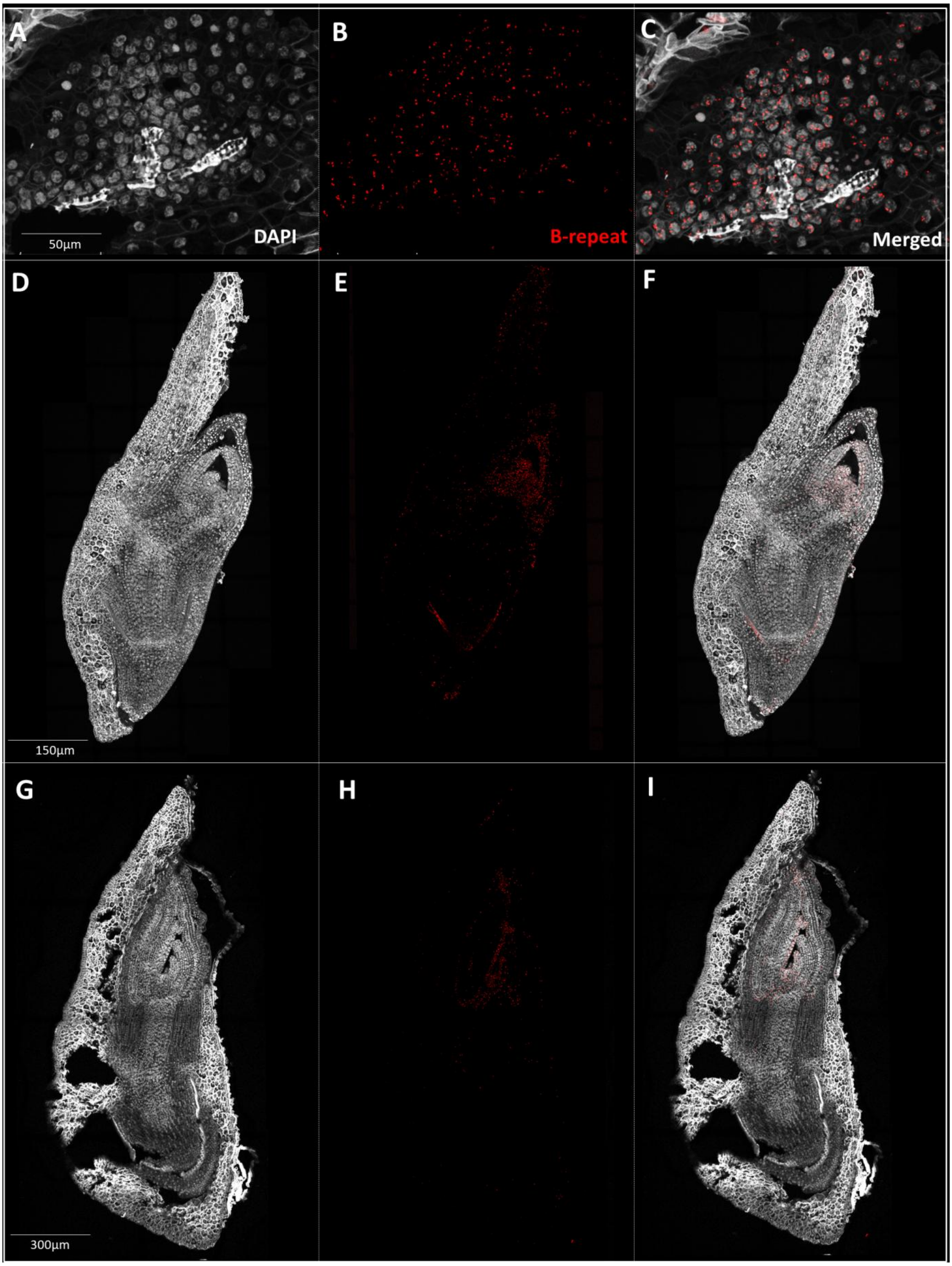

**Fig. S1**: In situ visualization of the B chromosome on embryo cryo-section during studied stages of embryonal development in wild sorghum. A-C – early stage; D-F – mid-stage; G-I – late stage. B-positive cells were detected using B-specific probe (red). DNA is counterstained with DAPI (grey*).*

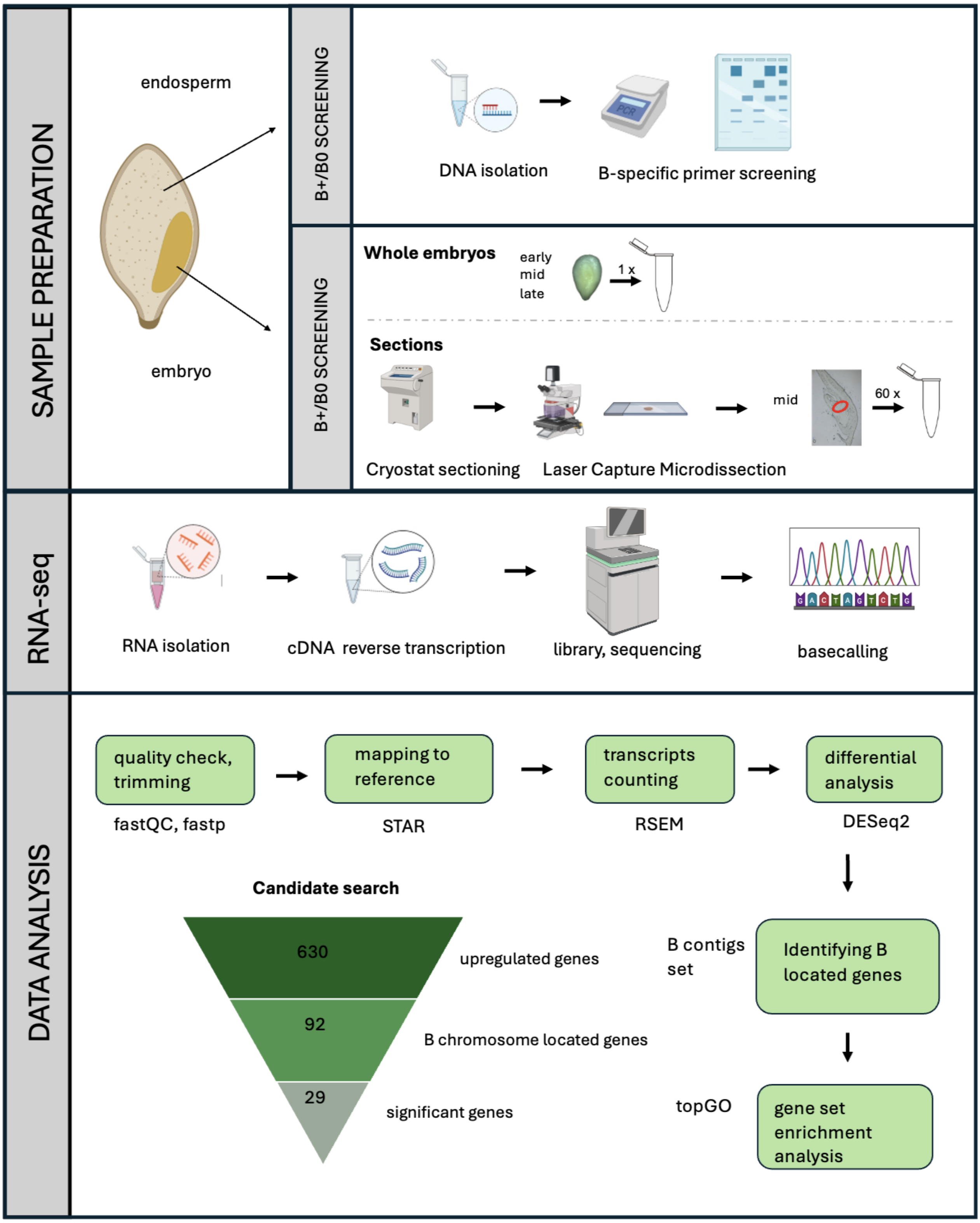


**Fig. S2:** Pipeline of transcriptomic experiments. Created with BioRender.com

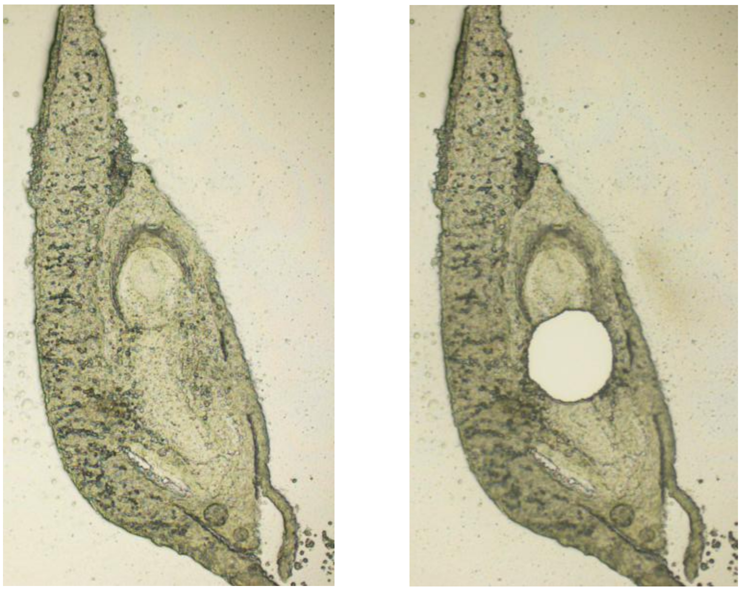

**Fig. S3**: Laser Capture Microdissection of embryonic region undergoing B chromosome elimination. A – Mid-stage embryo section before dissection, B – Mid-stage embryo section after dissection.

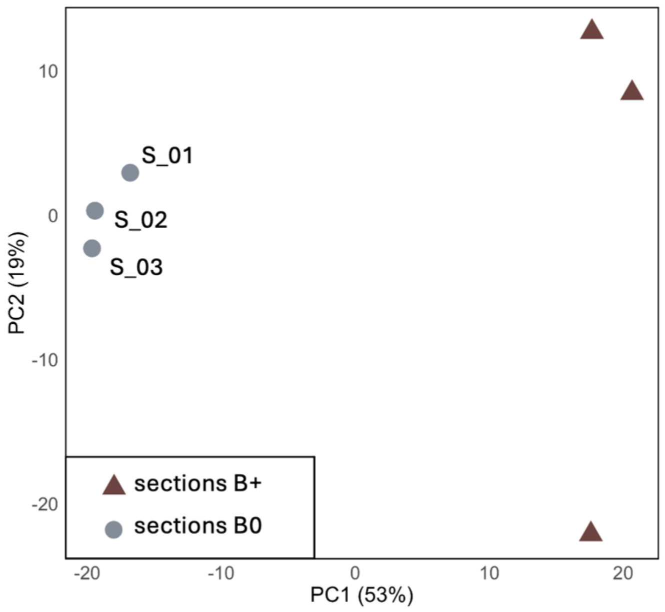


**Fig. S4**: Principal Component Analysis (PCA) variance of 6 section RNA-seq samples.

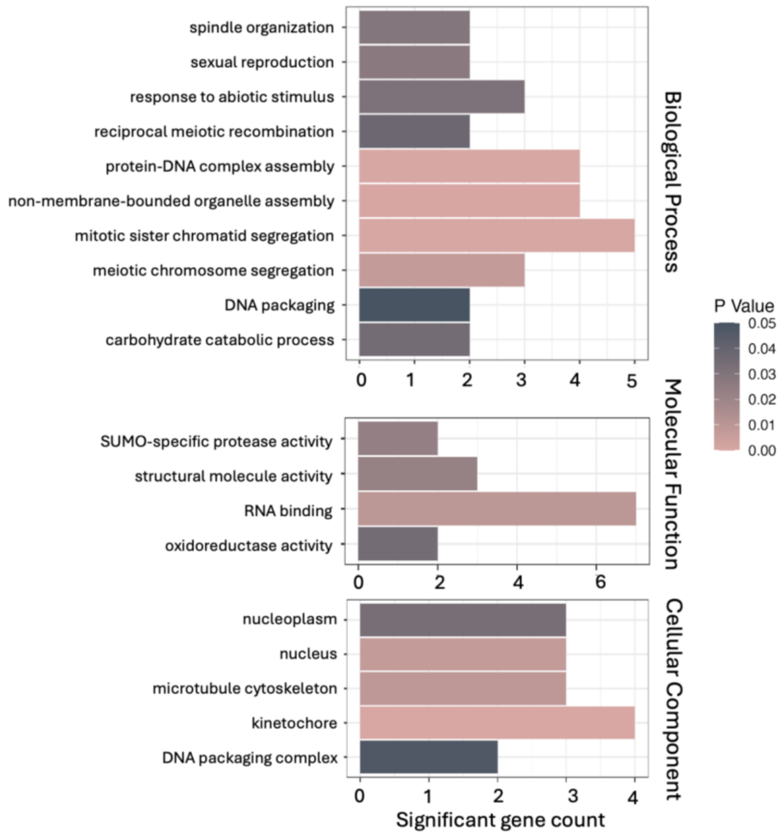


**Fig. S5**: Gene set enrichment analysis of upregulated B located genes in the sections dataset. The barplot visualizes the enrichment of all significantly enriched gene ontology (GO) terms (p-value < 0.05). Bar length represent number of identified upregulated genes; colour correspond to significance level.

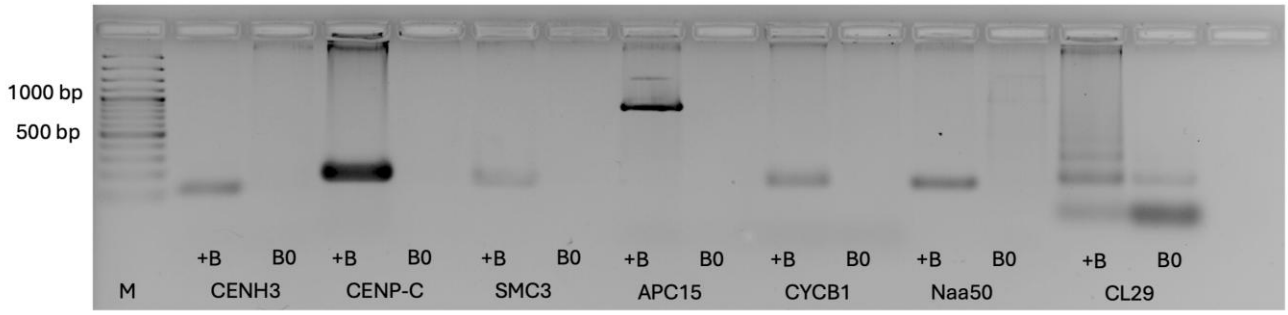


**Fig. S6**: B-specific PCR amplicons for candidates using B+ and B0 DNA templates. Due to impossibility to design B-specific primer for gene sequence, downstream sequence of utg10093 is included. GeneRuler 100 bp DNA Ladder was used for size reference. As a negative control, centromeric repetition CL29 (Karafiátová et al., 2024) was used.

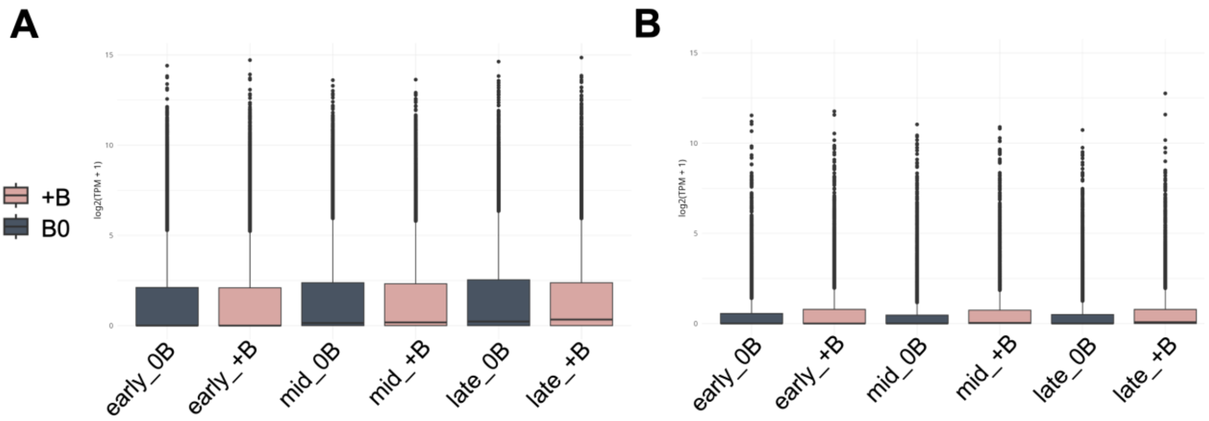


**Fig. S7**: Boxplot showing the log (TPM+1) transformed expression levels of all genes (**A**) and B-localized genes (**B**).

Table S1: Sequence content in *S. purpureosericeum* of A contigs and 420 Mb of B contigs

| | *A contigs (bp)* | *B contigs (bp)* |
|---|---|---|
| *Genes* | 93,728,661 | 5,531,721 |
| *Tandem repeats* | 678,413 | 4,322,828 |
| *Ty1/copia* | 205,590,413 | 4,2254,642 |
| *Ty3/gypsy* | 443,738,910 | 91,104,319 |
| *Other LTRs* | 590,643,748 | 102,165,677 |
| *non-LTR* | 31,812,783 | 4,878,781 |
| *DNA transposons* | 46,689,765 | 6,605,022 |
| *Other sequences* | 992,953,798 | 163,042,214 |

Table S2: Primers for B-specific repeat clusters

| *B-repeat* | *primer sequencies* | *annealing t (°C)* | *product size (bp)* |
|---|---|---|---|
| *CL137_F*<br>*CL137_R* | CGAGAGCCAACGTTCATTTT<br>TTAGCAATGGGATGGCTCTT | 60 | 1125 |
| *CL166_F*<br>*CL166_R* | CCTGTATCAAAATGTCTCCATGTC<br>ACTGCGTCCTAAAACGGTGA | 60 | 344 |
| *CL193_F*<br>*CL193_R* | CGAGAAAATGGAGCACAACC<br>AAGGGATGGTGCACTGGA | 60 | 242 |
| *CL220_F*<br>*CL220_R* | AAAACAATGGTCGGATGGAA<br>AAATGTAAGCTGCCAATTCTGA | 60 | 1561 |

Table S3: B chromosome specific primers for candidate genes

| *candidate* | *primer sequencies* | *annealing t (°C)* | *product size (bp)* |
|---|---|---|---|
| *CENH3_F*<br>*CENH3_R* | CTCTCCTTTCGTTTCGCCAT<br>AGCTTCTTCTTGGGCTTCTG | 56 | 152 |
| *CENP-C_F*<br>*CENP-C_R* | GCTTTTGTTGTGCATGACCA<br>GAGAACTCAAGCCATTCACACT | 55 | 198 |
| *SMC3_F*<br>*SMC3_R* | AGTGACTGGTTTGCTGGTTG<br>TTGTTGTTCCCAAGGTAGGTT | 55 | 245 |
| *Naa50_F*<br>*Naa50_R* | TGCGGGATAAGAACGAGCTC<br>AACAGAGAAGGGGAAGACGG | 59 | 150 |
| *APC15_F*<br>*APC15_R* | CTGTGATGGATCTCGTTGCG<br>ACTGCCTTCCAAGCCTAGAG | 60 | 887 |
| *CYC1B_F*<br>*CYC1B_R* | TAATGGAGGGTGGGGATT GC<br>TCATACAAGACTGACCTGGCT | 58 | 153 |
