## Additional file 7 for "Sorghum embryos undergoing B chromosome elimination express B-variants of mitotic-related genes"

Additional File 7: TopGO-directed hierarchical trees of all Gene Ontology (GO) terms identified through the GSEA analysis. Significant terms (p > 0.05) are depicted as boxes within the plot, with deeper colors representing greater levels of significance.

BP terms in early whole embryos

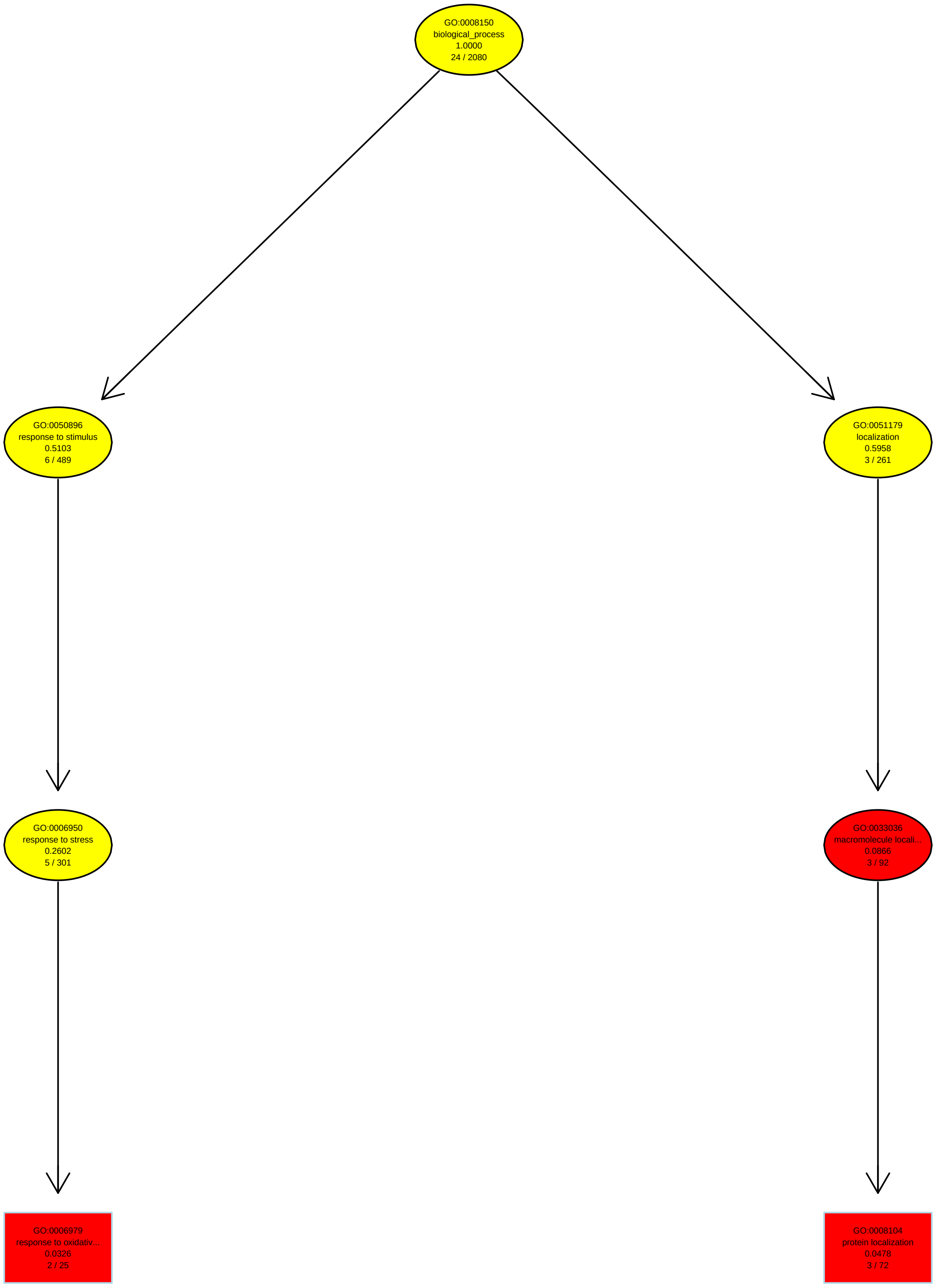

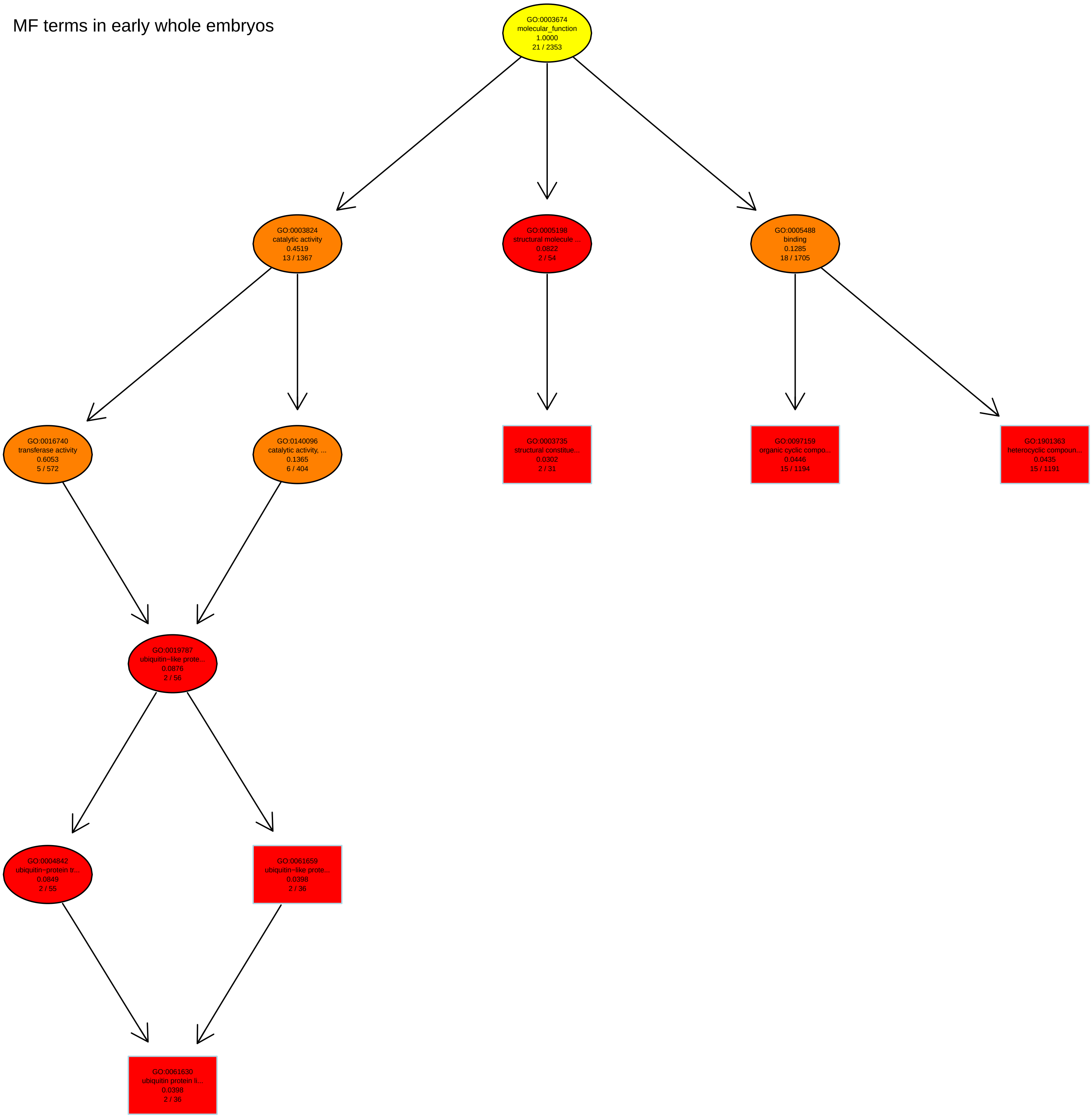
MF terms in early whole embryos
GO:0003674
molecular_function
1.0000
21 / 2353
GO:0003824
catalytic activity
0.4519
13 / 1367
GO:0005198
structural molecule ...
0.0822
2 / 54
GO:0005488
binding
0.1285
18 / 1705
GO:0016740
transferase activity
0.6053
5 / 572
GO:0140096
catalytic activity, ...
0.1365
6 / 404
GO:0003735
structural constitue...
0.0302
2 / 31
GO:0097159
organic cyclic compo...
0.0446
15 / 1194
GO:1901363
heterocyclic compoun...
0.0435
15 / 1191
GO:0019787
ubiquitin−like prote...
0.0876
2 / 56
GO:0004842
ubiquitin−protein tr...
0.0849
2 / 55
GO:0061659
ubiquitin−like prote...
0.0398
2 / 36
GO:0061630
ubiquitin protein li...
0.0398
2 / 36

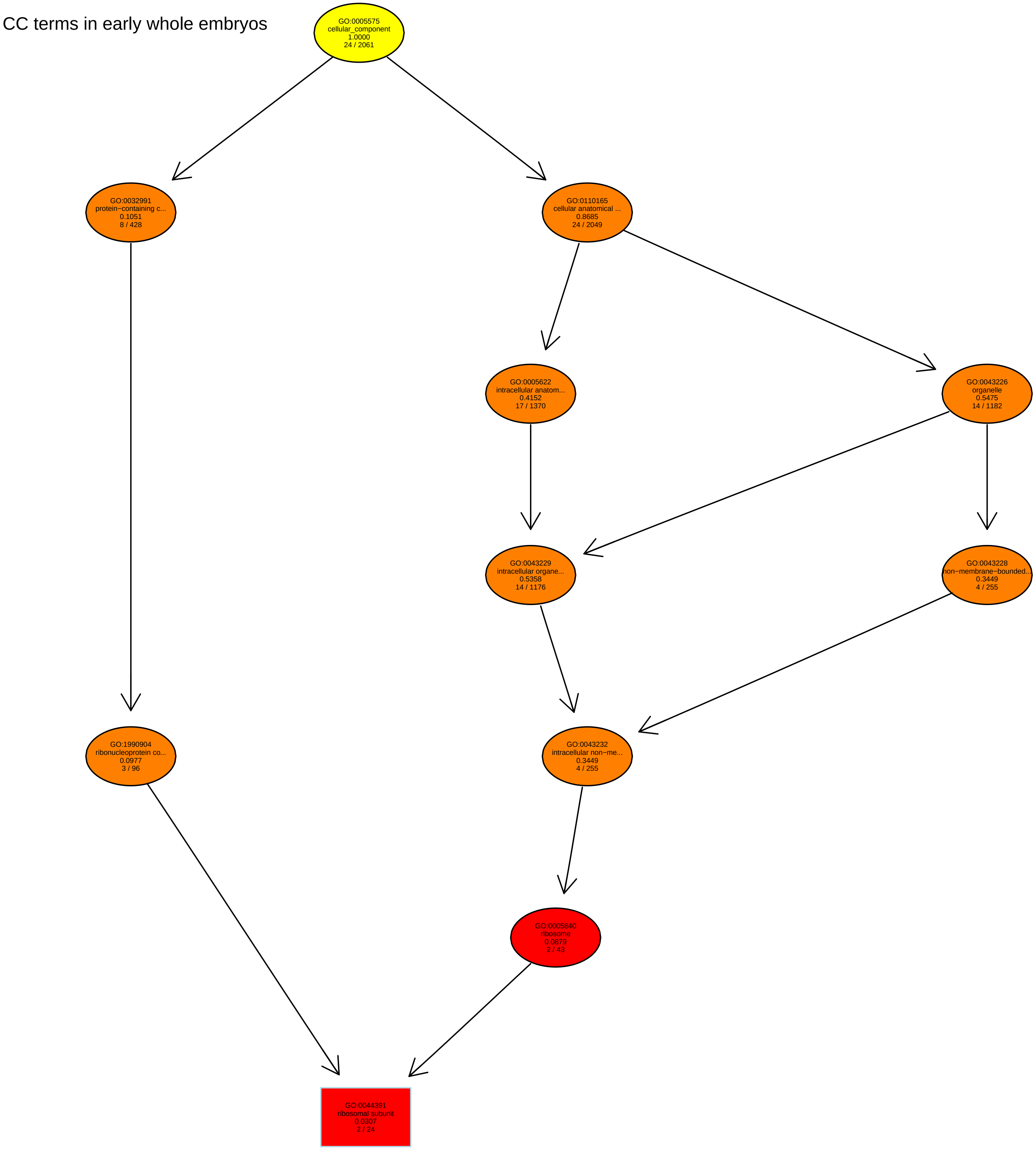
CC terms in early whole embryos
GO:0005575
cellular_component
1.0000
24 / 2061
GO:0032991
protein−containing c...
0.1051
8 / 428
GO:0110165
cellular anatomical ...
0.8685
24 / 2049
GO:0005622
intracellular anatom...
0.4152
17 / 1370
GO:0043226
organelle
0.5475
14 / 1182
GO:0043229
intracellular organe...
0.5358
14 / 1176
GO:0043228
non−membrane−bounded...
0.3449
4 / 255
GO:1990904
ribonucleoprotein co...
0.0977
3 / 96
GO:0043232
intracellular non−me...
0.3449
4 / 255
GO:0005840
ribosome
0.0879
2 / 43
GO:0044391
ribosomal subunit
0.0307
2 / 24

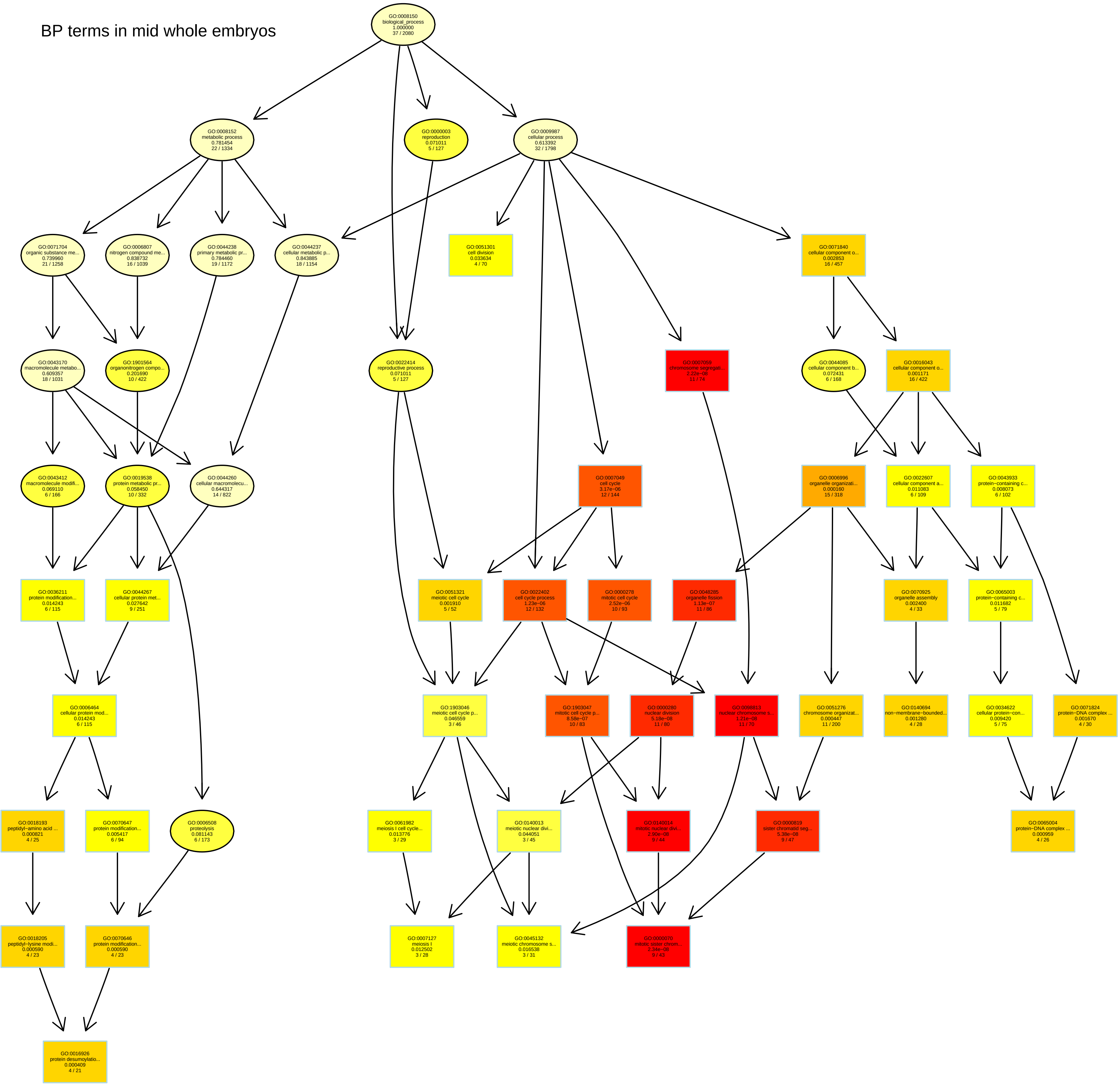
BP terms in mid whole embryos
GO:0008150 biological_process 1.000000 37 / 2080
GO:0008152 metabolic process 0.781454 22 / 1334
GO:0000003 reproduction 0.071011 5 / 127
GO:0009987 cellular process 0.613392 32 / 1798
GO:0071704 organic substance me... 0.739960 21 / 1258
GO:0006807 nitrogen compound me... 0.838732 16 / 1039
GO:0044238 primary metabolic pr... 0.784460 19 / 1172
GO:0044237 cellular metabolic p... 0.843885 18 / 1154
GO:0051301 cell division 0.033634 4 / 70
GO:0071840 cellular component o... 0.002853 16 / 457
GO:0043170 macromolecule metabo... 0.609357 18 / 1031
GO:1901564 organonitrogen compo... 0.201690 10 / 422
GO:0022414 reproductive process 0.071011 5 / 127
GO:0007059 chromosome segregati... 2.22e−08 11 / 74
GO:0044085 cellular component b... 0.072431 6 / 168
GO:0016043 cellular component o... 0.001171 16 / 422
GO:0043412 macromolecule modifi... 0.069110 6 / 166
GO:0019538 protein metabolic pr... 0.058450 10 / 332
GO:0044260 cellular macromolecu... 0.644317 14 / 822
GO:0007049 cell cycle 3.17e−06 12 / 144
GO:0006996 organelle organizati... 0.000160 15 / 318
GO:0022607 cellular component a... 0.011083 6 / 109
GO:0043933 protein−containing c... 0.008073 6 / 102
GO:0036211 protein modification... 0.014243 6 / 115
GO:0044267 cellular protein met... 0.027642 9 / 251
GO:0051321 meiotic cell cycle 0.001910 5 / 52
GO:0022402 cell cycle process 1.23e−06 12 / 132
GO:0000278 mitotic cell cycle 2.52e−06 10 / 93
GO:0048285 organelle fission 1.13e−07 11 / 86
GO:0070925 organelle assembly 0.002400 4 / 33
GO:0065003 protein−containing c... 0.011682 5 / 79
GO:0006464 cellular protein mod... 0.014243 6 / 115
GO:1903046 meiotic cell cycle p... 0.046559 3 / 46
GO:1903047 mitotic cell cycle p... 8.58e−07 10 / 83
GO:0000280 nuclear division 5.18e−08 11 / 80
GO:0098813 nuclear chromosome s... 1.21e−08 11 / 70
GO:0051276 chromosome organizat... 0.000447 11 / 200
GO:0140694 non−membrane−bounded... 0.001280 4 / 28
GO:0034622 cellular protein−con... 0.009420 5 / 75
GO:0071824 protein−DNA complex ... 0.001670 4 / 30
GO:0018193 peptidyl−amino acid ... 0.000821 4 / 25
GO:0070647 protein modification... 0.005417 6 / 94
GO:0006508 proteolysis 0.081143 6 / 173
GO:0061982 meiosis I cell cycle... 0.013776 3 / 29
GO:0140013 meiotic nuclear divi... 0.044051 3 / 45
GO:0140014 mitotic nuclear divi... 2.90e−08 9 / 44
GO:0000819 sister chromatid seg... 5.38e−08 9 / 47
GO:0065004 protein−DNA complex ... 0.000959 4 / 26
GO:0018205 peptidyl−lysine modi... 0.000590 4 / 23
GO:0070646 protein modification... 0.000590 4 / 23
GO:0007127 meiosis I 0.012502 3 / 28
GO:0045132 meiotic chromosome s... 0.016538 3 / 31
GO:0000070 mitotic sister chrom... 2.34e−08 9 / 43
GO:0016926 protein desumoylatio... 0.000409 4 / 21

MF terms in mid whole embryos

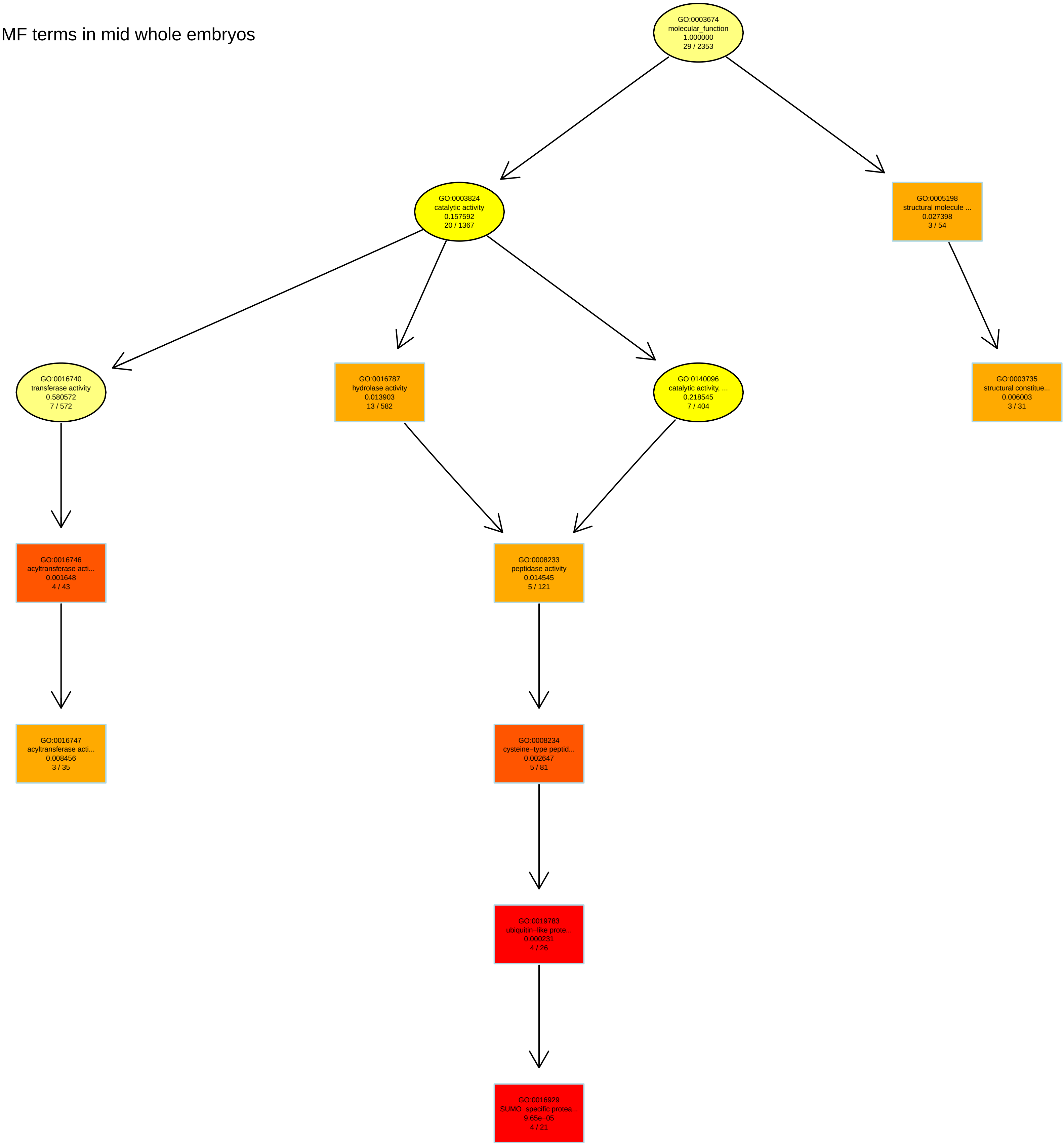

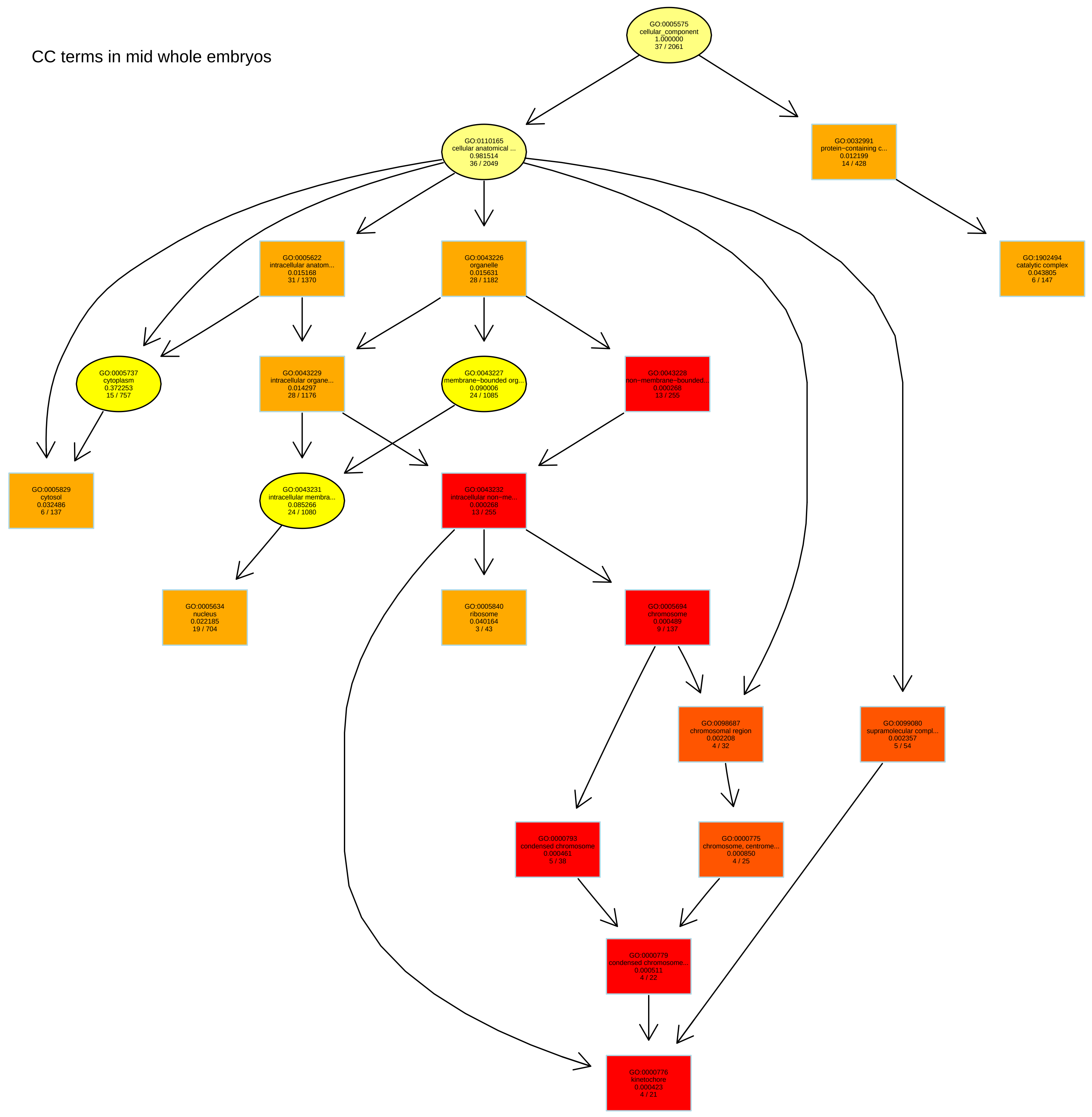
CC terms in mid whole embryos
GO:0005575
cellular_component
1.000000
37 / 2061
GO:0110165
cellular anatomical ...
0.981514
36 / 2049
GO:0032991
protein−containing c...
0.012199
14 / 428
GO:0005622
intracellular anatom...
0.015168
31 / 1370
GO:0043226
organelle
0.015631
28 / 1182
GO:1902494
catalytic complex
0.043805
6 / 147
GO:0005737
cytoplasm
0.372253
15 / 757
GO:0043229
intracellular organe...
0.014297
28 / 1176
GO:0043227
membrane−bounded org...
0.090006
24 / 1085
GO:0043228
non−membrane−bounded...
0.000268
13 / 255
GO:0005829
cytosol
0.032486
6 / 137
GO:0043231
intracellular membra...
0.085266
24 / 1080
GO:0043232
intracellular non−me...
0.000268
13 / 255
GO:0005634
nucleus
0.022185
19 / 704
GO:0005840
ribosome
0.040164
3 / 43
GO:0005694
chromosome
0.000489
9 / 137
GO:0098687
chromosomal region
0.002208
4 / 32
GO:0099080
supramolecular compl...
0.002357
5 / 54
GO:0000793
condensed chromosome
0.000461
5 / 38
GO:0000775
chromosome, centrome...
0.000850
4 / 25
GO:0000779
condensed chromosome...
0.000511
4 / 22
GO:0000776
kinetochore
0.000423
4 / 21

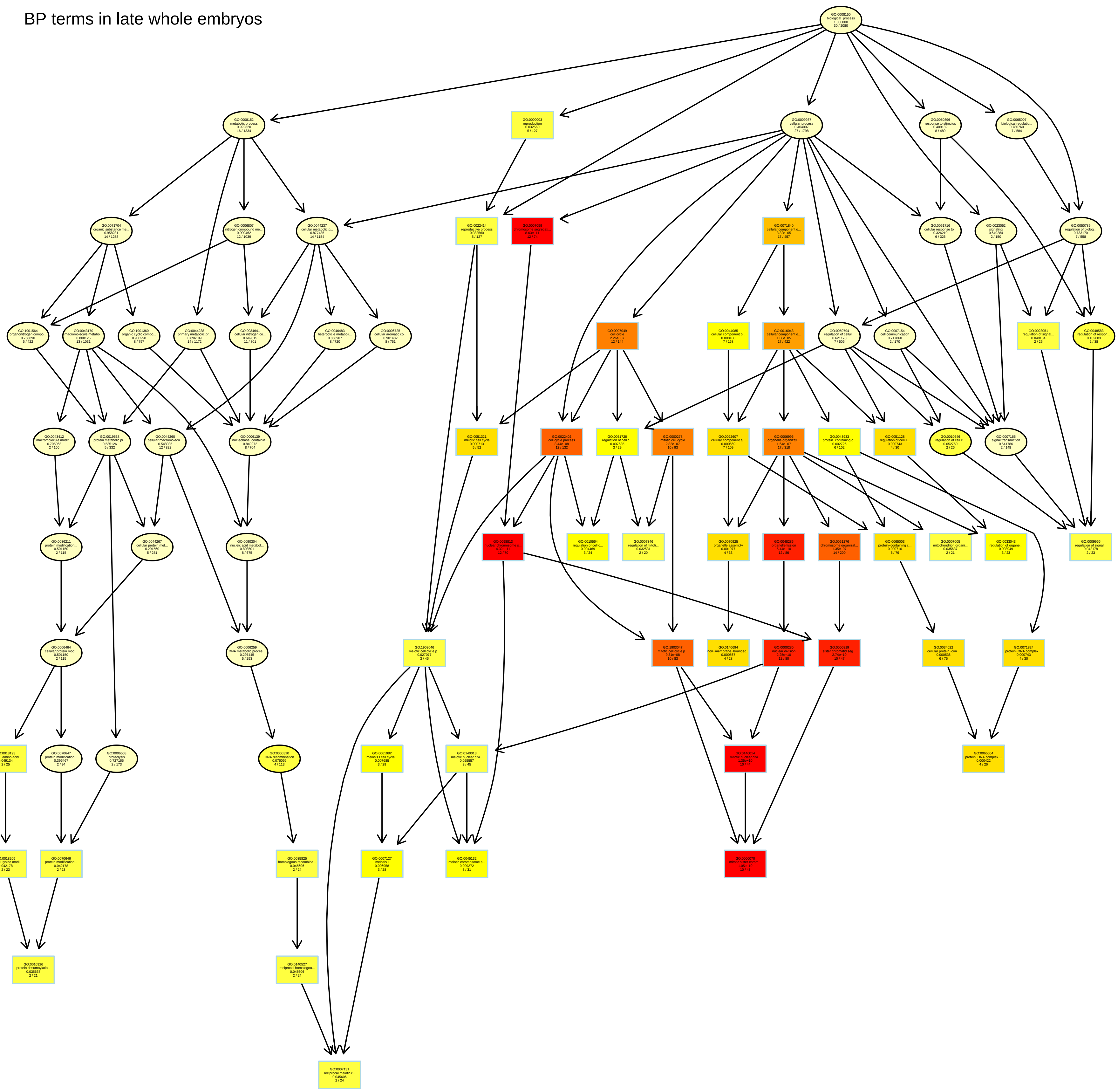
BP terms in late whole embryos

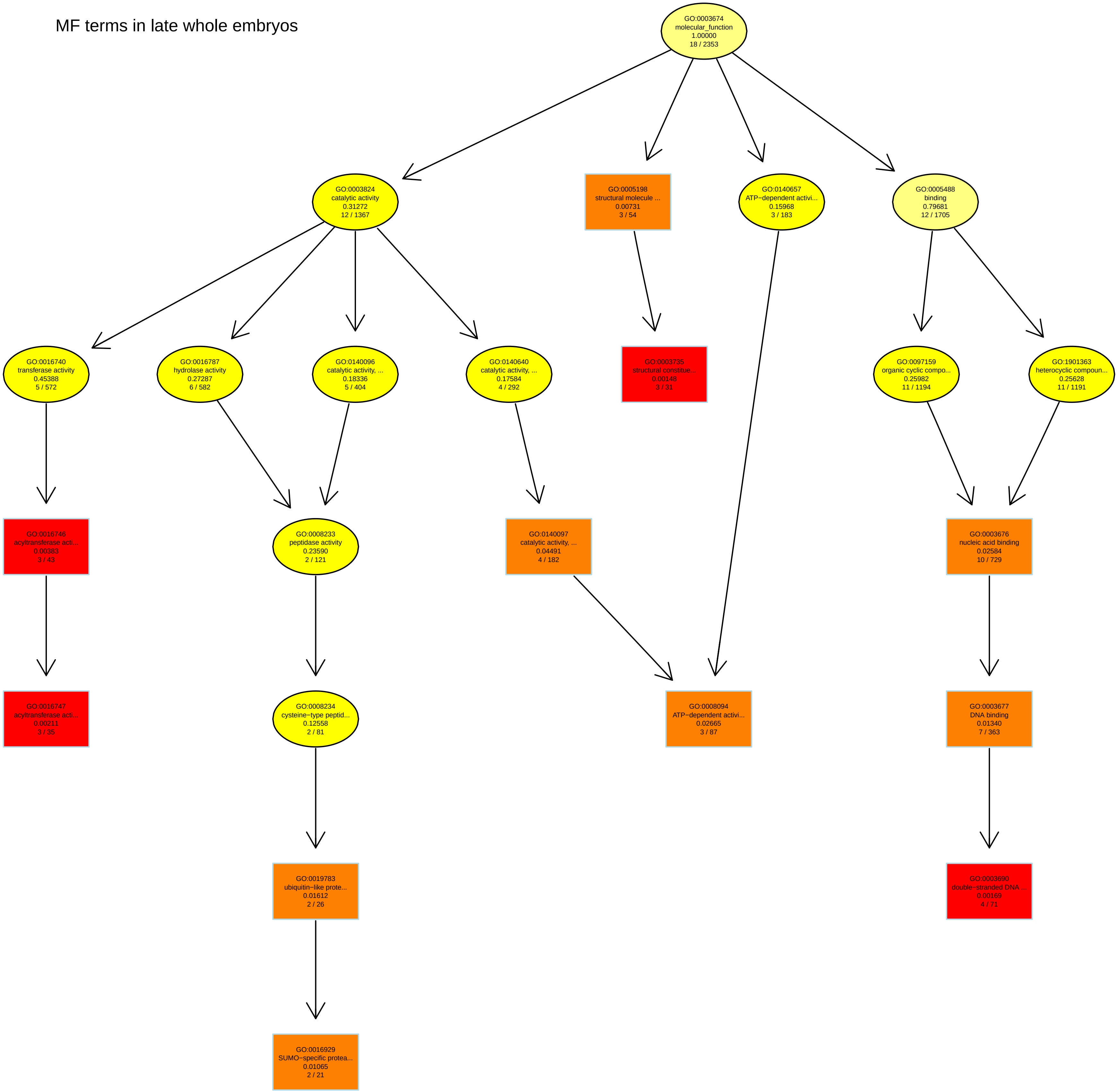
MF terms in late whole embryos
GO:0003674 molecular_function 1.00000 18 / 2353
GO:0003824 catalytic activity 0.31272 12 / 1367
GO:0005198 structural molecule ... 0.00731 3 / 54
GO:0140657 ATP−dependent activi... 0.15968 3 / 183
GO:0005488 binding 0.79681 12 / 1705
GO:0016740 transferase activity 0.45388 5 / 572
GO:0016787 hydrolase activity 0.27287 6 / 582
GO:0140096 catalytic activity, ... 0.18336 5 / 404
GO:0140640 catalytic activity, ... 0.17584 4 / 292
GO:0003735 structural constitue... 0.00148 3 / 31
GO:0097159 organic cyclic compo... 0.25982 11 / 1194
GO:1901363 heterocyclic compoun... 0.25628 11 / 1191
GO:0016746 acyltransferase acti... 0.00383 3 / 43
GO:0008233 peptidase activity 0.23590 2 / 121
GO:0140097 catalytic activity, ... 0.04491 4 / 182
GO:0003676 nucleic acid binding 0.02584 10 / 729
GO:0016747 acyltransferase acti... 0.00211 3 / 35
GO:0008234 cysteine−type peptid... 0.12558 2 / 81
GO:0008094 ATP−dependent activi... 0.02665 3 / 87
GO:0003677 DNA binding 0.01340 7 / 363
GO:0019783 ubiquitin−like prote... 0.01612 2 / 26
GO:0003690 double−stranded DNA ... 0.00169 4 / 71
GO:0016929 SUMO−specific protea... 0.01065 2 / 21

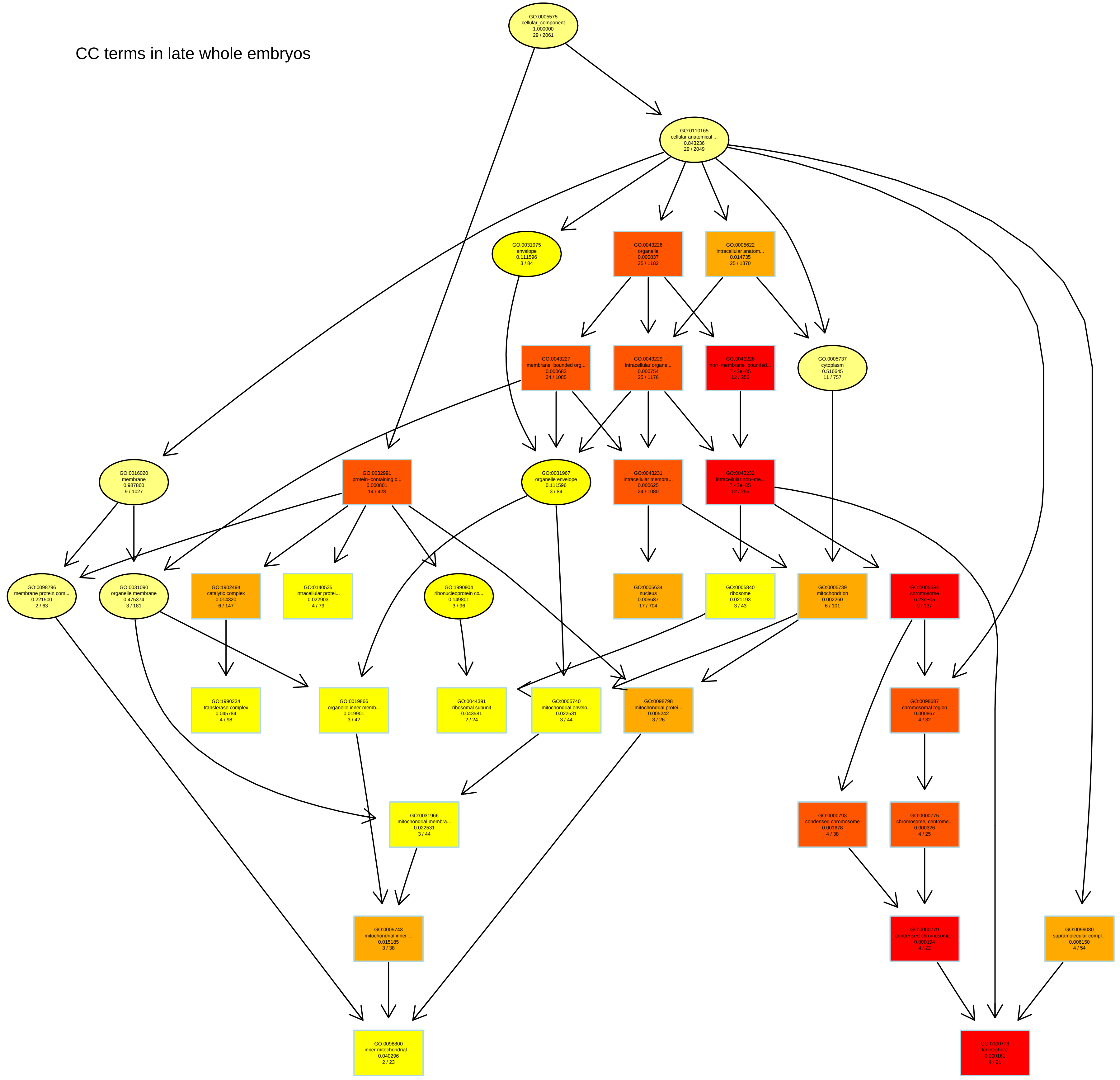
CC terms in late whole embryos
GO:0005575 cellular_component 1.000000 29 / 2061
GO:0110165 cellular anatomical ... 0.843236 29 / 2049
GO:0031975 envelope 0.111596 3 / 84
GO:0043226 organelle 0.000837 25 / 1182
GO:0005622 intracellular anatom... 0.014735 25 / 1370
GO:0043227 membrane−bounded org... 0.000683 24 / 1085
GO:0043229 intracellular organe... 0.000754 25 / 1176
GO:0043228 non−membrane−bounded... 7.43e−05 12 / 255
GO:0005737 cytoplasm 0.516645 11 / 757
GO:0016020 membrane 0.987860 9 / 1027
GO:0032991 protein−containing c... 0.000801 14 / 428
GO:0031967 organelle envelope 0.111596 3 / 84
GO:0043231 intracellular membra... 0.000625 24 / 1080
GO:0043232 intracellular non−me... 7.43e−05 12 / 255
GO:0098796 membrane protein com... 0.221500 2 / 63
GO:0031090 organelle membrane 0.475374 3 / 181
GO:1902494 catalytic complex 0.014320 6 / 147
GO:0140535 intracellular protei... 0.022903 4 / 79
GO:1990904 ribonucleoprotein co... 0.149801 3 / 96
GO:0005634 nucleus 0.005687 17 / 704
GO:0005840 ribosome 0.021193 3 / 43
GO:0005739 mitochondrion 0.002260 6 / 101
GO:0005694 chromosome 6.23e−05 9 / 137
GO:1990234 transferase complex 0.045784 4 / 98
GO:0019866 organelle inner memb... 0.019901 3 / 42
GO:0044391 ribosomal subunit 0.043581 2 / 24
GO:0005740 mitochondrial envelo... 0.022531 3 / 44
GO:0098798 mitochondrial protei... 0.005242 3 / 26
GO:0098687 chromosomal region 0.000867 4 / 32
GO:0031966 mitochondrial membra... 0.022531 3 / 44
GO:0000793 condensed chromosome 0.001678 4 / 38
GO:0000775 chromosome, centrome... 0.000326 4 / 25
GO:0005743 mitochondrial inner ... 0.015185 3 / 38
GO:0000779 condensed chromosome... 0.000194 4 / 22
GO:0099080 supramolecular compl... 0.006150 4 / 54
GO:0098800 inner mitochondrial ... 0.040296 2 / 23
GO:0000776 kinetochore 0.000161 4 / 21

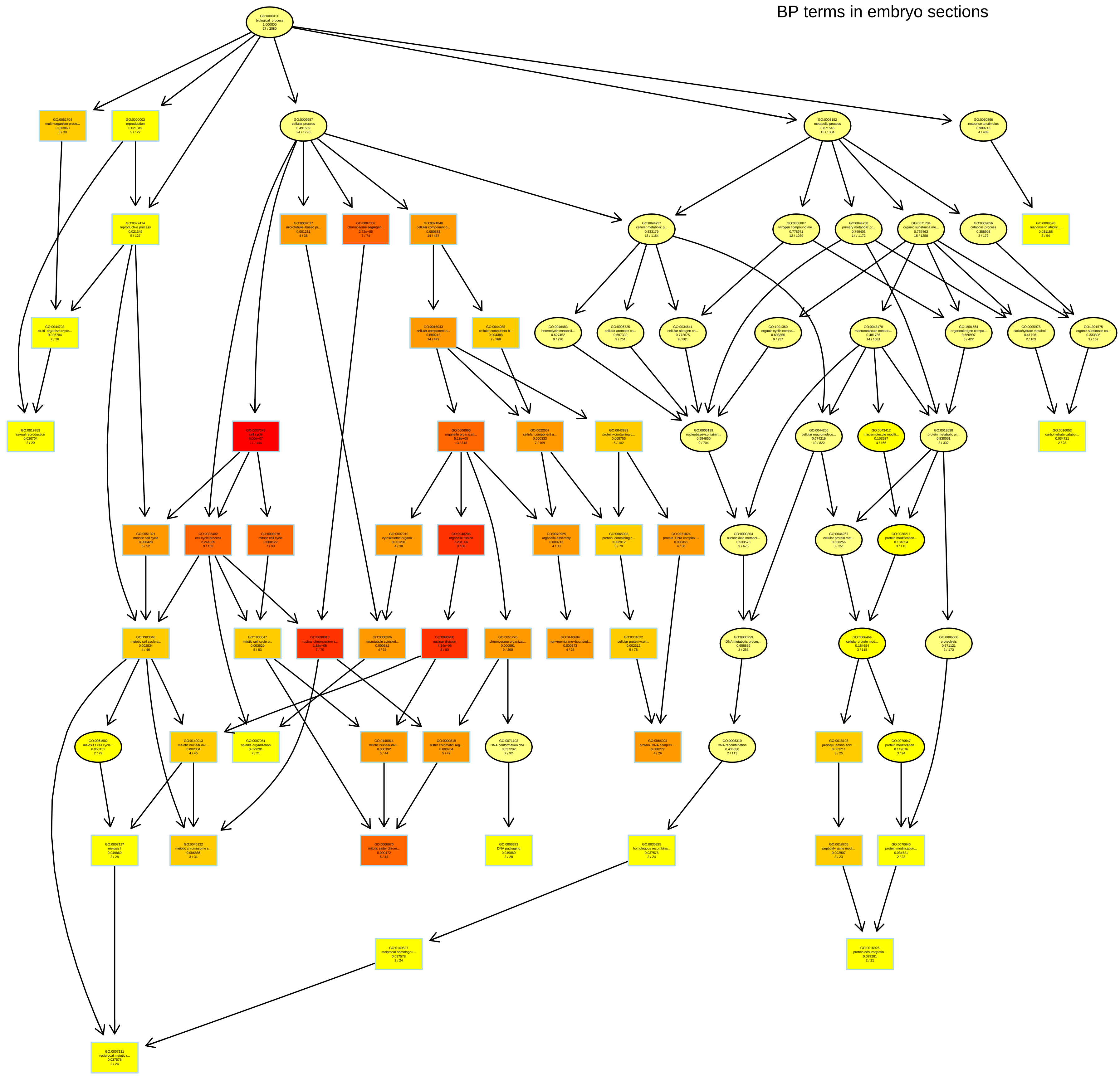
BP terms in embryo sections

MF terms in embryo sections

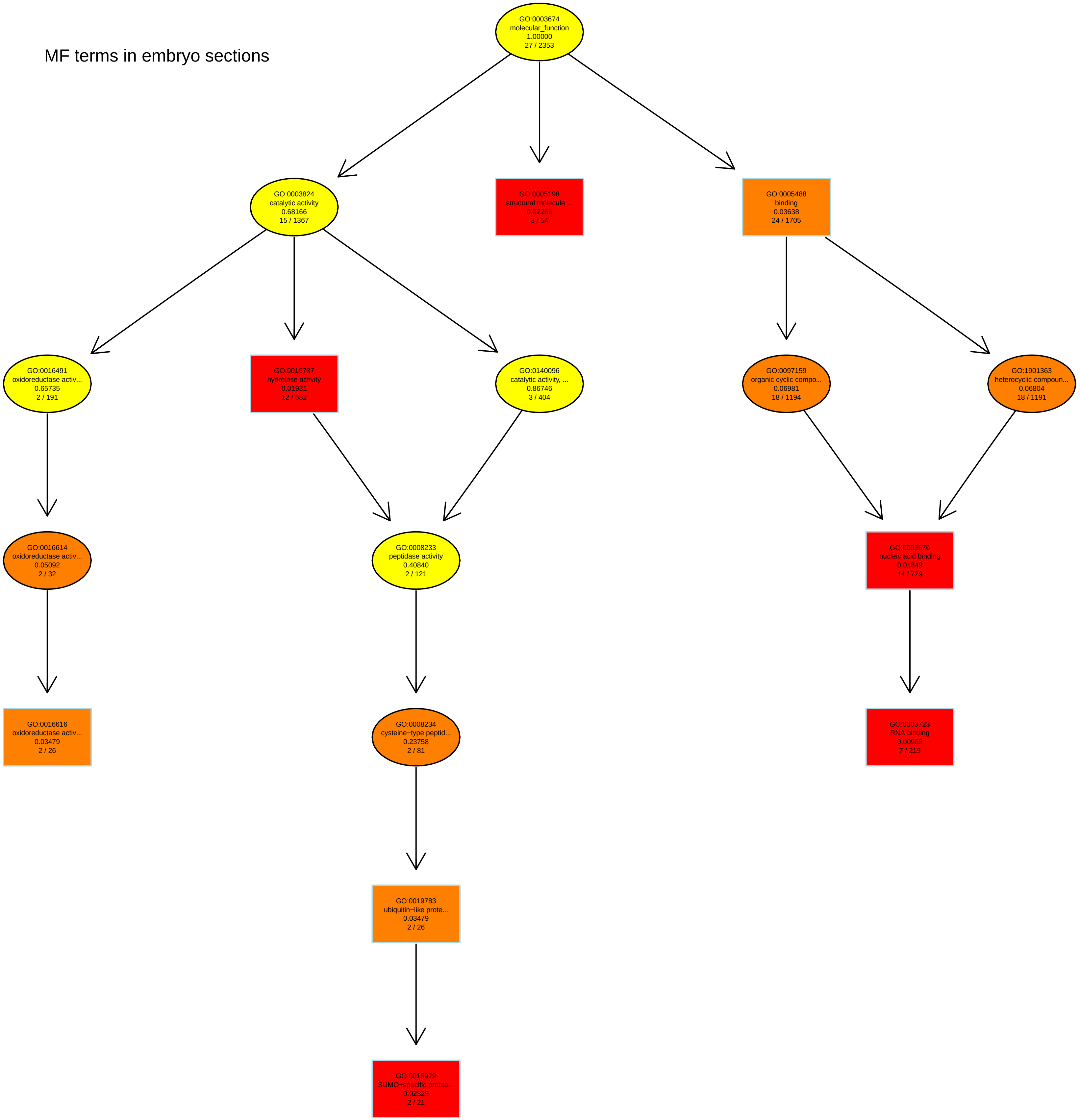

CC terms in embryo sections

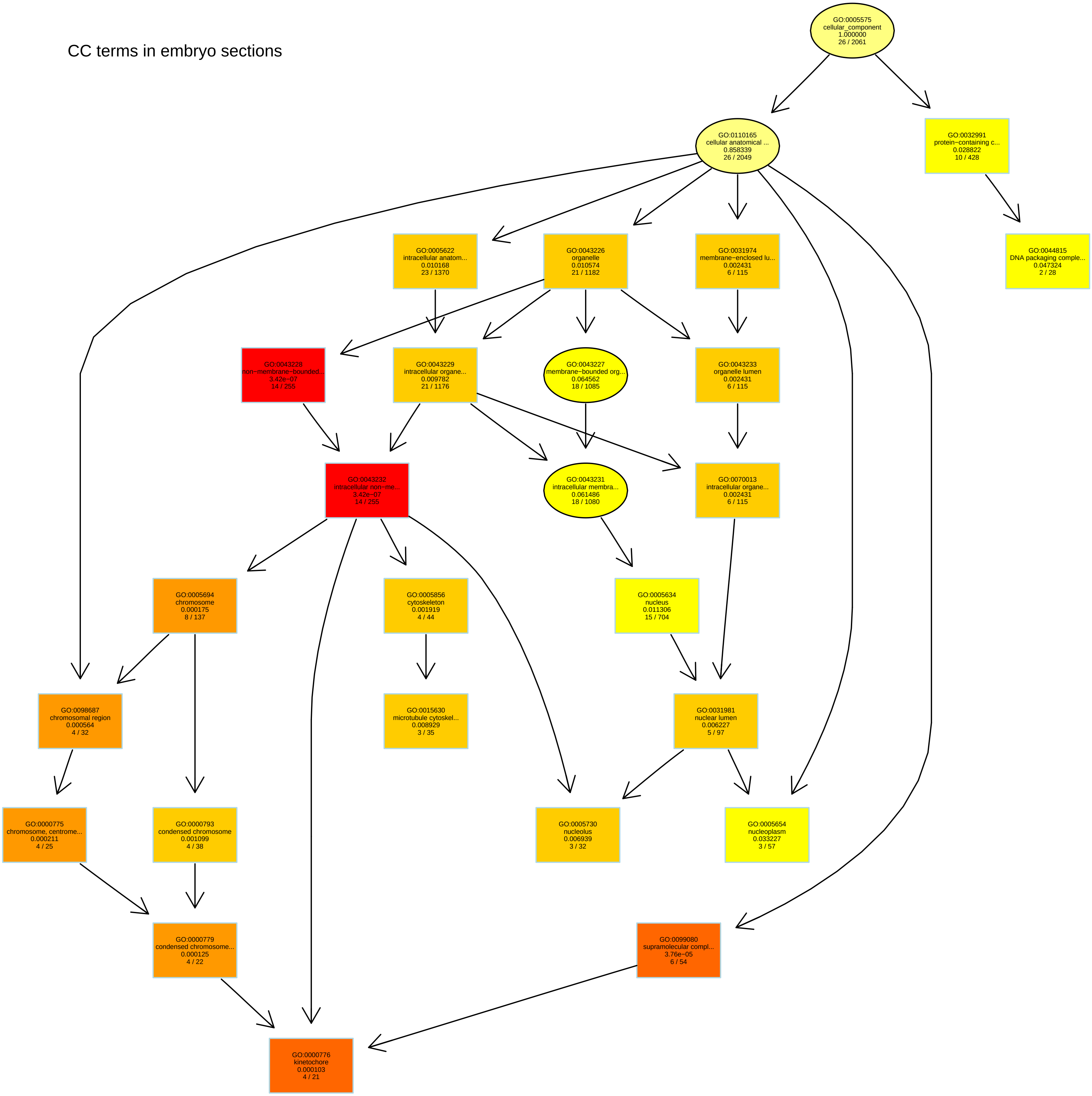
