## Additional file 8 for "Sorghum embryos undergoing B chromosome elimination express B-variants of mitotic-related genes"

Additional file 8: Alignments of A/B chromosome variants of candidate genes. Alignments were obtained using EMBOSS Needle Pairwise Sequence Alignment tool. An asterisk (*) indicates positions with identical residues in both sequences, a colon (:) marks positions with conserved substitutions, and a period (.) signifies positions with semi-conserved substitutions.

### CENH3 – 98.7% identity

```
CENH3_A,g22204      MARTKHQAVRRPTQKPKKKLQLERGGAGTSATPERNAGAGEGTAARGARGRGEKKKMRWR      60
CENH3_B,g22416      MARTKHQAVRRPTQKPKKKLQLERGGAGTSATPERNAGAGEGTAARGARGRGEKKKMRWR      60
                    ************************************************************

CENH3_A,g22204      PGTVALREIRRYQKSTEPLIPFAPFVRVVKEITGFVTYWRIGRYTPEALLALQEAAEFHL     120
CENH3_B,g22416      PGTVALREIRRYQKSTEPLIPFAPFVRVVKEVTGFVTYWRIGRYTPEALLALQEAAEFHL     120
                    *******************************:****************************

CENH3_A,g22204      IELFEVANLCAIHAKRVTVMQKDIQLARRIGGRRWS*   156
CENH3_B,g22416      IELFEVANLCAIHAKRVTVMQKDMQLARRIGGRRWS*   156
                    ***********************:*************
```

### CENP-C – 74.32% identity

```
CENP-C_A,g28463     MDVADPLCAISSPARLLPRTLGPAPASSSSSSSSTGLLEAIAVARSLKGSEELLKQAKMV      60
CENP-C_B,g5138      ------------------------------------------------------------       0

CENP-C_A,g28463     LKEHGDIQALYPDDGVQARPPVNGSKEQQGRRPALNRKRSRFTMKETASKPMPVVDRSKL     120
CENP-C_B,g5138      ---------------------------------------------------MPVVDRSKL       9
                                                                       *********

CENP-C_A,g28463     TNISDPVEFFMTLDRLD--EAEEEILRLTGAAEKRVLNFDPVDEPKRQPGFRG---RKSV     175
CENP-C_B,g5138      TNTSDPFKNFKTVDPLDIAEAEEEILRLSGAAEKRVLNFDPVDEPKRQPGFRGYASRKSV      69
                    ** ***.: * *:* **  *********:************************   ****

CENP-C_A,g28463     CSFRIIEDADTQDPLEVPASQTGSQFPQDVMHVVADKNERVPSSSDEAISGKEDSLAEKD     235
CENP-C_B,g5138      CSFRTNEDADTQDPLEVPASQTGSQFPQDVMHVVADKNERVPSSSGEAISGKE-------     122
                    ****  ***************************************.*******

CENP-C_A,g28463     GRDDLTYLLTSMQHLDESKEEEFIRKTLGVKDIRKERVSLRNSIPGVRPLRTEREVSMRV     295
CENP-C_B,g5138      -FDDLTYLLTSMQHLDESEEEEFIRKTLGVKEIRKERVSL-------RPLRTEREVSMRV     174
                      ****************:************:********       *************

CENP-C_A,g28463     HPPESPLPQPLQDRISELEKHLFHEDAANAKCTDDESEGSPDIVMGEPSLVHDSSDVPMT     355
CENP-C_B,g5138      HPPESSLPQPLQDQISELEKHLFHEDAANAECTDDEYEGSPDIVMGEPSLVHDSSDVPMT     234
                    ***** *******:****************:***** ***********************

CENP-C_A,g28463     DENSTVSEIDRDTPNLGARAADHILDPEPDMPDHAYERQPGDSSVGLCRDTQVAKENEAC     415
CENP-C_B,g5138      DENSTVSEIHRDTPNLGARAADHILDPEPDTPDHAYERQPGGSSVGLCRDTEVTKENEAW     294
                    *********.******************** **********.*********:*:*****

CENP-C_A,g28463     RRSNISV----------EEDDVPIDHPTIGRSTSETEASSHHLERSSTEELVNKPGRHGA     465
CENP-C_B,g5138      RRSNISVEACISIYFALQEDDVPIDHPTIGRSTSETEASSHHLEGSSTEELVNKPGRHGA     354
                    *******          :************************** ***************

CENP-C_A,g28463     PDGINRTLHAAEDIIQHLEVVKEGGVLQDKSSQSLEMPLEDIDPVNQPQMHGGSTKKLAP     525
CENP-C_B,g5138      PDGIDSTLHAAEDIIQHLST--------DKSSQSLEMPLEDIDPVNQPQMHGGSTKKLAP     406
                    ****: ************..        ********************************

CENP-C_A,g28463     DVCNALSLTKQKKQQAAQEGKMKRQSKRGKKVADESSHVLEIPQANLDSENQPHNDDVNI     585
CENP-C_B,g5138      DLCNALSLTKRKKQQAAQAGKMKRQSKRGKKVADEPSHVLEIPKANLDNDDVNIEKSAHD     466
                    *:********:******* **************** *******:****.::   :...:

CENP-C_A,g28463     EQQTVLSITP-----------------------SPNHA----------------------     600
CENP-C_B,g5138      -SCNALSLTKQKQQAAQEGKMKKQPKRGKKVADEPSHVLEIPKANLDNDDVNIEKSAHDS     525
                     . ..**:*                        .*.*.

CENP-C_A,g28463     --------EGQKGAQITNKTKKMNQRKILGDGGLAQPSVVRRSTRTRSRPLEHWLGERLL     652
```

```
CENP-C_B,g5138       CNALSLTKQKQQAAQEGKMKKQPKRGKKVADGGVAQP--LRRSTRTRSRPLKHWLGERLL      583
                             : *:.**  : .*: :: * :.***:***  :***********:********

CENP-C_A,g28463      YGPINDTLPAVIGIKAYSPDQDGKRTLKVKSFVPDQFSDLVAKSAKY*    699
CENP-C_B,g5138       YGPINDTLPAVIGIKTYSPDQDGKRTLKVESFVPDQFSDLIAKSAKY*    630
                     **************:*************:**********:*******
```

### Mis12 – 81.37% identity

```
Mis12_B,g14525      MEDCDESAAT-AAEAALGLNPQHFFNEVHGIIADISAGAFE----AAAAPGVVGAAKAAE      55
Mis12_A,g32810      MEDCDESAAAAAVEAALGLNPQLFVNEVHGIIADIGAGAFEYGLQAAAAPGVVGAAKAAE      60
                    *********: *.********* *.**********.*****    ***************

Mis12_B,g14525      KATDLQRGLNAIHHVVKNRLDKRMTNWKAFCLRHCFDVPEGFVAAEDDRC-AKESHKDET      114
Mis12_A,g32810      KATDLQRGLNAIHHVVKGRLDKRMTNWKAFCLRHCFDVPEGFVAAEDDRSRAKESHKDET      120
                    *****************.*******************************. *********

Mis12_B,g14525      SGLNLELDSLRRKLESATKESQNLEREMSSLERQTTCKRQLDSSLSEIQKLFKDKSVQEN      174
Mis12_A,g32810      SDLDLELDSLRRKLESANKESQNLEREISSLERQTTYKRQLDSSLSEIQKLFEEKSVQEN      180
                    *.*:*************.*********:******** ***************::******

Mis12_B,g14525      FEGVVKAASVLKQKIIDMKKKRTATTCSDQSVWNTNNLTDNRRQTLDNGFTACAEDIQET      234
Mis12_A,g32810      FEGLLKAVPVLKQKIIDLNKKRTATTCSDQQVWNTNNLTDNRRQTLDN-------DIQET      233
                    ***::**. ********::***********.*****************       *****

Mis12_B,g14525      ISILKNKCHVGPLSLPQGAQDQGRRHLDGTSSSNIPGISKKA-RVKGIKEGKNQGI*   289
Mis12_A,g32810      ISIVKNKCRGAVI-TPTGSAGPGPQ---------APGWHQQQHSRDQQEGKN*---   276
                    ***:****: . :  * *: . * :          **  ::  : :. :****
```

### Nuf2 – 89.47% identity

```
Nuf2_B,g8044       MAMQKVQEKTNTLEMYTKFSEKLANHLSKISAVLEKSAAAKASEKDVKAHKEKISDQNLE 60
Nuf2_A,g53293      MAMQKVQEKTNTLEVYTKVSEKLAKHLSKISTVLEKSAAAKASEKDVKAHKEKISDQNLE 60
                   **************:***.*****:******:****************************

Nuf2_B,g8044       IKALRNKAAEWQMRVLENEAKLKAKEKERDQRVGENNRKMTALKSEVELEHKCLEERQRK 120
Nuf2_A,g53293      IKALRNKAAEWQMKVLENEAKLKAKE-ERDQRVGENKRKMAALKSEVESEHKCLEEKQRK 119
                   *************:************ *********:***:******* *******:***

Nuf2_B,g8044       IKEKIDKGSELCSQADSVAEAGWKKIEEIYAKFDQVCEAAKVYMDGMDQSFDETDEAAVT 180
Nuf2_A,g53293      IKEKIDKGSELCSQADSVAEAGRKKIEEIHGKFDQVSEAAKMYVDGMDQSFDETDEDAVM 179
                   ********************** ******:.*****.****:*:************ **

Nuf2_B,g8044       LSTVARGGA*    189
Nuf2_A,g53293      LSTIARNGA*    188
                   ***:**.***
```

### Naa50 – 98.36% identity

```
Naa50_A,g12466      MGAGDGEVAASKEKGSGGAGGGGVERTSLDGVRDKNVMQLKKLNTALFPVRYNEKYYQDA      60
Naa50_B,g13475      MGAGDGEVAASKEKGSGGAGGGGVERTSLDGVRDKNE--LKKLNTALFPVRYNEKYYQDA      58
                    ************************************   *********************

Naa50_A,g12466      IASKDFSKLAYYSDICVGAIACRLEKKEGGAIRVYIMTLGVLAPYRGLGIGTKLLNHVFD      120
Naa50_B,g13475      IASKDFSKLAYYSDICVGAIACRLEKKEGGAIRVYIMTLGVLAPYRGLGIGTKLLNHVFD      118
                    ************************************************************

Naa50_A,g12466      LSAKQNISEIYLHVQTNNDDAIAFYKKFGFEITQTIHNYYMNITPPDCYVLTKFIGQAAT      180
Naa50_B,g13475      LSAKQNISEIYLHVQTNNDDAIAFYKKFGFEITQTIHNYYMNITPPDCYVLTKFIGQAAT      178
                    ************************************************************

Naa50_A,g12466      KK*    182
Naa50_B,g13475      KK*    180
                    ***
```

### SMC3 – 99.01% identity

```
SMC3_A,g38923      MYIKKVIIEGFKSYREEISTEPFSPKVNVVVGANGSGKSNFFHAIRFVLSDMFQNLRSED 60
SMC3_B,g36126      MYIKKVIIEGFKSYREEISTEPFSPKVNVVVGANGSGKSNFFHAIRFVLSDMFQNLRSED 60
                   ************************************************************

SMC3_A,g38923      RGALLHEGAGHSVVSAFVEIVFDNSDNRIPVDKEEVRLRRTVASKKDEYYLDGKHVSKTE 120
SMC3_B,g36126      RGALLHEGAGHSVVSAFVEIVFYNSDNRIPVDKEEVRLRRTVASKKDEYYLDGKHVSKTE 120
                   ********************** *************************************

SMC3_A,g38923      VMNLLESAGFSRSNPYYVVQQGKIASLTLMKDSERLDLLKEIGGTRVYEDRRKESLKIMT 180
SMC3_B,g36126      VMNLLESAGFSRSNPYYVVQQGKIASLTLMKDSERLDLLKEIGGTRVYEDRRKESLKIMT 180
                   ************************************************************

SMC3_A,g38923      ETANKRKQIDQVVHYLEERLRELDEEKDELKKYQQLDKQKRSLEYTILDHELNDARNELA 240
SMC3_B,g36126      ETANKRKQIGQVVHYLEERLRELDEEKDELKKYQQLDKQKRSLEYTILDHELNDARNELA 240
                   *********.**************************************************

SMC3_A,g38923      SMDDNRRKISESMSLADNEVVDVREMIKSFDKEIKVSTKGINDTKAQKEGVEKRRTEALK 300
SMC3_B,g36126      SMDDNRRKISESMSLADNEVVDVREMIKSFDKEIKVSTKGINDTKAQKEGVEKRHTEALK 300
                   ******************************************************:*****

SMC3_A,g38923      VVAQIELDLRDIKDRIVNEKRAKDEAARDLQSVRRESEKSKSELAEISKVHLTKLKEEEE 360
SMC3_B,g36126      VVAQIELDLRDIKDRIVNEKRAKDEAARDLQSVRRESEKSKSELAEISKVHQTKLKEEEE 360
                   *************************************************** ********

SMC3_A,g38923      ISKSIMDREKRLSILYQKQGRATQFANKAARDKWLQKEIEDLKPVLLSNRKQEGLLQEEI 420
SMC3_B,g36126      ISKSIMDREKRLSILYQKQGRATQFANKAARDKWLQKEIEDLKPVLLSNRKQEGLLQEEI 420
                   ************************************************************

SMC3_A,g38923      QKLKDDITELTNYIESRKNESSKLEEALAKRHNDYNDLRKQRDVLQEERKSYWKEESEVT 480
SMC3_B,g36126      QKLKDDITELTNYIESRKNESSKLEEALAKRHNDYNDLRKQRDVLQEERKSYWKEESEVT 480
                   ************************************************************

SMC3_A,g38923      AELDRLQEELVKAQKSLDHATPGDIRRGLTSVNNIIKECSITGVFGPVLELIDCEEKFFT 540
SMC3_B,g36126      AELDRLQEELVKAQKSLDHATPGDIRRGLTSVNNIIKECSITGVFGPVLELIDCEEKFFT 540
                   ************************************************************

SMC3_A,g38923      AVEVTAGNSLFHVVVENDDISTRIIEHLNKRKGGRVTFIPLNRVKASDLSCPQSPDFVPL 600
SMC3_B,g36126      AVEVTAGNSLFHVVVENDDISTRIIEHLNKRKGGRVTFIPLNRVKAPDLSCPQSPDFVPL 600
                   ********************************************** *************

SMC3_A,g38923      LKKLKYRAEHRRAFEQVFGRTVICRDLETATKVARSNGLDCITLDGDQVGKKGAMTGGFY 660
SMC3_B,g36126      LKKLKYRAEHRRAFEQVFGRTVICRDLETATKVARSNGLDCITLDGDQVGKKGAMTGGFY 660
                   ************************************************************

SMC3_A,g38923      DSRRSKLKLVKIFRDNKTAIEKKATHLEAVGNKLKDIDKKITDLVTKQQQMDAERDHAKL 720
SMC3_B,g36126      DSRRSKLRLVKIFRDNKTAIEKKATHLEAVGNKLKDIDKKITDLVTKQQQMDAERDHAKL 720
                   *******:****************************************************

SMC3_A,g38923      ELEQFKVDIARAMKQKASLEKALGKKEKSLDNIRNQIEQVQSSIAMKNDEMGTELIDQLT 780
SMC3_B,g36126      ELEQFKVDIARAMKQKASLEKALGKKEKSLDNIRNQIEQVQSSIAMKNDEMGTELIDQLT 780
                   ************************************************************

SMC3_A,g38923      SEERDLLSRLNPEITDLKERFLMCKNSRIEIETRKEELETNLSTNLIRRQKELEAIISSA 840
SMC3_B,g36126      SEERDLLSRLNPEITDLKERFLMCKNSRIEIETRKEELETNLSTNLIRRQKELEAIISSA 840
                   ************************************************************

SMC3_A,g38923      DSRTLPLEAEAKEQELKSSKRNLDELTSLLKANVDAINNFTRKMDDLKRKRDDLKTREAI 900
SMC3_B,g36126      DSRTLPLEAEAKEQELKSSKRNLDELTSLLKANVDAINNF-RKMDDLKRKRDDLKTREAI 899
                   **************************************** *******************

SMC3_A,g38923      LEQSVQDGAKDLEQLMNSRSTYLAKQEECTKKIRDLGSLPADAFEAYKRKNKKQLHKMLY 960
SMC3_B,g36126      LEQTVQDGAKDLEQLMNSRSTYLAKQEECTKKIRDLGSLPADAFEAYKRKNKKQLHKMLY 959
                   ***:********************************************************

SMC3_A,g38923      DCNEQLKKFSHVNQKALDQYVNFTEQREQLQRRRAELDAGDVKIKELISVLDQRKDESIE 1020
SMC3_B,g36126      DCNEQLKKFSHVNQKALDQYVNFTEQREQLQRRRAELDAGDVKIKELISVLDQRKDESIE 1019
                   ************************************************************
```

```
SMC3_A,g38923      RTFKGVARHFREVFSELVQGGHGYLVMMKKKDGDAVDDD--DEDEDGPRDPGPEGRIEKY 1078
SMC3_B,g36126      RTFKGVARHFRKVFSELVQGGHGYLVMMKKKDGNAVDDDDDDEDEDGPRDPGPEGRIEKY 1079
                   ***********:**********************:*****  ******************

SMC3_A,g38923      IGVKVKVSFTGKGETQSMKQLSGGQKTVVALTLIFAIQRCDPAPFYLFDEIDAALDPQYR 1138
SMC3_B,g36126      IGVKVKVSFTGKGETQSMKQLSGGQKTVVALTLIFAIQRCDPAPFYLFDEIDAALDPQYR 1139
                   ************************************************************

SMC3_A,g38923      TAVGNMIRRLADMADTQFIATTFRPEIAKVADKIYGVTHKNRVSYINVVSKEQALDFIEH 1198
SMC3_B,g36126      TAVGNMIRRLADMADTQFIATTFRPEIAKVADKIYGVTHKNRVSYINVVSKEQALDFIEH 1199
                   ************************************************************

SMC3_A,g38923      DQTHNAS*      1205
SMC3_B,g36126      DQTHNAS*      1206

                   ********
```

### ESD4 – 85.44 % identity

```
ESD4_B,g25900      ------------------------------------------------------------ 0
ESD4_A,g3140       MLVAMSLHARRPEAALRRCRWPCPSTLSLHRWVGRPRAHTGGRARPWSLHPTFAGGHGSS 60


ESD4_B,g25900      ----------MDNEAIQQQQSNVGSPAGAGRGGHMLADLCPWEVTVQHSDERDGVEDAEA 50
ESD4_A,g3140       PSRHALQQLTMDNEATQQQQSNVGSPAGAGRGGHVLADFCPPEVTVQHGDERDGIEDAEA 120
                             ***** ******************:***:** ******.*****:*****

ESD4_B,g25900      TPPLALGPDEDLPSSCRKNLSKAKRRKKSHNSNRTTNILSQRTSSFLHEKYDGDQEEILN 110
ESD4_A,g3140       TLSLALGPDEDLPSSGRKNLSKVKRRKKSHNSNRTTNILSQ---SFLQQKCDGDQEEMLN 177
                   *  ************ ******.******************   ***::* ******:**

ESD4_B,g25900      NTKKDNGLPLCESEDTSLDSRSKISMSIVAIPEDYVSNQECDGDQEEMLNNAKKDNTLPL 170
ESD4_A,g3140       NTKKDNGLSFCESEETSLDSRSAISVSIVAIPQDYVCNQECDGDQEELLNNAKKDNALPL 237
                   ******** :****:******* **:******:***.**********:********:***

ESD4_B,g25900      CEYEET-SLDNKSEKSVSIVAIPEDYVCNQECDGDREEMLNDIEKDNGLSLRESEETSLD 229
ESD4_A,g3140       CESEETSSLDNKSKKPVSIVAIPEDYVCNQECDGDQQEMLNDTEKDNGLPLCESEETSLD 297
                   ** *** ******:* *******************::***** ****** * ********

ESD4_B,g25900      SKYEKSVSIVAIPEDYVCNQADLDIIEPIKKFPYKPGKEQVVLIDDAFIDRMNMECLFQP 289
ESD4_A,g3140       SKSEKLVSILAIPEDYVCNQADLDIIEPIKKSLINRERNKLCS--SMMLS*--------- 345
                   ** ** ***:*********************   :  ::::    . ::.

ESD4_B,g25900      NAFLNDQVINAYITLLRAQDHLKLRACGKVFLENSLISSILRRDGDDKIKMEDLYPTGDK 349
ESD4_A,g3140       ------------------------------------------------------------ 345


ESD4_B,g25900      NGISTVKKRVLSYLDHDMVFIPINIENTHWYLGVVNAKEREIQVLDSMGTIFGRQDLILT 409
ESD4_A,g3140       ------------------------------------------------------------ 345


ESD4_B,g25900      IKGLQKQIDIVSQHKNLNGHKWPDIQVSSWPVREIHFEQKMQTDGCSCGLFLLKYVEHWT 469
ESD4_A,g3140       ------------------------------------------------------------ 345


ESD4_B,g25900      GEGLSKNITQEDMTQFRTKLAAILLSSDLNKRKGSLLVKVDDEAIGSQSEVEILQSSNSP 529
ESD4_A,g3140       ------------------------------------------------------------ 345


ESD4_B,g25900      CKRKNYHQTPASCSPDRTMDPFLSCGLSTFDMPVTKEDMIDLLCDYLMTIDDAETLETSW 589
ESD4_A,g3140       ------------------------------------------------------------ 345


ESD4_B,g25900      VRSFQPYNITLTVRQLQASLRMNQPMPTDCFNMGVRLLAYRENKRLTSANLMISKHYMDL 649
ESD4_A,g3140       ------------------------------------------------------------ 345
```

```
ESD4_B,g25900      RFSTTYEAAQKPRSKKLNVQELAKSLETWPYMKYDASLCRFLLMPWKRGGLNLFVFDREE 709
ESD4_A,g3140       ------------------------------------------------------------ 345


ESD4_B,g25900      KTLTVLDPTPIPDWCKDMPYKNYVRRIINVSNGYLLAMGVQAPERAVDVFAWKHILPSGI 769
ESD4_A,g3140       ------------------------------------------------------------ 345


ESD4_B,g25900      PVIEDRNLNAFILLQFMSAWNNGKLMPISMDLKRLRKKFVIDLLAYDGNSRRCMIPLSIR 829
ESD4_A,g3140       ------------------------------------------------------------ 345


ESD4_B,g25900      EYLSRITGIRQ   840
ESD4_A,g3140       -----------   345
```

ESD4 – 40.82 % identity

```
ESD4_B,g37251      -----MVETRRQKRVNPQDNEDEQVPNFSMPASPAAAEHHIAVEVCMEAPPSRRTRARCA 55
ESD4_A,g56148      MNIAAIIGHTKFEEFRPTSSANEHIHALGVSYEPRSLEQSI------------------- 41
                        ::   : :...* .. :*::  :.:  .* : *: *

ESD4_B,g37251      ASQEAPPSHRTRARCAASRATTTASDSDAPPSSSSTADFWK-QANDALKPRPMKGAVWTD 114
ESD4_A,g56148      -------------------------------NKQQMEQIFQRMNNNHLVFQNDMGRQFMN 70
                                                  .... ::::   *: *  :   *  : :

ESD4_B,g37251      EE------NILFCNCCVAMIEAGEMGKREPSLKGDMLEKFCERFNSSSEERQRREPKQFL 168
ESD4_A,g56148      LEHKLDAKLLAYGEKIAEIRMGGDVGFRISKLEDDFAE--------------------LK 110
                    *       : : :  . :  .*::* *  .*:.*: *                    :

ESD4_B,g37251      KKFEKLKELYDK---------YH-EHSALGNER-------------RL-----RMKTKDA 200
ESD4_A,g56148      REFQELRQLLLAHLQSMTTAPYIQERSTTQPEKTTEFFKPTEASTRQDKIMAENMVPQDA 170
                   ::*::*::*            *  *:*:   *:             :      .*  :**

ESD4_B,g37251      TKDLKDLVHFPPKYYDLLHKIFEKPENLNPINNIQDRDDWFEKYKNQSTKYLDYFTLTPE 260
ESD4_A,g56148      P--TTQVTPQKPAFQDKIMAENTVPED-----------APTHQIAPLKPAFDNDYLITTE 217
                       .::.   * : * :      **:              .:    .  : : : :* *

ESD4_B,g37251      DCEAIKFIQDSYQYAEVVDIEGNLLRVLQLRPFVYGCCVKDDLINAYAHIAASEEKNNNK 320
ESD4_A,g56148      DAEAFYFITHSYAEAEVVQINDLVLRIEQLRTHLTGGFIHDQAINAYAHISSVET--DST 275
                   *.**: ** .**  ****:*:. :**: *** .: *  ::*: *******:: *   :..

ESD4_B,g37251      GFITTFEAQKLSQDNGELDD--KRRSWVINVGKKCLEKELIFIPVFNKQKYDWSLLVLNK 378
ESD4_A,g56148      SFIPTFQVQKLLGETGGINNPKQTKKWAELIAKKCIGKNLVFVPMN-VNTNHWVLLVLNF 334
                   .** **:.***  :.* :::  : :.*.  :.***: *:*:*:*:   :. .* *****

ESD4_B,g37251      KEEGGEFQILSP---LPGLRNETVEKTLVKSLQKCIDEAVKDGQST----VDLSQWEIKD 431
ESD4_A,g56148      I--KGEVQILNSLASNPNNRDVVKEHTVVGNIQECIDSSIADGSVSVPQPINIMQWETEV 392
                       **.***.     *. *: . *:*:* .:*:***.:: **. :    ::: *** :

ESD4_B,g37251      YSTHIPQQSDMTSSGVYMIKYMLGWDGSKMDQNFTQDDMNVFRLKICCSLLRSKHNKIQR 491
ESD4_A,g56148      YS-NIPQQTDGHSCGAFMLKYMLTWTGDKMSEHFTQAHINIFKRKISSALLRSDCNKLRL 451
                   ** :****:*  *.*.:*:**** * *.**.::*** .:*:*: **..:****. **::

ESD4_B,g37251      ASYDVPIMKKAYLATSQKDDCDD--NDDDDDLQMADNLTDLMTNN*-------------- 534
ESD4_A,g56148      GSYKDLITKAAYDAKRAEIQREEMAQAADGDIQVINNALDASNSNTKTIKRRGRPKKNEA 511
                   .**.  * * ** *.  : : ::  :  *.*:*: :*  *  ..*

ESD4_B,g37251      ------------------------------------------------------------ 534
ESD4_A,g56148      AENDGKDISNPIDASNSNKRKRGRPKKNDKTPQSVKDLLPTTPIANRVERPNRRVSNPGP 571


ESD4_B,g37251      ----------    534
ESD4_A,g56148      LQLSPYSKF*    580
```

ULP1B – 85.44% identity

```
ULP1B_B,g40710      ------------------------------------------------------------      0
ULP1B_A,g3140       MLVAMSLHARRPEAALRRCRWPCPSTLSLHRWVGRPRAHTGGRARPWSLHPTFAGGHGSS      60


ULP1B_B,g40710      ----------MDNEAIQQQQSNVGSPAGAGRGGHMLADLCPWEVTVQHSDERDGVEDAEA      50
ULP1B_A,g3140       PSRHALQQLTMDNEATQQQQSNVGSPAGAGRGGHVLADFCPPEVTVQHGDERDGIEDAEA      120
                              ***** ******************:***:** ******.*****:*****

ULP1B_B,g40710      TPPLALGPDEDLPSSCRKNLSKAKRRKKSHNSNRTTNILSQRTSSFLHEKYDGDQEEILN      110
ULP1B_A,g3140       TLSLALGPDEDLPSSGRKNLSKVKRRKKSHNSNRTTNILSQ---SFLQQKCDGDQEEMLN      177
                    *  ************ ******.******************   ***::* ******:**

ULP1B_B,g40710      NTKKDNGLPLCESEDTSLDSRSKISMSIVAIPEDYVSNQECDGDQEEMLNNAKKDNTLPL      170
ULP1B_A,g3140       NTKKDNGLSFCESEETSLDSRSAISVSIVAIPQDYVCNQECDGDQEELLNNAKKDNALPL      237
                    ******** :****:******* **:******:***.**********:********:***

ULP1B_B,g40710      CEYEET-SLDNKSEKSVSIVAIPEDYVCNQECDGDREEMLNDIEKDNGLSLRESEETSLD      229
ULP1B_A,g3140       CESEETSSLDNKSKKPVSIVAIPEDYVCNQECDGDQQEMLNDTEKDNGLPLCESEETSLD      297
                    ** *** ******:* *******************::***** ****** * ********

ULP1B_B,g40710      SKYEKSVSIVAIPEDYVCNQADLDIIEPIKKFPYKPGKEQVVLIDDAFIDRMNMECLFQP      289
ULP1B_A,g3140       SKSEKLVSILAIPEDYVCNQADLDIIEPIKKSLINRERNKLC--SSMMLS*---------      345
                    ** ** ***:*********************   :  ::::   .. ::.

ULP1B_B,g40710      NAFLNDQVINAYITLLRAQDHLKLRACGKVFLENSLISSILRRDGDDKIKMEDLYPTGDK      349
ULP1B_A,g3140       ------------------------------------------------------------      345


ULP1B_B,g40710      NGISTVKKRVLSYLDHDMVFIPINIENTHWYLGVVNAKEREIQVLDSMGTIFGRQDLILT      409
ULP1B_A,g3140       ------------------------------------------------------------      345


ULP1B_B,g40710      IKGLQKQIDIVSQHKNLNGHKWPDIQVSSWPVREIHFEQKMQTDGCSCGLFLLKYVEHWT      469
ULP1B_A,g3140       ------------------------------------------------------------      345


ULP1B_B,g40710      GEGLSKNITQEDMTQFRTKLAAILLSSDLNKRKGSLLVKVDDEAIGSQSEVEILQSSNSP      529
ULP1B_A,g3140       ------------------------------------------------------------      345


ULP1B_B,g40710      CKRKNYHQTPASCSPDRTMDPFLSCGLSTFDMPVTKEDMIDLLCDYLMTIDDAETLETSW      589
ULP1B_A,g3140       ------------------------------------------------------------      345


ULP1B_B,g40710      VRSFQPYNITLTVRQLQASLRMNQPMPTDCFNMGVRLLAYRENKRLTSANLMISKHYMDL      649
ULP1B_A,g3140       ------------------------------------------------------------      345


ULP1B_B,g40710      RFSTTYEAAQKPRSKKLNVQELAKSLETWPYMKYDASLCRFLLMPWKRGGLNLFVFDREE      709
ULP1B_A,g3140       ------------------------------------------------------------      345


ULP1B_B,g40710      KTLTVLDPTPIPDWCKDMPYKNYVRRIINVSNGYLLAMGVQAPERAVDVFAWKHILPSGI      769
ULP1B_A,g3140       ------------------------------------------------------------      345


ULP1B_B,g40710      PVIEDRNLNAFILLQFMSAWNNGKLMPISMDLKRLRKKFVIDLLAYDGNSRRCMIPLSIR      829
ULP1B_A,g3140       ------------------------------------------------------------      345


ULP1B_B,g40710      EYLSRITGIRQ*   840
ULP1B_A,g3140       ------------   345
```

NCAPH – 94.96 % identity

```
css2_B,g47463       MPPAEDAPPQTPPPARGTAAALRVLLQSPPPAFPLGSNDDQQERARARAAARAASVRRRS      60
```

```
css2_A,g47542      MPPAEDAPPLTPPPARGTAAASRVLLQSPPPAFPLGSNDDQQERARARAAARAASVRRRS      60
                   ********* *********** **************************************

css2_B,g47463      LAASIAPSKDPRHDLLNREQVMDLFHNCIKLASENKINQKNTWELGLIDHLSEIIQAGAD      120
css2_A,g47542      LAASIAPSKDPRHDLLNREQVMDLFHNCIKLASENKINQKNTWELGLIDHLSEIIQAGAD      120
                   ************************************************************

css2_B,g47463      EDEETNFQKASCTLEAGVKIYSLRVDSVHSEAYKVLGGINRAGRGEEADLEEGSNVEPAQ      180
css2_A,g47542      EDEETNFQKASCTLEAGVKIYSLRVDSVHSEAYKVLGGINRAGRGEEADLEEGSNVEPAQ      180
                   ************************************************************

css2_B,g47463      DEGINKKNADRRISPASTLESSFEALNVKKFDVAFTVDPLYHQTTAQFDEGGAKGLLLYN      240
css2_A,g47542      DEGINKKDADRRISPASTLESSFEALNVKKFDVAFTVDPLYHQTTAQFDEGGAKGLLLYN      240
                   *******:****************************************************

css2_B,g47463      LGVYGSCHVLFDSFEAPDNCILSDMQTEQAEAIDLSFAKEQIEEMVTQIHLCDDISPTLR      300
css2_A,g47542      LGVYGSCCVLFDSFEAPDNCILSDMQTEQAEAIDLSFAKEQIEEMVTQMRLCNDISPTLR      300
                   ******* ****************************************::**:*******

css2_B,g47463      DIVAQFDEENQRPSHRLSPGQMPVMEDPMDEDNEADDDDSMLPDSGTWDFGGCHDHEDAY      360
css2_A,g47542      DIVAQFDEENQRPSHRLSPGQMPVMEDPMDEDNEADDNDSMQPDSGTWDFGGCHDHEDAY      360
                   *************************************:*** ******************

css2_B,g47463      NENCNPMDSISTNYQEEFNEYTVEIPQGTIVDERLEKIADLLLLGMGSSKANAWAGPEHW      420
css2_A,g47542      NENCNPMDSISTNYQEEFNEYTVEIPQGTIVDERLEKIADLLLLGMGSSKANAWAGPEHW      420
                   ************************************************************

css2_B,g47463      KYRKAKDLEAVPTSSGDSEITNKTQKRSKDKPDIDFTKALDNEHPNIFAAPKNPKLLVLP      480
css2_A,g47542      KYRKAKDLEAVPTSSGDSEITNKMKKRSKDKPDIDFTKALDNEHPNIFAAPKNAKSLLLP      480
                   *********************** :**************************** * *:**

css2_B,g47463      ANRAMCSNKLPEDCHYQPESLVKLFLLPDVLCLAKRRRKSLDAPVDNGDEFIPSEPWEDD      540
css2_A,g47542      ANRAICSNKLPEDCHYRPESLVKLFLLPDVLCLAKRRRQSLDAPLDNGDEFIPSEPWEDG      540
                   ****:***********:*********************:*****:**************.

css2_B,g47463      SFCTDHVDEGHVCSDLEEPVNLINKPRQVNKIDIQYDKVSKQVDVHALKEVLWNHIHASA      600
css2_A,g47542      SFCTDHVDEGHMCSDVEEPINLINKPRQVNKIDIQYDKVSKQVDVHALKEVLWNHIHASA      600
                   ***********:***:***:****************************************

css2_B,g47463      ETDGQEREETGSPLCLSRVLH---------VTSDISPHLYFICLLHLANEHSLKLCDRPT      651
css2_A,g47542      ETDGQEREETGSPLCLSQVLHDLPSSNPDAVTPDISPHLYFICLLHLANEHSLKLCDRPT      660
                   *****************:***         ** ***************************

css2_B,g47463      LDEIDIYMPTSPLVK*       666
css2_A,g47542      LDEIDIYMPTSPPVK*       675
                   ************ ***
```

### NCAPG – 98.2% identity

```
css3_B,g10797     MAPAVAVTGAGDSDRLAREVARVLDECYASHAVHPRKLRELAALRSSSSGG-GGGGGPFL  59
css3_A,g5388      MAPAAAVTGAGDSDRLAREVARVLDECYASHAVHPRKLRELAALRSSSSGGGGGGGGPFL  60
                  ****.********************************************** ********

css3_B,g10797     AAFCVAVTPLFALARRSAGSDRIARFVAAFASASAS-SADGGGSGNGFLEEFLRFLVTAS  118
css3_A,g5388      AAFCVAVTPLFALARRSAGSDRIARFVAAFASASASSSADGGGSGNGFLEEFLRFLVTAS  120
                  ************************************ ***********************

css3_B,g10797     KAAHRPARFRACQIISEIIMRLPDDAEVSDQIWDEAIDAMKVRVQDKIAAIRTFAVRALS  178
css3_A,g5388      KAAHRPARFRACQIISEIIMQLPDDAEVSDEIWDEVIDAMKVRVQDKIAAIRTFAVRALS  180
                  ********************:*********:****.************************

css3_B,g10797     RFAIDGEDGGIVDLFLGTLDIEQNAEVRKAIVFSLPPSNNTLESVVESTLDISESVRRAA  238
css3_A,g5388      RFAIDGEDGGIVDLFLRTLDIEQNAEVRKAIVFSLPPSNNTLESVVESTLDISESVRRAA  240
                  **************** *******************************************

css3_B,g10797     YSVLSTKFPLQSLTIKQRTTVLHRGLSDRSVSVNNVCLKMLKDEWLVKNCEGDVISLLRF  298
css3_A,g5388      YSVLSTKFPLQSLTIKQRTTVLHRGLSDRSVSVNNVCLKMLKDEWLVKNCGGDVISLLRF  300
```

```
                  ************************************************** *********

css3_B,g10797     LDVETYESVGESVMAVLLKDGALRVHDGHSIRQYITANGEKEQDSNIQLMDAEVALYWRI  358
css3_A,g5388      LDVETYESVGESVMAVLLKDGALRVHDGHSIRQYITANGEKEQDSNIQLMDAEVALYWRI  360
                  ************************************************************

css3_B,g10797     MCKHLQAEAQAKGSEAAATTGAEAAVYASEATDKNDLLDNVLPSTITDYVDLVKAHLSAG  418
css3_A,g5388      MCKHLQAEAQAKGSEAAATTGAEAAVYASEATDKNDLLDNVLPSTITDYVDLVKAHLSAG  420
                  ************************************************************

css3_B,g10797     PNYHFTSRQLLLLGEMLDFSDTMNRKIASSFLHELLIRPLEHEVDDDGNQIAIGDGVSLG  478
css3_A,g5388      PNYHFTSRQLLLLGEMLDFSDTMNRKIASSFLHELLIRPLEHEVDDDGNQIAIGDGVSLG  480
                  ************************************************************

css3_B,g10797     GDKDWAKAVAELAKKVHSSVGEFEMVVSSVVEELARPCRERTADFMQWIHCLAVTGLLLQ  538
css3_A,g5388      GDKDWAKAVAELAKKVHSSVGEFEMVVSSVVEELARPCRERTADFMQWMHCLAVTGLLLQ  540
                  ************************************************:***********

css3_B,g10797     NTSTLRNLQATAIEPSELLHSLLLPAAKQNHVDVQRAALRCLCLLGLLENRPNAELVKQL  598
css3_A,g5388      NTSTLRNLQATAIEPSELLHSLLLPAAKQNHVDVQRAALRCLCLLGLLENRPNAELVKQL  600
                  ************************************************************

css3_B,g10797     RLSFINGPDLVSAIACKALIDLVTWHGPQEIDRAIGIDLPDPSYKKSQFTQVDLSDMNDD  658
css3_A,g5388      RLSFINGPDLVSAIACKALIDLVTWHGPQEIDRAIGIDSPDPSYKKSQFTQVDLSDMNDD  660
                  ************************************** *********************

css3_B,g10797     DLNIGVLDILFSGFYKGDWEFDLEGDNHDKIPTILGEGFAKILLLSGNFASIPIDLHTVI  718
css3_A,g5388      DLNIGVLDILFSGFYKGGWEFDLEGDNHDNIPTILGEGFAKILLLSGNFASIPIDLHTVI  720
                  *****************.***********:******************************

css3_B,g10797     VAQLIRLYFSEETKELERLKQCLSVFFQHYPALSDKHKSCISNAFVPVMKAMWPGLYGNA  778
css3_A,g5388      VAQLIRLYFSEETKELERLKQCLSVFFQHYPALSDKHKSCISNAFVPVMKAMWPGLYGHA  780
                  **********************************************************:*

css3_B,g10797     GGSPVVISKRRKLAVQASRFMVQMVQTQLLSTESMGQASKSPESAPVSANVSNNFDISEE  838
css3_A,g5388      GGSPVVISKRRKLAVQASRFMVQMVQTQLLSTESMGQASKSPESAPVSANVSNNFDISEE  840
                  ************************************************************

css3_B,g10797     GLAIRIALEVAGCPDKKTAAGKAYALALCKVAVLLRFRQSEQKAIKCMRGLVNHLAASVA  898
css3_A,g5388      GLAIRIALEVAGCPDKKTAAGKAYALALCKVAVLLRFRQSEQKAIKCMRGLVNHLAASVA  900
                  ************************************************************

css3_B,g10797     SDKELVKELAQMAARLKALDACPDEELSQDDADVIFNKLGLDDGFKLNSNQAVPPTPAPR  958
css3_A,g5388      SDKELVKELAQMAARLKALDACPDKELSQDDADVIFKKLGLDDGFKLDSNQAVPPTPAPR  960
                  ************************:***********:**********:***********

css3_B,g10797     SARPPAPARRRARQAPPPSSSDESDEGGDVSVPESVSRVPATPSMTAAAHSQRASKTTAL  1018
css3_A,g5388      SARPPAPARRRARQAPPPSSSDESDEGGDVSVPESVSRIPATPSMTAAARSQRASKTAAL  1020
                  **************************************:**********:*******:**

css3_B,g10797     SKMSAKPPAIASDGSESDDQSDVTSEEDSSAEESS*      1053
css3_A,g5388      SKMSAKPPAIASDGSESDDQSDVTSEEDSSAEESS*      1055
                  ************************************
```

HGV2 – 93.29 % identity

```
HGV2_B,g43483      MASSSENTGAPPEVEQQPQAPPTPNPEPTEAAAAEEEEEEEEPRTLERAQELFDRGAKAIED 60
HGV2_A,g31157      MASSSENTGAPPEVEQQPQAPPTPNPEPTEAAAAEE--EEEPRTLERAQELFDRGAKAIED 58
                   ***********************************  ***********************

HGV2_B,g43483      EDFVEAVDCLSQALEIRTSHYGELAPECASTYFKYGCALLYKAQEESDFLGNVPKSVPNE 120
HGV2_A,g31157      EDFVEAVDCLSQALEIRTSHYGELAPECASTYFKYGCALLYKAQEESDFLGNVPKSVPNE 118
                   ************************************************************

HGV2_B,g43483      ESVKSTASKDDSGTSKVSCTNVEDAMSSKKADAEEGQNSNGKDQETGNGEVEKDDEDDDN 180
HGV2_A,g31157      ESVKSTTSKDDSGTSKVSCTYVAYLLQFKDFLTIETSISLNVSL*--------------- 162
                   ******:************* *   :. *.  : * . * . .
```

```
HGV2_B,g43483      DEKMGDEEDNDLDLSWKMLDIARAIVEKTPDNSMEKVKIYSALAEVATEREDIDNSLSDY 240
HGV2_A,g31157      ------------------------------------------------------------ 162


HGV2_B,g43483      MKALSMLEHLVEPDHRRVVELNFRICLVYELVSKIGDAIPYCAKAISLCKSRIQSLKSSK 300
HGV2_A,g31157      ------------------------------------------------------------ 162


HGV2_B,g43483      DALLAGKDGDASAAEAEGGSEKSDAEKELEQLTSILPDLEKKLEDLEQANPSPAMDEMLK 360
HGV2_A,g31157      ------------------------------------------------------------ 162


HGV2_B,g43483      TIASRVTDAMPRAASFTSSQMATSSNGFDSSVLSTAATTGSTGSTVTDLGVVGRGVKRAS 420
HGV2_A,g31157      ------------------------------------------------------------ 162


HGV2_B,g43483      IKPISAEPAAKKPALDSPSVQGDSSINSEVVPTTQNGDESVSK*    463
HGV2_A,g31157      --------------------------------------------    162
```

### ANAPC15 – 98.97% identity

```
ANAPC15_B,g3841      MLQFPALMRQWPSPPLLPASTLLPVPATSQEDELLLAMAESDLDDKLNEIRKTNSHLVII      60
ANAPC15_A,g6253      MLQFPALMRQWPSPPLIPASTLLPVPATSQEDELLLAMAESDLDDKLNEIRKTNSHLVII      60
                     ***************:********************************************

ANAPC15_B,g3841     *      96
ANAPC15_A,g6253      GKPTGDTKEEYDAEVEDDDADNVEESDGDDFDQETG*  96
                     *************************************
```

### CYCB1_5 – 66% identity

```
g5077.t1        MATRNHHA----ASAAQPANRGAARIAGKQNGA-ATSRPDAARRVLGDVGNVVSDVLNGK     55
g13110.t1       MATRNHRAAVPAAAAPQPVNRGAARIAGKQKDAAAAGRPNATRAALGDIGNVAPSDVLDG     60
                ******:*    *:* **.***********:.* *:.**:*:* .***:***. . : .

g5077.t1        NTLPEGIHGPITMSFGAALVNDVLANNNTIAPAQPVAARTITNPATIVPAKNTNTPHGEK     115
g13110.t1       IKLPEGIHRPITRSFGAQLLKQQALAKNAGAPAQPVAARAVTKPVKKVPAKNIPRPEQE-     119
                 .****** *** **** *:::    :*: *********::*:*.. *****   *. *

g5077.t1        APKVNRKPSDGVAGSSSS------CSVQRNMRTKLVRTPSTILSDLFEAACGLTEKPKEL     169
g13110.t1       ----NRKPSEGAAKDSKGNMNTSEGVAAVQRRKKLVCTLSTVLSARSKAACGLTEKPKPL     175
                    *****:*.* .*..         .  : *.*** * **:**   :********** *

g5077.t1        IEDIDKFDGDDQFAVVDYVEDIYKFYMTAEHESRPNDYMGNQPEITSKTRASLVDRLIHS     229
g13110.t1       VEDIDKFDGDNQLALVDYVEEIYTFYKTAQHEIRPIDYMGNQPEINLNMRATLTDWLIDS     235
                :*********:*:*:*****:**.** **:** ** *********. : **:*.* **.*

g5077.t1        HQRFHLKPETLYLTIYIVDQYLSLQPVPSMELGLVGAAAMLIAWKYEEAWAEKEKIFIEI     289
g13110.t1       HLRFHLMPETLYLTIYIVDRYLSLQPVPRREFQLVGMAAMLIACKYEEIWAPEVNDFIEI     295
                * **** ***********:********  *: *** ****** **** ** : : ****

g5077.t1        --RPFDRHQILHMEKAILNSMNWELAVPTPYHFLLRFAKAASSDDEQLQHMVHFFGELAL     347
g13110.t1       AARAFSRTQILVTEKAILNSIEWNLTVPTPYHFLLRFAKAAGSADEQLQHMIYFFGELAL     355
                  * *.* ***  *******::*:*:**************.* *******::*******

g5077.t1        MDYGMMMTYASRVAACAVYAARLTLKKIPFWTQTLKHHTGLEEQQLIEGARILVSSHAAA     407
g13110.t1       MEYGMVTTYPSTIAACAVYAARLTLRKSPLWTETLRHHTGLHEPQLMEGARMLLRSHA-A     414
                *:***: ** * :***********:* *:**:**:*****.* **:****:*: *** *

g5077.t1        PDAKLKAVYQKYAVEQFGKVALHPPAALSDLV*      439
g13110.t1       PDAKRKTVHEKYATEQFGRVALHPPAALPDLV*      446
                **** *:*::***.****:********* ****
```

### putative helicase – 53,07% identity

```
g5838.t1        MQMNVTPLSNIRPGILDYTICVRVSRLWEFRGDSEENDVKHLDLVLIDQKANAIYAEIPP     60
g31445.t1       -------------------------------------------MVVIDQKGYAMYCEVPP     17
                                                           :*:****. *:*.*:**

g5838.t1        EAIPMTKPHLEEGNIVYMMKITVQRSKPEYRVVENPYMIKLNRRTIIIEAKDQPPEFPKY     120
g31445.t1       EVLSHKKQYLEEGKVLYIWKACVERAKPGYRVVDAPYMLKLIMRTEIFEGNSNDTSFPKY     77
                *.:  .* :****:::*: *  *:*:** ****: ***:**  ** *:*.:.:  .****

g5838.t1        TFSLIPFDKLHEFTRKTQKFLDVIGKITAVSNAARVYSSSGDLRIRRVIQLRDLSGNTID     180
g31445.t1       VFSLTPIEILPQYARRTDRFLDVIGKITAISNAAVARNTSGDLMMRRMITLQDHKGNTVD     137
                .*** *:: * :::*:*::**********:**** . .:**** :**:* *:* .***:*

g5838.t1        LSLSGKRALEFDGETVLKVGQQNHVIAIFVGTLMKLYQMKHHFLSGTSACRWYINQNDIP     240
g31445.t1       LSLSGQRALEFDAEAVFDIGQNHHVIAIFVGTLMKIYREDYKFLSGTSACRWYINENDIP     197
                *****:******.*:*:.:**::************:*: .::*************:****

g5838.t1        EIKAFQKSLPAQAEPIQTLYLQSEDDTKQNFEEKTLLQLKEVDPFTEKDKRFQCTVTIAA     300
g31445.t1       AMRTFQRGLPSQVTPVQQLQLQSEDYMEQGVEEKTLFDLKQIDPLNDKNKRFQCTVTLIS     257
                 :::**:.**:*. *:* * *****  :*..*****::**::**:.:*:********: :

g5838.t1        VPERQPWCYRACRSCNCTIRVQRSGFKCSNDNCFTTQFDWKYKIPFIGSDGTYNLEFILF     360
g31445.t1       LAEKEQWCYRACKVCNSTLILGNDGYDCTKDGCSCKQYDWKYKVCFVVADDTYNLQFMFF     317
                : *:: ******: **.*: : ..*:.*::*.*  .*:*****: *: :*.****:*::*

g5838.t1        EKRGAELIGKTAEALRKQYDMLDTPPEILAWIGHKFTFVVKILSTKSINTIDPSFEVVLI     420
g31445.t1       EKKGVELIGKSAETLRKKYDPSSIPPEIAQWIGHKFTFIVKVLFKRSIKSIEPSFEVVMI     377
                **:*.*****:**:***:**  . ****  ********:**:* .:**::*:******:*

g5838.t1        KEKFGKQPTIPTLSPDHGLSMRASSTKFQADEELPLLVPIGSKYIQEQDSSSQLQVHANM     480
g31445.t1       KERHGKQVTLPNITHKD-----------VSNDDLPPLVTISSKKKIKQASLSSTPQFTDI     426
                **:.*** *:*.:: ..            ::::** ** *.**   :* * *.   .:::

g5838.t1        HTMDIDSS--SYDSNKRALDLTDDENQDDSLLQPKNKRP-R*-- 518
g31445.t1       QHMDIDQQSDAWDTTEKHIQSQIDDEDDERHEEQKRKKLKINE* 469
                : ****..  ::*:.:: ::   *:::*:   : *.*:
```

### TPR repeat-containing protein ZIP4-like – 97% identity

```
g3867.t1        MKISELSPEYRISQLSPECRSPPAHAALLTDLNRVVTDVEALDASDSSSLEKLAADLRCV     60
g10502.t1       MKISELSPEYRISQLSPECRSPPAHAALLTDLNRVVTDVEALDASDSSSLEKLAADLRCV     60
                ************************************************************

g3867.t1        LTNLASAVSSSSSGLNGAFRLKVWNLAFRLWNACVDRANHKFPARGPEAAVAETEIRQAA     120
g10502.t1       LTNLASAASSSSSGLNGAFRLKVWNLAFRLWNACVDRANHKFPARGPEAAVAETEIRQAA     120
                *******.****************************************************

g3867.t1        PELLLIAGLPEGVPNAAAKAASLFHRTGLVWLDLGRADLASACFEKATPLVCAADT--GR     178
g10502.t1       PELLLIAGLPEGVPNAAAKAASLFHRTGLVWLDLSRADLASACFEKATPLVCAADTEEDR     180
                **********************************.*********************  .*

g3867.t1        DILLDLNLARARTASSQGKHALAVALLSRSKPLAAASSQGFKALAEPYLLLGKAALATRS     238
g10502.t1       DILLDLNLARARTASSQGKHALAVGLLSRSKPLAAASSEGFKALAEAYLLLGKAALATKS     240
                ************************.*************:******* ***********:*

g3867.t1        PDPAIDASSLLTEALDLCEKAAASPCCATPTTPRSTPATTKLQVIKDQCLRFLAAERLEA     298
g10502.t1       PDPAIDASSLLTEALDLCEKAAASPCCATPTTPRSTPATTKLQLIKDQCLRFLAAERLEA     300
                *******************************************:****************

g3867.t1        NDYEGTLQCTRASRASPGLGKEHSSIAFMALRACLSSGKLVDAKRELGRLMANQEAPEFL     358
g10502.t1       NDYEGTLQCTRASRASPGLGKEHSSIAFMALRACLSSGKLVDAERELGRLMANEEAPEFL     360
                *******************************************:*********:******

g3867.t1        CVSAAELYLASAGLDAALKVLVALAARCRASAAAAAVRVLKTVVQGAGGGAGLARAIAEL     418
```

```
g10502.t1      CVSAAELYLASAGLDAALKVLVALASRCRASAAAAAVRVLKTVVQGAGGGAGRARAIAEL      420
               *************************:************************** *******

g3867.t1       VSDERVVALFNGTANTHERDTMHALLWTCGSEHFHAKNCEIAADLIERSMLYVSRDEESR      478
g10502.t1      VSDERVVALFNGPANTHERDTMHALLWTCGSEHFHAKNCEIAADLIERSMLYVSRDEESR      480
               ************ ***********************************************

g3867.t1       SRRAKCFRVLCLCHMALRHLDRAQEFITEAEKVEPNIHCAFLKFKILLHKKEDDEAIKLM      538
g10502.t1      SRRAKCFRVLCLCHMALRHLDRAQEFITEAEKVEPNIHCAFLKFKILLHKKDDDEAIKLM      540
               ***************************************************:********

g3867.t1       KTMVGYVDFNPHFLALSIHEAIGCKSFRVAVASLTFFLGLYSVGKPMPMGEAAVHRNLIA      598
g10502.t1      KTMVGYVDFNPHFLALSIHEAIGCKSFRVAVASLTFFLGLYSVGKPMPMSEAAVHRNLIA      600
               *************************************************.**********

g3867.t1       LLLLEPGSEAEILKYSRRAKLLMDELGVETFLGKGTVGLRELNWFAVSSWNMALKVVKEK      658
g10502.t1      LLLREPGSDAEILKYSRRAKLRMDELGVETFLGKGTVGLRELNWFAVSSWNMALKVVKEK      660
               *** ****:************ **************************************

g3867.t1       KYDYSSEFFELAAEFFSSESDLKKGIEMLRRAGKLLPLTSSSAPVTSDPLENNLPFLHTF      718
g10502.t1      KYDYSSEFFELAAEFFSSESDLKKGIEMLRRAGKLLPLTLPSAPVTSDPLENNLPFLHTF      720
               ***************************************  *******************

g3867.t1       NFYQLLNRLDTSAHPQQLQLVKSFAASKAYTPDHLLILGNMASEGTQPNLQVAEFVLKAS      778
g10502.t1      NFYQLLNRLDTSAHPQQLQLVKSFAASKACTPDHLLILGNMASEGTLPNLQVAEFLLKAS      780
               ***************************** **************** ********:****

g3867.t1       ISTALASHSPNYGVISAALRKLVYLSGLQDFSGSMSDAAYDVFQQAYQIVVGLRDGEYPF      838
g10502.t1      ISTALASHSPNYGVISAALRKLVYLSGLQDFSGSMSDAAYDVFQQAYQIVVGLRDGEYPF      840
               ************************************************************

g3867.t1       EEGRWLAITAWNKSYLPGQIGQHSVAKKWMKMGLDLARHFDRMKLYIPGMEECFENFQKL      898
g10502.t1      EEGRWLAITAWNKSYLPGRIGQHSVAKKWMKMGLDLARHFDRMKLYIPGMEECFENFQKL      900
               ******************:*****************************************

g3867.t1       SGKEPYERSQQDGEPSTSMSGTGSMSQPVLV*      929
g10502.t1      SGKEPDECSQQDGEPSTSMSGTGSMSQPVLV*      931
               ***** * ************************
```

protein argonaute MEL1 – 96.38% identity

```
g48565_B      MASRGRG-GGGGQGAGGGRRGEGRGRGVGGRGGYPQPYGRGEHGGGEPGGRGGG------      53
g9296_A       MASRGRGSGGGGQGSGGGRGGEGRGRGVGGRGGYPQPYGRGEHGGGEPGGRGGGMGRGRG      60
              ******* ******:**** **********************************

g48565_B      -----------------MGYQQPPPPPVGNVEGGGGRGRGGVAAAPARPAAPAPRPQAPP      96
g9296_A       IGGRGDGGGRGTGGRGGVGYQQPPPPPLGNVEGGGGRGRGGVAAAPARPAAPAPRPQAPP      120
                               :*********:********************************

g48565_B      VAAPAFPAAASSSAPSPTPAQAPGAPAGAAPVAPLAAGMGRLAVADSNPRPAVPPAPAAV      156
g9296_A       VAAPAFPAAASSSAPSPTPAQAPRAPAGAAPVAALAAGMGRLAVADNNPRPAAPPAPAAV      180
              *********************** ********* ************.*****.*******

g48565_B      RSEAQAAAPARQPPQAPPLSSKGIAPPARPGFGTSGRKVLVRANHFAVQVADNDICHYDV      216
g9296_A       RSEAQAAAPARQPPQAPPLSSKGIAPPARPGLGTSGRKLLVRANHFAVQVADNDICHYDV      240
              *******************************:******:*********************

g48565_B      LINPEPKARRINRVILSELVKVHGATSLARKIPAYDGSKSLYTAGELPFKSMEFVVKLGR      276
g9296_A       LINPEPKARRINRVILSELVKVHGATSLARKIPAYDGSKSLYTAGELPFKSMEFVVKLGR      300
              ************************************************************

g48565_B      REIEYKVTIRYAARPNLFHLKQFLKGQQRDAPYDTIQALDVALRESPSLNYVTLSRSFFS      336
g9296_A       REIEYKVTIRYAARPNLFHLKQFLKGQQRDAPYDTIQALDVALRESPSLNYVTLSRSFFS      360
              ************************************************************

g48565_B      KNFGEVKDIGGGLECWRGYYQSLRPTQMGLSLNIDICSTSFYQSISVVKFVGDCLRLTNP      396
g9296_A       KNFGEGEDIGGGLECWRGYYQSLRPTQMGLSLNIDICSTSFYQSIPVVKFVSDCLRLTNP      420
              ***** :************************************** *****.********
```

```
g48565_B      AQPFPNRDRLKLKKALRGVRVETTHQQGKRSIYKITGITPVPLTQLSFSCEEGPQLTVVQ      456
g9296_A       AQPFSDRDRLKLKKALRGVRVETTHQQGKRSIYKITGITPVPLTQLSFSCEEGPQLTVVQ      480
              **** :******************************************************

g48565_B      YFARRYNYRLHYTAWPCLQSGNDSKPIYLPMEVCQIIEGQRYPRKLSDTQVANILKATCK      516
g9296_A       YFAQRYNYRLRYTSWPCLQSGNDSKPIYLPMEVCQIIEGQRYPRKLSDTQVANILKATCK      540
              ***:******:**:**********************************************

g48565_B      PPQEREDSIIKMVRQNNYSADRMAQVFGITVANQMANVQARVLPAPTLKYHESGKEKTVA      576
g9296_A       RPQEREDSIIKMVRHNNYSADKMAQVFGITVANQMANVQARVLPAPMLKYHESGKEKTVA      600
               *************:******:************************ *************

g48565_B      PSLGQWNMINKKMVNGGTVDSWTCLSFSRIPLHEVNRICEDLAQMCNSIGMRFNPRPVTE      636
g9296_A       PSLGQWNMINKKMVNGGTVDSWTCLSFSRIPLHEVNRICEDLAQMCNSIGMRFNPRPVTE      660
              ************************************************************

g48565_B      VKSASPNHIEGALRDVHTRAPNLQLLIVILPDVSGHYGTIKRICETDIGIVSQCMNPKKN      696
g9296_A       VKSASPNHIEAALRDVHMRAPNLQLLIVILPDVSGHYGTIKRICETDLGIVSQCMNPKKN      720
              **********.****** *****************************:************

g48565_B      KNKQYFENVALKVNVKVGGCNTVLERALVKNGIPFVNDVPTIIFGADVTHPTAGEDSSAS      756
g9296_A       KNKQYFENVALKVNVKVGGRNTVLERALVPNGIPYVTDVPTIIFGADVTHPTAGEDSSAS      780
              ******************* ********* ****:*.***********************

g48565_B      IAAVVASMDWPQVTTYKALVSAQAHREEIIQNLFWTATDPEKGTPVNGGMIRELMISFFR      816
g9296_A       IAAVVASMDWPQVTTYKALVSAQAHREEIIQNLFWTATDPEKGTPVNGGMIRELMTSFFR      840
              ******************************************************* ****

g48565_B      RTARKPRRIIFYRDGVSEGQFSHVLLHEMDAIRKACASMEDGYLPPVTFVVVQKRHHTRL      876
g9296_A       RTGRKPRRIIFYRDGVSEGQFSHVLLHEMDAIRKACASMEDRYQPPVTFVVVQKRHHTRL      900
              **.************************************** * ****************

g48565_B      FPEVHGRRDLTDNSGNILPGTVVDTSICHPSQFDFYLCSHAGIKGTSRPTHYHVLYDEND      936
g9296_A       FPEVHGRRDLTDKSGNILPGTVVDTSICHPSEFDFYLCSHAGIKGTSRPTHYHVLYDENH      960
              ************:******************:***************************.

g48565_B      FSADGLQMLTNSLCYTYARCTRAVSVVPPAYYAHLAAFRARYYDEQGSTDGASVVSGGAA      996
g9296_A       FSADGLQMLTNSLCYTYARCTRAVSVVPPAYYAHLAAFRARYYDEQGSTDGTSVVSGGAA      1020
              ***************************************************:********

g48565_B      AAGGGAPAFRRLPQIKENVKDVMFFC*      1022
g9296_A       AAGGGAPAFRRLPQIKENVKDVMFFC*      1046
              ***************************
```

mediator of RNA polymerase II transcription subunit 23 – 99,75% identity

```
g56330_B      MDGGHGARQPMSPAISASAVLPQQRQMQPHHHPARTAIADLFTLYLGMNSKQRAEDPMRE      60
g44089_A      MDGGHGARQPMSPAISASAVLPQQRQMQPHHHPARTAIADLFTLYLGMNSKQRAEDPMRE      60
              ************************************************************

g56330_B      SPNKLQKRVTALNRDLPPRDEQFISDYEQLRMPFPDAEQLQAVTESVLISFVLQCSSHAP      120
g44089_A      SPNKLQKRVTALNRDLPPRDEQFISDYEQLRMPFPDAEQLQAVTESVLISFVLQCSSHAP      120
              ************************************************************

g56330_B      QSEFLLFATRCLCARGHLRWDSLLPALLNTVSSIEAPMVQGVSVTGAGPATPSSAIAMPN      180
g44089_A      QSEFLLFATRCLCARGHLRWDSLLPALLNTVSSIEAPMVQGVSVTGAGPATPSSAIAMPN      180
              ************************************************************

g56330_B      APNFHPSNPASPLSVMNTIGSPTQSGIDQPVGANVSPIKAAEFSSSAQLGTAARGDQSRR      240
g44089_A      APNFHPSNPASPLSVMNTIGSPTQSGIDQPVGANVSPIKAAEFSSSAQLGTAARGDQSRR      240
              ************************************************************

g56330_B      GAEASYLHHLSCRIILAGLEFNLKPATHAVIFQHMVNWLVNWDQRPHGMDEADVMQTCRL      300
g44089_A      GAEASYLHHLSCRIILAGLEFNLKPATHAVIFQHMVNWLVNWDQRPHGMDEADAMQTCRL      300
              ****************************************************.*******

g56330_B      EKPLHEWMHLCLDVIWILVNEDKCRIPFYELVRCNLQFLENIPDDEALVSIIMEIHRRRD      360
```

```
g44089_A      EKPLHEWMHLCLDVIWILVNEDKCRIPFYELVRCNLQFLENIPDDEALVSIIMEIHRRRD     360
              ************************************************************

g56330_B      MVCMHMQMLDQHLHCPTFGTHRFLSQSYPSIAGESVTNLRYSPITYPSVLGEPLHGEDLA     420
g44089_A      MVCMHMQMLDQHLHCPTFGTHRFLSQSYPSIAGESVTNLRYSPITYPSVLGEPLHGEDLA     420
              ************************************************************

g56330_B      NSIPKGGLDWERALRCLRHALRTTPSPDWWRRVLLVAPCYRSQSQQSSTPGAVFSPDMIG     480
g44089_A      NSIPKGGLDWERALRCLRHALRTTPSPDWWRRVLLVAPCYRSQSQQSSTPGAVFSPDMIG     480
              ************************************************************

g56330_B      EAVADRTIELLRLTNSETQSWQDWLLFADIFFFLMKSGCIDFLDFVDKLASRVTSSDQQI     540
g44089_A      EAVADRTIELLRLTNSETQSWQDWLLFADIFFFLMKSGCIDFLDFVDKLASRVTSSDQQI     540
              ************************************************************

g56330_B      LRSNHVTWLLAQIIRIEIVMNTLSSDPRKVETTRKIISFHKEDKSLEANNIGPQSILLDF     600
g44089_A      LRSNHVTWLLAQIIRIEIVMNTLSSDPRKVETTRKIISFHKEDKSLEANNIGPQSILLDF     600
              ************************************************************

g56330_B      ISSSQTLRIWSFNTSIREHLNSDQLQKGKQIDEWWKQMTKASGERMIDFMNLDERETGMY     660
g44089_A      ISSSQTLRIWSFNTSIREHLNSDQLQKGKQIDEWWKQMTKASGERMIDFMNLDERATGMF     660
              ******************************************************* ***:

g56330_B      WVLSFTMAQPACEAVMNWFTSAGMADLIQGPNMQPSERIMMMRETYPLSMSLLSGLSINL     720
g44089_A      WVLSFTMAQPACEAVMNWFTSAGMADLIQGPNMQPSERIMMMRETYPLSMSLLSGLSINL     720
              ************************************************************

g56330_B      CLKLAFQLEETIFLGQAVPSIAMVETYVRLLLIAPHSLFRPHFTTLTQRSPSILSKSGVS     780
g44089_A      CLKLAFQLEETIFLGQAVPSIAMVETYVRLLLIAPHSLFRPHFTTLTQRSPSILSKSGVS     780
              ************************************************************

g56330_B      LLLLEILNYRLLPLYRYHGKSKALMYDVTKIISMIKGKRGEHRLFRLAENLCMNLILSLK     840
g44089_A      LLLLEILNYRLLPLYRYHGKSKALMYDVTKIISMIKGKRGEHRLFRLAENLCMNLILSLK     840
              ************************************************************

g56330_B      DFFFVKKELKGPTEFTETLNRITIISLAITIKTRGIAEVEHMIYLQPLLEQIMATSQHTW     900
g44089_A      DFFFVKKELKGPTEFTETLNRITIISLAITIKTRGIAEVEHMIYLQPLLEQIMATSQHTW     900
              ************************************************************

g56330_B      SEKTLRYFPPLIRDFLMGRMDKRGQAIQAWQQAETTVINQCNQLLSPSAEPNYVMTYLSH     960
g44089_A      SEKTLRYFPPLIRDFLMGRMDKRGQAIQAWQQAETTVINQCNQLLSPSAEPNYVMTYLSH     960
              ************************************************************

g56330_B      SFPQHRQYLCAGAWMLMNGHLEINSANLARVLREFSPEEVTANIYTMVDVLLHHIQFEVQ     1020
g44089_A      SFPQHRQYLCAGAWMLMNGHLEINSANLARVLREFSPEEVTANIYTMVDVLLHHIQFEVQ     1020
              ************************************************************

g56330_B      RGHLAQDLLSKAITNLSFFIWTHELLPLDILLLALIDRDDDPYALRLVISLLEKPELQQR     1080
g44089_A      RGHLAQDLLSKAITNLSFFIWTHELLPLDILLLALIDRDDDPYALRLVISLLEKPELQQR     1080
              ************************************************************

g56330_B      VKNFCNTRSPEHWLKNQHPKRAELQKALGSHLSWKDRYPPFFDDIAARLLPVIPLIIYRL     1140
g44089_A      VKNFCNTRSPEHWLKNQHPKRAELQKALGSHLSWKDRYPPFFDDIAARLLPVIPLIIYRL     1140
              ************************************************************

g56330_B      IENDATDIADRVLAFYSSLLAFHPLRFTFVRDILAYFYGHLPIKLIGRILNLLGVSTKTP     1200
g44089_A      IENDATDIADRVLAFYSSLLAFHPLRFTFVRDILAYFYGHLPIKLIGRILNLLGVSTKTP     1200
              ************************************************************

g56330_B      FSESFAKYLVSSNSSICPPPEYFANLLLNLVNNVIPPLSSKSKSNPADTTRSTFNKHHAS     1260
g44089_A      FSESFAKYLVSSNSSICPPPEYFANLLLNLVNNVIPPLSSKSKSNPADTTRSTFNKHHAS     1260
              ************************************************************

g56330_B      SQPGGIGNTDGQRAFYQNQDPGSYTQLVLETAAIEILSLPVPAAQIVSSLVQIIAHVQAM     1320
g44089_A      SQPGGIGNTDGQRAFYQNQDPGSYTQLVLETAAIEILSLPVPAAQIVSSLVQIIAHVQAM     1320
              ************************************************************

g56330_B      LIQSNSGQGMSGGLGQSSGLPTSPSGAAESSGPNQANSAASGINATNFISRSGYSCQQLS     1380
```

```
g44089_A      LIQSNSGQGMSGGLGQSSGLPTSPSGAAESSGPNQANSAASGINATNFISRSGYSCQQLS      1380
              ************************************************************

g56330_B      VLMIQACGLLLAQLPPEFHMQLYSEAARVIKDCWWLADSSRPVKELDSAVGYALLDPTWA      1440
g44089_A      VLMIQACGLLLAQLPPEFHMQLYSEAARVIKDCWWLADSSRPVKELDSAVGYALLDPTWA      1440
              ************************************************************

g56330_B      SQDNTSTAIGNTVALLHSFFSNLPQEWLESTHTVIKHLRPVNSVAMLRIAFRILGPLLPR      1500
g44089_A      SQDNTSTAIGNTVALLHSFFSNLPQEWLESTHTVIKHLRPVNSVAMLRIVFRILGPLLPR      1500
              *************************************************.**********

g56330_B      LAFARPLFMKTLALLFNVLGDVFGKNPPVPNPNPVEASEIADIIDFLHHAVMYEGQGGPV      1560
g44089_A      LAFARPLFMKTLALLFNVLGDVFGKNPPVPNPNPVEASEIADIIDFLHHAVMYEGQGGPV      1560
              ************************************************************

g56330_B      QSTSKPKLEILTLCGKVIEILRPDVQHLLSHLKIDPTSSIYAATHPKLVQNSS*      1613
g44089_A      QSTSKPKLEILTLCGKVIEILRPDVQHLLSHLKIDPTSSIYAATHPKLVQNSS*      1613
              ******************************************************
```

DUF724 domain-containing protein 7-like – 84,72% identity

```
g3237_B       MAAAASGSAPPPDAAGARRGRGRPRGSKNGSGRGRSRS----------------LRSPSG 44
g5789_A       MAAAASGSAPPPDAAGARRGRGRPRGSKNGSGRGRGRGRSRSLRWSKAPKRPRLSVSPSG 60
              ***********************************.*.                  ****

g3237_B       SPDGPHSPRPAVDHSAPLPPGTEVEVRVDDDGFHGSWFEATVLDFSPARGYCHPARYTVS 104
g5789_A       SPDGPHSPRPAVDLSAPLPPGTEVEVRVDDDGFHGSWFEATVLDFSPARGYRHPARYTVS 120
              ************* ************************************* ********

g3237_B       YVHLLADDDEGVLAEPFAPSHIRPRPPPPAPTDPPRFQTHDIVEAFHNEGWWSGIVVSAP 164
g5789_A       YVHLLADDDEGVLAEPFAPSHIRPRPPPPPPTDPPRFQTHDIVEAFHNEGWWSGIVMSAP 180
              ***************************** **************************:***

g3237_B       DSTDSPDGAGAAGVTVAFPITREVIEFAPGLVRPRRDYVDGEWIPSQVAMIVRPKRAVKV 224
g5789_A       DSTDSPDGAGAAGVTVAFPITREVLEFAPGLVRPRRDYVDGEWIPSQVAMIVRPKRAVKV 240
              ************************:***********************************

g3237_B       YEVGDKVEVVRQRNGYGESWFPATVRVVVDDLSYIVEYFDLEEG-EG-GQKATEYLHWRF 282
g5789_A       YEVGDKVEVVRQRNGYGESWFPATVRVVVDDLSYIVEYFDLEEEGEGGQEKATEYLHWRF 300
              *******************************************  **  :**********

g3237_B       IRPAVEHSPRESEFQLQPGAAVEAYCDGAWSLGVVRTVLGEGEYEIGIVAKKSEMLVTKV 342
g5789_A       IRPAVEHSPRESEFQLQPGAAVEAYCDGAWSPGVVRTVLGEGEYEIGIVAKKSEMLVTKV 360
              ******************************* ****************************

g3237_B       VPLLKPQYKWNGKQWRIATRKRRANTRRRSVSGNSPRSPVEVSSIDQEHSLGKNTLAE-- 400
g5789_A       VPLLKPQYKWNGKQWRIATRKRRANKRRRSVSGNSPRSPVEVSSSDQEHSLGKNTLAEGS 420
              *************************.****************** *************  

g3237_B       -------------------------------------------------------GHSGP 405
g5789_A       GHASVSEMDIPLSALCKSPESTRSPNSFVSEKNSPPGSHGIVNSVPMNGLVCASPGHSGP 480
                                                                     *****

g3237_B       VDNQEILSDMVVTDGELNGPVSGRSGDG-------------------------------- 433
g5789_A       VDNQEILSDMVVTDGELNGPVSGRSGDGNDMLSITELRKKMSSARRNSAVNRKQDNLAKS 540
              ****************************

g3237_B       ------ISNIKAGKTHSIQGLEGKIQLKGNMNFSAPDIVLALSVSGTGRTIISPDGLVSI 487
g5789_A       VSVKKCISNIKAGKTHSIQGLEGKIQLKGNMNFSAPDIVLALSVSGTGRTIISPDGLVSI 600
                    ******************************************************

g3237_B       GTERGSSTKVSVGTKRRSSTKVLACKKLANRRGSKELCSPNSSLDVTGTVQQRGRKEVAE 547
g5789_A       GTERGSSTKVSVGTKRGSSTKVLACKKLANRRGSKELCSPNSSLDVTGTVQQRGRKEVAE 660
              **************** *******************************************

g3237_B       PMEECPFALECPNSGTREQLDRTLEDAQNIIELSNPDLFLMVPPGFESMYNGKGINTNDT 607
g5789_A       PMEECPFALECPNSGTREQLDRTLEDAQNIIELSNPDLFPMVPPGFESMDNGKGINTNDA 720
              *************************************** ********* *********:
```

```
g3237_B      QFDEEPSGTTNSLIEPKGNGDMCTNHAATKLTESNHVMETAILSLDSPAQQAYGKVDERS 667
g5789_A      QFDEEPSGTTNSLIEPKGNGDMCTNHAATKLTESNHVMETAILSLDSPAQQACGKVDERS 780
             **************************************************** *******

g3237_B      VLQNARSSQCIINSSPLRSCPAFESLLPLPQPLSQVSKHQALFVKNSPLWHLVETMHVFK 727
g5789_A      VLQNARSSQCIINSSPLRSCPAFESLLPLPQPLSQVSKHPTLFVKNSPMWHLVETMHVFK 840
             *************************************** :*******:***********

g3237_B      ELPQQPHFLPLQEHLPMLREGMALGMMVSFADLVKITMEASMDNSMEWFKDKIKTISYLE 787
g5789_A      ELPQQPHFLPLQEHPPMLREGMALGMMVSFADLVKITMEASMDNSMEWFEDKIKTISYLE 900
             ************** **********************************:**********

g3237_B      ENGFNVQFIKSNMTELVKVKSELTSYYGLIGKLDSNFVEKTASSSRVGALLDEKDIAAAE 847
g5789_A      ENGFNVQFIKSNMTELVKVKSELTSYYGEIGKLDSKFVEKTASSSRVGALLDEKDIAAAE 960
             **************************** ******:************************

g3237_B      LEQELGRIRQESQRIAKEKEKIDAEVASIKTELSGYKDLCNGAESKAKDILARLRLKRLT 907
g5789_A      LEQELGRIRQESQRIAKEKEKIDAEVASIKTALSGYEDLCNGAESKAKDILARLRLKRLT 1020
             ******************************* ****:***********************

g3237_B      *      907
g5789_A      *      1020
             *
```

histone-lysine N-methyltransferase, H3 lysine-9 specific SUVH5-like, 100% identity

```
g52579.t1      MEMGAAAAAKPSPREDGVRRHKVLVPWRFQPGYFRQPLKHAAPANGVAPNGDGGYTAAGV      60
g10978.t1      MEMGAAAAAKPSPREDGVRRHKVLVPWRFQPGYFRQPLKHAAPANGVAPNGDGGYTAAGV      60
               ************************************************************

g52579.t1      PDVNNCGSDGAPSPVGEANGSRDAKIFGSDGAPSPGKSGAVGGGEEQSDRCTRGQSLKSP      120
g10978.t1      PDVNNCGSDGAPSPVGEANGSRDAKIFGSDGAPSPGKSGAVGGGEEQSDRCTRGQSLKSP      120
               ************************************************************

g52579.t1      GVDNGGRPEPGNACNLGDSVRGGWVKSSPLEGTGNSRGGANDGGEAVAGDDCNMGSSNCD      180
g10978.t1      GVDNGGRPEPGNACNLGDSVRGGWVKSSPLEGTGNSRGGANDGGEAVAGDDCNMGSSNCD      180
               ************************************************************

g52579.t1      GSLKDVGTQDFTGAGDGAACDPEVIEITEECFAKGSKKPFVDQTGSKSNGSASGLRSEDP      240
g10978.t1      GSLKDVGTQDFTGAGDGAACDPEVIEITEECFAKGSKKPFVDQTGSKSNGSASGLRSEDP      240
               ************************************************************

g52579.t1      EGNVGLGDSAYHAAKGCSMGDEAAKENDATAKGCSSATPGSNGNGTYVRKGRKVGVPWRF      300
g10978.t1      EGNVGLGDSAYHAAKGCSMGDEAAKENDATAKGCSSATPGSNGNGTYVRKGRKVGVPWRF      300
               ************************************************************

g52579.t1      QVGYKRSFSDAFDSNNGSPDRPAYKFDGSSTQCTPTTRSSVRCYASSHSGVRVSAMHNLS      360
g10978.t1      QVGYKRSFSDAFDSNNGSPDRPAYKFDGSSTQCTPTTRSSVRCYASSHSGVRVSAMHNLS      360
               ************************************************************

g52579.t1      MKGENGTGTECKKRKTDSDYQDEVMPNNGDPIVRESIMRSLQDLRLIYREILDEEEDNSR      420
g10978.t1      MKGENGTGTECKKRKTDSDYQDEVMPNNGDPIVRESIMRSLQDLRLIYREILDEEEDNSR      420
               ************************************************************

g52579.t1      EKVINNGADMRAYKIFRERFSTEFDDEKYIGSVPGIYPGDIFHLRVELCVVGLHRQHRLG      480
g10978.t1      EKVINNGADMRAYKIFRERFSTEFDDEKYIGSVPGIYPGDIFHLRVELCVVGLHRQHRLG      480
               ************************************************************

g52579.t1      IDCTKKDDGITVAVSIVSCAQSSDAKYNLDVLVYTGPVAVTVNQRIEGTNLALKKSMDTS      540
g10978.t1      IDCTKKDDGITVAVSIVSCAQSSDAKYNLDVLVYTGPVAVTVNQRIEGTNLALKKSMDTS      540
               ************************************************************

g52579.t1      TPVRVIHGFTTKNGKKKFPIYIYGGLYLVEKYWREKEHGDRYVYMFRLRRMKGQKHIEIQ      600
g10978.t1      TPVRVIHGFTTKNGKKKFPIYIYGGLYLVEKYWREKEHGDRYVYMFRLRRMKGQKHIEIQ      600
               ************************************************************

g52579.t1      EILQTGDSGSNDSVIIKDLSLGLERVPVPVVNKISDECPMPYRYTSHLQYPRNYRPTPPA      660
```

```
g10978.t1      EILQTGDSGSNDSVIIKDLSLGLERVPVPVVNKISDECPMPYRYTSHLQYPRNYRPTPPA     660
               ************************************************************

g52579.t1      GCGCVDGCSDSKKCACAMKNGGEIPFNDKGRILEAKPLVYECGPSCKCPPTCHNRVGQHG     720
g10978.t1      GCGCVDGCSDSKKCACAMKNGGEIPFNDKGRILEAKPLVYECGPSCKCPPTCHNRVGQHG     720
               ************************************************************

g52579.t1      LKFRLQIFKTKSMGWGVRTLDFIPSGSFVCEYIGEVLEDEEAQKRTNDEYLFAIGHNYYD     780
g10978.t1      LKFRLQIFKTKSMGWGVRTLDFIPSGSFVCEYIGEVLEDEEAQKRTNDEYLFAIGHNYYD     780
               ************************************************************

g52579.t1      ESLWEGLSRSIPSLQKGPGKDDETGFAVDASEMGNFAKFINHSCTPNIYAQNVLYDHEDI     840
g10978.t1      ESLWEGLSRSIPSLQKGPGKDDETGFAVDASEMGNFAKFINHSCTPNIYAQNVLYDHEDI     840
               ************************************************************

g52579.t1      SVPHIMFFACDDIRPNQELLYHYNYKIDQVHDANGNIKKKKCLCGSVECDGWLY*   894
g10978.t1      SVPHIMFFACDDIRPNQELLYHYNYKIDQVHDANGNIKKKKCLCGSVECDGWLY*   894
               *******************************************************
```

### tubulin beta-1 chain – 96.50% identity

```
g28522_B      MREILHIQGGQCGNQIGAKFWEVVCAEHGIDATGRYGGDSDLQLERVNVYYNEASCGRFV     60
g13706_A      MREILHIQGGQCGNQIGAKFWEVICDEHGIDHTGKYAGDSDLQLERINVYYNEASGGRYV     60
              ***********************:* ***** **:*.*********:******** **:*

g28522_B      PRAVLMDLEPGTMDSVRSGPYGHIFRPDNFVFGQSGAGNNWAKGHYTEGAELIDSVLDVV     120
g13706_A      PRAVLMDLEPGTMDSVRSGPYGQIFRPDNFVFGQSGAGNNWAKGHYTEGAELIDSVLDVV     120
              **********************:*************************************

g28522_B      RKEAENCDCLQGFQVCHSLGGGTGSGMGTLLISKIREEYPDRMMLTFSVFPSPKVSDTVV     180
g13706_A      RKEAENCDCLQGFQVCHSLGGGTGSGMGTLLISKIREEYPDRMMLTFSVFPSPKVSDTVV     180
              ************************************************************

g28522_B      EPYNATLSVHQLVENADECMVLDNEALYDICFRTLKLTTPSFGDLNHLISATMSGVTCCL     240
g13706_A      EPYNATLSVHQLVENADECMVLDNEALYDICFRTLKLATPTFGDLNHLISATMSGVTCCL     240
              *************************************:**:*******************

g28522_B      RFPGQLNSDLRKLAVNLIPFPRLHFFMVGFAPLTSRGSQQYRALTVPELTQQMWDAKNMM     300
g13706_A      RFPGQLNSDLRKLAVNLIPFPRLHFFMVGFAPLTSRGSQQYRALTVPELTQQMWDAKNMM     300
              ************************************************************

g28522_B      CAADPRHGRYLTASAMFRGKMSTKEVDEQMLNVQNKNSSYFVEWIPNNVKSTVCDIPPTG     360
g13706_A      CAADPRHGRYLTASAMFRGKMSTKEVDEQMLNVQNKNSSYFVEWIPNNVKSSVCDIPPTG     360
              ***************************************************:********

g28522_B      LKMASTFIGNSTSIQEMFRRVSEQFTAMFRRKAFLHWYTGEGMDEMEFTEAESNMKDLVF     420
g13706_A      LKMASTFIGNSTSIQEMFRRVSEQFTAMFRRKAFLHWYTGEGMDEMEFTEAESNMNDLVA     420
              *******************************************************:***

g28522_B      EYQVYQDAIVDEVGEYEDEEEADLQD*      446
g13706_A      EYQQYQDATADDEEEYEDEEEEEAA*-      445
              *** **** .*:  ******* :
```

### tubulin alpha-3 chain – 97.98% identity

```
g24105_A      MRECISVHIGQAGIQVGNACWELYCLEHGIQPDGQVPGDKTAGHHDDAFSTFFSQTGAGK     60
g8510_B       MRECISVHIGQAGIQVGNACWELYCLEHGIQPDGQVPGDETAGHHDDAFSTFFSQTGAGK     60
              ***************************************:********************

g24105_A      HVPRAIFVDLEPTVIDEVRTGTYRQLFHPEQLISGKEDAANNFARGHYTIGKEIVDLCLD     120
g8510_B       HVPRAIFVDLEPTVIDEVRTGTYRQLFHPEQLISGKEDAANNFARGHYTIGKEIVDLCLD     120
              ************************************************************

g24105_A      RIRKLADNCTGLQGFLVFNAVGGGTGSGLGSLLLERLSVEYGKKSKLGFTVYPSPQVSTS     180
g8510_B       HIRKLADNCTGLQGFLVFNAVGGGTGSGLGSLLLERLSVEYGKKSKLGFTVYPSPQVSTS     180
              :***********************************************************
```

```
g24105_A      VVEPYNSVLSTHSLLEHTDVSILLDNEAIYDICRRSLDIERPNYSNLNRLVSQVISSLTA      240
g8510_B       VVEPYNSVLSTHYLLEHTDVSILLDNEAIYDICRRSLDIERPNYSNLNRLVSQVISSLTT      240
              ************ **********************************************:

g24105_A      SLRFDGALNVDVNEFQTNLVPYPRIHFMLSSYAPVISSAKAFHEQLSVAEITSSAFEPAS      300
g8510_B       SLRFDGAINVDVNEFQTNLVPYPRIHFMLSSYAPVISSAKAFHEQLSVAEITSSAFEPAS      300
              *******:****************************************************

g24105_A      MMVKCDPRHGKYMACCLMYRGDVVPKDVNAAVSLIKTKRTIQFVDWCPTGFKCGINYQAP      360
g8510_B       MMVKCDPRQGKYMACCLMYRGDVVPKDVNAAVGLTKTKRTIQFVDWCPTGFKCGINYQAP      360
              ********:***********************.* *************************

g24105_A      TVVPGADLAKVQRAVCMISNSTSVVEVFSRINSKFDLMYAKRAFVHWYVGEGMEEGEFSE      420
g8510_B       TVVPGADLAKVQRAVCMISNSTSVVEVFSRINSKFDLMYAKRAFVHWYVGEGMEEGEFSD      420
              ***********************************************************:

g24105_A      AREDLAALEKDYEEVAAEGGGDEGEEDEEY*-----------------------------      450
g8510_B       AREDLAALEKDYEEVAAEGGGDEGEVVQRRRDALQCGDGLTVGINMVARSLRAGPGGYVL      480
              *************************  :.

g24105_A      ---------------   450
g8510_B       VVVGGRYVSGAAAAW*   495
```

### mRNA turnover protein 4 homolog – 83.78% identity

```
g35119_B      ------------------------------------------------------------      0
g16140_A      MPKSKRNRPVTLSKTKKKPGLERKGKVVAEIKDAIDRYSSAYVFTYDNMRNQKLKDLREQ      60

g35119_B      ------------------------------------------------------------      0
g16140_A      LKSSSRIFLAGKKVMQIALGRSPADEAKTGLHKLSKFLQGDSGLFFTNLPRDDVERMFRE      120

g35119_B      -----MKR---RSSKIVPMRWGERQIF-------VDFFCNPERLKNQVRELTSRVKALQN      45
g16140_A      FEEHDFARTGSTATETVELKEGPLEQFTHEMEPFLRKQGLPVRLNKGV------------      168
                   : *    ::: * :: *  : *       :     * **:: *

g35119_B      CSCTLVLHVIHFLCIKFNFTPTSSWGRCAGTRLLGVQMVTLRLNLVCCWSCDDFKVYKEG      105
g16140_A      ----IELVADHVVCEEG---KPLSPEAAQTLRLLGIQMATFRLYLVCRWSCDDFEVYKEG      221
                  : * . *.:* :       *   .   ****:**.*:** *** ******:*****

g35119_B      LMHLGADDFLLSLCFLYTRQHFAVLTRTVFGSLCVAM* 142
g16140_A      LMHLGADDSS*--------------------------- 231
              ********
```

### ATP-dependent RNA helicase DEAH11 – 98.91% identity

```
g1204_B       MRRSQDRGLLRPPDWVPRPPP--HHRDHHYHNEHRYPPHSHPHRDRHYSAERRYQPRAQQ      58
g46421_A      MRRSQDRGLLRPPDWVPRPPPQHHHRDHHYHNEHRHPPHSHPHRDRHYSAERRYQPRAQQ      60
              *********************  ************:************************

g1204_B       PSPPPSQFEVLLVRPGPDLSAPTAIEVEGLVAGLPSPPPASVSVHSSGRHAARLVFASVS      118
g46421_A      PSPPPSQFEVLLVRPGPDLSAPTAIEVEALVAGLPPPAPASVSVHSSGRHAARLAFASVS      120
              ****************************.****** * ****************.*****

g1204_B       DAAAAAARQLWALRLEGLHLLALDLPHAAVAAHAKPLIASLFSDHASRLLDSDLVAVSAAR      178
g46421_A      AAAAAAARQLWALRLEGLHLLALDLPHAAVAAHAKPLIASLFADHASRLLDSDLVAVSAAR      180
               ****************************************:******************

g1204_B       SADLAASIRDVKRRLAGRNRVRDFHQLDLEKKTLESEKELVDAKIAEYKEAMLSIQRAML      238
g46421_A      SADLAASIRDAKRRLGGRNRVRDFHQLDLEKKTLESEKELVDAKIAEYKEAMLSIQRAML      240
              **********.****.********************************************
```

```
g1204_B         RRSGDKEEGVHLFGAVEGADVDFVRVHMMLLRECRRLKEGLPIYAYRRKILNHILANQAM    298
g46421_A        RGSGDKEEGVHLFGAVEGADVDFVRVHMMLLRECRRLKEGLPIYAYRRRILNHILANQAM    300
                * **********************************************:***********

g1204_B         VLIGETGSGKSTQLVQFLADSGLAGGRSIVCTQPRKLAAISLAHRVDEESKGCYGDSSVM    358
g46421_A        VLIGETGSGKSTQLVQFLADSGLAGGRSIVCTQPRKLAAISLAHRVDEESKGCYGDSSVM    360
                ************************************************************

g1204_B         SYSTLLNSQGFGTKIIFTTDSCLLHNCMSDMSLDGISYVIIDEAHERSLNTDLLLAMIKK    418
g46421_A        SYSTLLNSQGFGTKIIFTTDSCLLHNCMSDMSLDGISYVIIDEAHERSLNTDLLLAMIKK    420
                ************************************************************

g1204_B         KLLDRLDLRLIIMSATADADRLAEYFFGCQTFHVKGRTFPVEIKYVPDISAEASLNSIPS    478
g46421_A        KLLDRLDLRLIIMSATADADRLAEYFFGCQTFHVKGRTFPVEIKYVPDISAEASLNSVPS    480
                *********************************************************:**

g1204_B         MSSVASAAPSYVTDVVQMVNIIHKNEEEGAILAFLTSQLEVEWACETFSDPNAVVLPMHG    538
g46421_A        MSSVASAAPSYVTDVVQMVNIIHKNEEEGAILAFLTSQLEVEWACETFSDPNAVVLPMHG    540
                ************************************************************

g1204_B         KLSSIEQNLVFQSYPGKRKIIFCTNIAETSLTIKEVKYVVDCGLAKEYRFVPSSGLNVLK    598
g46421_A        KLSSIEQNLVFQSYPGKRKIIFCTNIAETSLTIKEVKYVVDCGLAKEYRFVPSSGLNVLK    600
                ************************************************************

g1204_B         VNWISQSSANQRAGRAGRTGAGKCYRLYPESDFGLMEAHQEPEIRKVHLGTAVLRILALG    658
g46421_A        VNWISQSSANQRAGRAGRTGAGKCYRLYPESDFGLMEAHQEPEIRKVHLGTAVLRILALG    660
                ************************************************************

g1204_B         VPDVKYFEFVDAPDPEAINMAVHNLEQLGAIKYKCSGFELTDTGRDLVKLGIEPRLGKIM    718
g46421_A        VPDVKYFEFIDAPDPEAINMAVHNLEQLGAIKYKCSGFELTDTGRDLVKLGIEPRLGKIM    720
                *********:**************************************************

g1204_B         LDCFSYGLMKEGLVLASVMANASSIFCRVGTNEEKYKADRLKVPFCHPDGDLFTSLAVYK    778
g46421_A        LDCFSYGLMKEGLVLASVMANASSIFCRVGTNEEKYKADRLKVPFCHPDGDLFTSLAVYK    780
                ************************************************************

g1204_B         KWEAGPDNKNMWCWQNSINAKTLRRCQETISELEKCLKHELNTIVPSYWSWNPEKPTMHD    838
g46421_A        KWEAGPDNKNMWCWQNSINAKTLRRCQETISELEKCLKHELNTIVPSYWSWNPEKPTMHD    840
                ************************************************************

g1204_B         TTLKKIILSSLRGNLAMFSGHEKFGYQVISADQPVQLHPSCSLLTYGSKPEWVVFSEILS    898
g46421_A        TTLKKIILSSLRGNLAMFSGHEKFGYQVISADQPVQLHPSCSLLTYGSKPEWVVFSEILS    900
                ************************************************************

g1204_B         VPNQYLVCVTAVDRNEVCTVNSMSFIEQVEESKLQRKVITGIGNKSLRRFCGKSGQNLQK    958
g46421_A        VPNQYLVCVTAVDRNEVCTVNSMSFIEQVEESKLQRKVITGIGNKSLRRFCGKSGQNLQK    960
                ************************************************************

g1204_B         IVSLLREDCRDDRIMVDLDFSSSEVLLFAKEHDMETVFCVVNDALELEAKMLSDECDERR    1018
g46421_A        IVSLLREDCRDDRIMVDLDFSSSEVLLFAKEHDMETVFCVVNDALELEAKMLSDECDERR    1020
                ************************************************************

g1204_B         PGGSTIALFGSGAEIKHLELGKRCLTVEIMHQNARDIDEKELIGLVYSHVPGIANFHKLG    1078
g46421_A        PGGSTIALFGSGAEIKHLELGKRCLTVEILHQNARDIDEKELIGLVYSHVPGIANFHKLG    1080
                *****************************:******************************

g1204_B         NFQSNTDETKWGRFTFLKPDYADDAISKLNGIEFHGSSLKVGHVSTYNHPGLPFPAVRAK    1138
g46421_A        NFQSNTDETKWGRFTFLKPDYADDAISKLNGIEFHGSSLKVGHVSTYNHPGLPFPAVRAK    1140
                ************************************************************

g1204_B         VSWPRKPSRGLALVTCASGEAEFIVKDCFALGVGGRYINCEVSKKFANCVFVRGVPLHVT    1198
g46421_A        VSWPRKPSRGLALVTCASGEAEFIVKDCFALGVGGRYINCEVSKKFANCVFVRGVPLHVT    1200
                ************************************************************

g1204_B         EPELYDAFRSTTTRRILDIRLLRGAPTASSSDSECAEALMRAISLFMPNRNFPGQNFRVH    1258
g46421_A        EPELYDAFRSTTTRRILDIRLLRGAPTASSSDSECAEALMRAISLFMPNRNFPGQNFRVH    1260
                ************************************************************
```

```
g1204_B        VIPPEEKDSMMRATITFDGSFHREAARALDHLQGSVLPCCLPWQIIQCQHVFHSTVSCPM      1318
g46421_A       VIPPEEKDSMMRATITFDGSFHREAARALDHLQGSVLPCCLPWQIIQCQHVFHSTVSCPM      1320
               ************************************************************

g1204_B        RIYNVISQEVGVLLESFRSEKGVSYNLEKNENGHFRVKLTANATKTIADLRRPLELLMKG      1378
g46421_A       RIYNVISQEVGVLLESFRSEKGVSYNLEKNENGHFRVKLTANATKTIADLRRPLELLMKG      1380
               ************************************************************

g1204_B        KIINHPDLMLSTVQLLWSRDGMEHLKSVEQETGTYILYDRQSRNIKVFGSTDKVAAAEEK      1438
g46421_A       KIINHPDLMLSTVQLLWSRDGMEHLKSVEQETGTYILYDRQSRNIKVFGSTDKVAAAEEK      1440
               ************************************************************

g1204_B        LVRALVQLHEKKPLEVCLRGRNLPPNLMKEVIKKFGADLEGLKTEVPAVDLQLNTRKQTL      1498
g46421_A       LVRALVQLHEKKPLEVCLRGRNLPPNLMKEVIKKFGADLEGLKSEVPAVDLQLNTRKQTL      1500
               *******************************************:****************

g1204_B        YVRGSKEDKQRVEEMISELIASSDHNAPLPSKNACPICLCELEDPFKLESCGHMFCFACL      1558
g46421_A       YVRGSKEDKQRVEEMISELIASSDHNAPLPSKNACPICLCELEDPFKLESCGHMFCFACL      1560
               ************************************************************

g1204_B        VDQCESAMKSQGGFPLCCLKNGCKNLLLLADLRSLVPDKLDELFRASLNAFVASSAGVYR      1618
g46421_A       VDQCESAMKSQDGFPLCCLKNGCKNLLLLADLRSLVPDKLDELFRASLNAFVASSAGLYR      1620
               ***********.*********************************************:**

g1204_B        FCPTPDCTSIYQVGAAGAEDKPFVCGACSVETCTKCHLEYHPFISCEAYKEYKADPTDAT      1678
g46421_A       FCPTPDCTSIYQVGAAGAEDKPFVCGACSVETCTKCHLEYHPFISCEAYKEYKADPTDAT      1680
               ************************************************************

g1204_B        LLEWRKGKENVKNCPSCGYTIEKAEGCNHVECRCGSHICWNCLESFKSSEECYGHLRSVH      1738
g46421_A       LLEWRKGKENVKNCPSCGYTIEKAEGCNHVECRCGSHICWNCLESFKSSEECYGHLRSVH      1740
               ************************************************************

g1204_B        LAYV* 1742
g46421_A       LAYV* 1744
               *****
```

CENH3 *Aegilops speltoides*

```
Aeg_CENH3_A_g18195      MARTKHPAVRKTKA-----PPKKQLGPRPAQRRQETDGAGTSATPRRAGRAAAPG--AAE   53
Aeg_CENH3_B_g94071      MTRTKKPPVSKLKMTRTKQPPVSKLKVRA------A-----------DGSARSPGGTQQT   43
                        *:***:* * * *      ** .:*  *       :            * * :**

Aeg_CENH3_A_g18195      GATGQPKQRKPHRFRPGTVALREIRKYQKSVDFLIPFAPFVRLIKEVTDFFCPEISRWTP   113
Aeg_CENH3_B_g94071      AASGQPRQRKPHRFRPGTVAQREIRKYQKSVDLLIPLAPFVRLIKEITNDFREG-IRFTP   102
                        .*:***:************* ***********:***:*********:*: *     *:**

Aeg_CENH3_A_g18195      QALVAIQEAAEYHLVDVFERANHCAIHAKRVTVMQKDIQLARRIGGRRLW*     163
Aeg_CENH3_B_g94071      GALMTIQEAAEYHLVDEFERANHCANHAKRVTVTLKDIELARLIGGRRLW*     152
                         **::*********** ******** *******  ***:*** ********
```

CENH3 *Zea mays*

```
Zea_mays_CenH3_A      MARTKHQAVRKTAEKPKKKLQFERSGGASTSATPERAAGTGGRAASGGDSVKKTKPRHRW     60
Zea_mays_CenH3_B      MARTKHQAVRKPAEKPKKKLQFERS-------------------------VKKTKPRHRW     35
                      *********** *************                         **********

Zea_mays_CenH3_A      RPGTVALREIRKYQKSTEPLIPFAPFVRVVRELTNFVTNGKVERYTAEALLALQEAAEFH     120
Zea_mays_CenH3_B      RPGTVALREIRKYQKSTEPLIPFAPFVRVVRELTNFLTNGKVERYTPEALLALQEAAEFH     95
                      ************************************:********* *************

Zea_mays_CenH3_A      LIELFEMANLCAIHAKRVTIMQKDIQLARRIGGRRWA 157
Zea_mays_CenH3_B      LIELFEIANLCAIHAKRVTVMHKDIQLARRIGGRRWA 132
                      ******:************:*:***************
```
