## Additional file 11 for "Sorghum embryos undergoing B chromosome elimination express B-variants of mitotic-related genes"

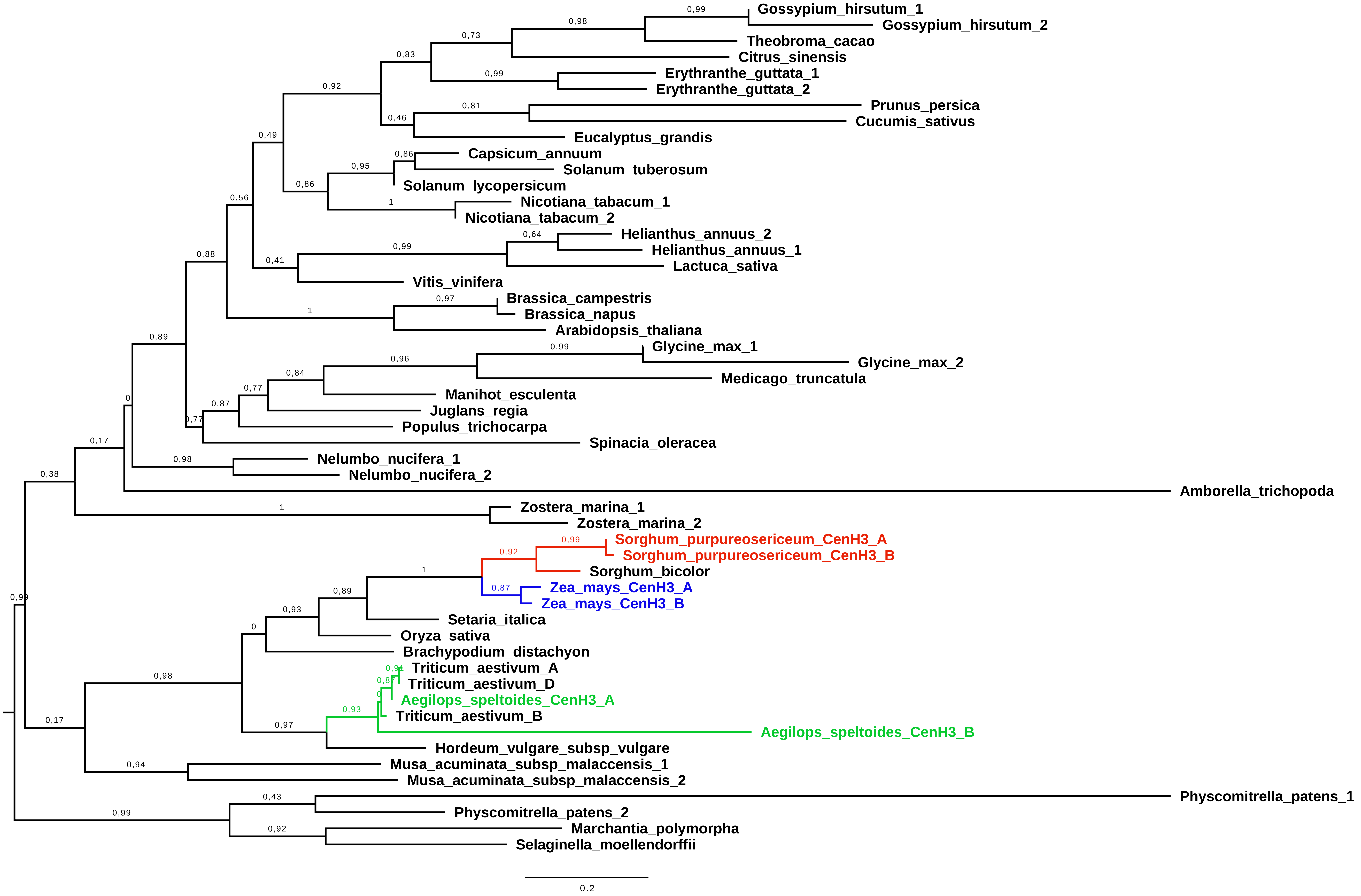


Additional File 11: Complete phylogenetic tree of CENH3 Proteins. Proteins from species with B chromosomes are highlighted in red (Sorghum purpureosericeum), green (Aegilops speltoides), and blue (Zea mays). Only protein sequences belonging to the PANTHER subfamilies PTHR11426:SF223 and PTHR11426:SF277 were included in the analysis.
