## Additional file 12 for "Sorghum embryos undergoing B chromosome elimination express B-variants of mitotic-related genes"

Additional file 12: Protocol for High-Molecular-Weight DNA Isolation from Flow-Sorted Nuclei for ONT Sequencing

Nuclei were isolated from *Sorghum purpureosericeum* young inflorescence tissue. Initially, young inflorescence leaves were sectioned and wrapped in nylon mesh packages and fixed in a PBS solution containing 2% formaldehyde under vacuum for 20 minutes. Post-fixation, these packages were rinsed with running water and subsequently washed three times in the TRIS buffer. A fraction of the leaves was then placed in a Petri dish with 1 ml of either HKS or LB01 buffer and sectioned using a razor blade. The resulting buffer, now containing the nuclei, was filtered through nylon mesh into cytological tubes. The nuclei were then flow-sorted into Protein LoBind tubes, each containing 100 µl ET and 6 µl EDTA, targeting a sorting density of 200 000 nuclei per tube, with a DNA content not exceeding 1 µg.

For DNA isolation, 17 µl of 10% SDS and 17 µl of proteinase K (20 mg/ml) were added to each tube containing the sorted nuclei, followed by incubation at 50 °C for 20 hours. The lysate was then transferred to Microcon Fast-flow columns and filled to maximum capacity with 5 mM Tris-HCl. Centrifugation was performed at 500 g for 6-15 minutes at 26 °C, and this step was repeated 7-9 times to ensure maximum sample throughput, with adjustments made for sample-specific flow rates. The final centrifugation step concentrated each sample to a volume of 50-100 µl, after which the column was inverted into a new tube and centrifuged at 1,000 g for 1 minute to collect the purified DNA. DNA concentration and purity were assessed using a NanoDrop spectrophotometer, with samples undergoing AMPure purification if necessary for enhanced purity.

Given the tendency for *Sorghum purpureosericeum* DNA to form insoluble aggregates with AMPure beads, a pre-treatment involving mechanical fragmentation was employed using a syringe to shear the longest DNA molecules. AMPure beads were equilibrated to room temperature and vortexed before being added to the sample in a 1:1 ratio. The mixture was incubated at 37 °C for 10 minutes, with intermittent tapping post-incubation for an additional 5 minutes at room temperature. Following magnetic separation, the supernatant was discarded, and the beads were washed with 70% ethanol twice, allowing the beads to dry for 30-60 seconds before re-suspension in 30 µl of nuclease-free water. This suspension was then incubated at 37 °C for 10 minutes and subsequently rotated for at least 30 minutes. The purified DNA was eluted into a new tube, and its concentration and purity reassessed using NanoDrop. For applications requiring higher DNA concentrations, such as Oxford Nanopore sequencing, sample pooling and concentration via SpeedVac were recommended. Notably, to mitigate the risk of clogging in ONT flow cells, an additional step of gentle DNA shearing using a syringe was introduced, slightly compromising read length to enhance sequencing efficiency. Finally, 6.2 µl of DNA was used in the Rapid library for the Oxford Nanopore Run. Oxford Nanopore MinION sequencing was conducted over 14 runs, with each run lasting 3 days, and each run performed on its own dedicated flow cell.
